# *PerturbSci-Kinetics*: Dissecting key regulators of transcriptome kinetics through scalable single-cell RNA profiling of pooled CRISPR screens

**DOI:** 10.1101/2023.01.29.526143

**Authors:** Zihan Xu, Andras Sziraki, Jasper Lee, Wei Zhou, Junyue Cao

## Abstract

Here we described *PerturbSci-Kinetics*, a novel combinatorial indexing method for capturing three-layer single-cell readout (*i.e.,* whole transcriptomes, nascent transcriptomes, sgRNA identities) across hundreds of genetic perturbations. Through *PerturbSci-Kinetics* profiling of pooled CRISPR screens targeting a variety of biological processes, we were able to decipher the complexity of RNA regulations at multiple levels (*e.g.,* synthesis, processing, degradation), and revealed key regulators involved in miRNA and mitochondrial RNA processing pathways. Our technique opens the possibility of systematically decoding the genome-wide regulatory network underlying RNA temporal dynamics at scale and cost-effectively.

## Main

Cellular functions are determined by the expression of millions of RNA molecules, which are tightly regulated across several critical steps, such as RNA synthesis, splicing, and degradation. However, our knowledge regarding how critical molecular regulators affect genome-wide RNA kinetics remains limited. For example, recent studies have combined single-cell transcriptome analysis with pooled CRISPR screens to gain insights into the gene regulation mechanisms^1–8^. Yet, these methods provide only a snapshot of gene expression programs and fail to capture the complexity of RNA dynamics (*e.g.,* synthesis, splicing, and degradation). Although RNA metabolic labeling coupled with single-cell sequencing can reveal time-resolved transcriptomic dynamics, there is still a need for scalable tools that can efficiently characterize the impact of genetic perturbations on RNA dynamics in a high-throughput manner^9^. To resolve this challenge, we developed *PerturbSci-Kinetics*, which integrates CRISPR-based pooled genetic screen, highly scalable single-cell RNA-seq by combinatorial indexing, and metabolic labeling to recover single-cell transcriptome dynamics across hundreds of genetic perturbations.

The new method features a novel combinatorial indexing strategy (referred to as ‘*PerturbSci’)* for targeted enrichment and amplification of sgRNA transcripts that carries the same cellular barcode with the single-cell whole transcriptome (**Fig 1a**). In brief, we adopted the modified CROP-seq vector system^5^ and developed a strategy for direct capture of sgRNA sequences^6, 7^, through reverse transcription using a sgRNA-specific primer followed by targeted enrichment of sgRNA sequences via PCR (**Extended Data Fig 1a-b**). A comparison of chemistries between *PerturbSci* and other similar approaches (e.g., CROP-seq^5^, and Direct-capture Perturb-seq^7^) is included in **Extended Data Fig 1c**. With extensive optimizations on the primer design and reaction conditions (**Extended Data Fig 2**), *PerturbSci* achieves a high capture rate of sgRNA (*i.e.,* up to 99.7%), comparable to previous approaches for single-cell profiling of pooled CRISPR screens^1–7^. Furthermore, built on an extensively improved single-cell RNA-seq by three-level combinatorial indexing (*i.e.,* EasySci-RNA^10^), *PerturbSci* substantially reduced the library preparation costs for single-cell RNA profiling of pooled CRISPR screens (**Fig 1b, Supplementary file 3**). In addition, to maximize the gene knockdown efficacy, we used *dCas9-KRAB-MeCP2*^11^, a highly potent dual-repressor dCas9 that outperforms conventional dCas9 repressors.

**Fig. 1.**
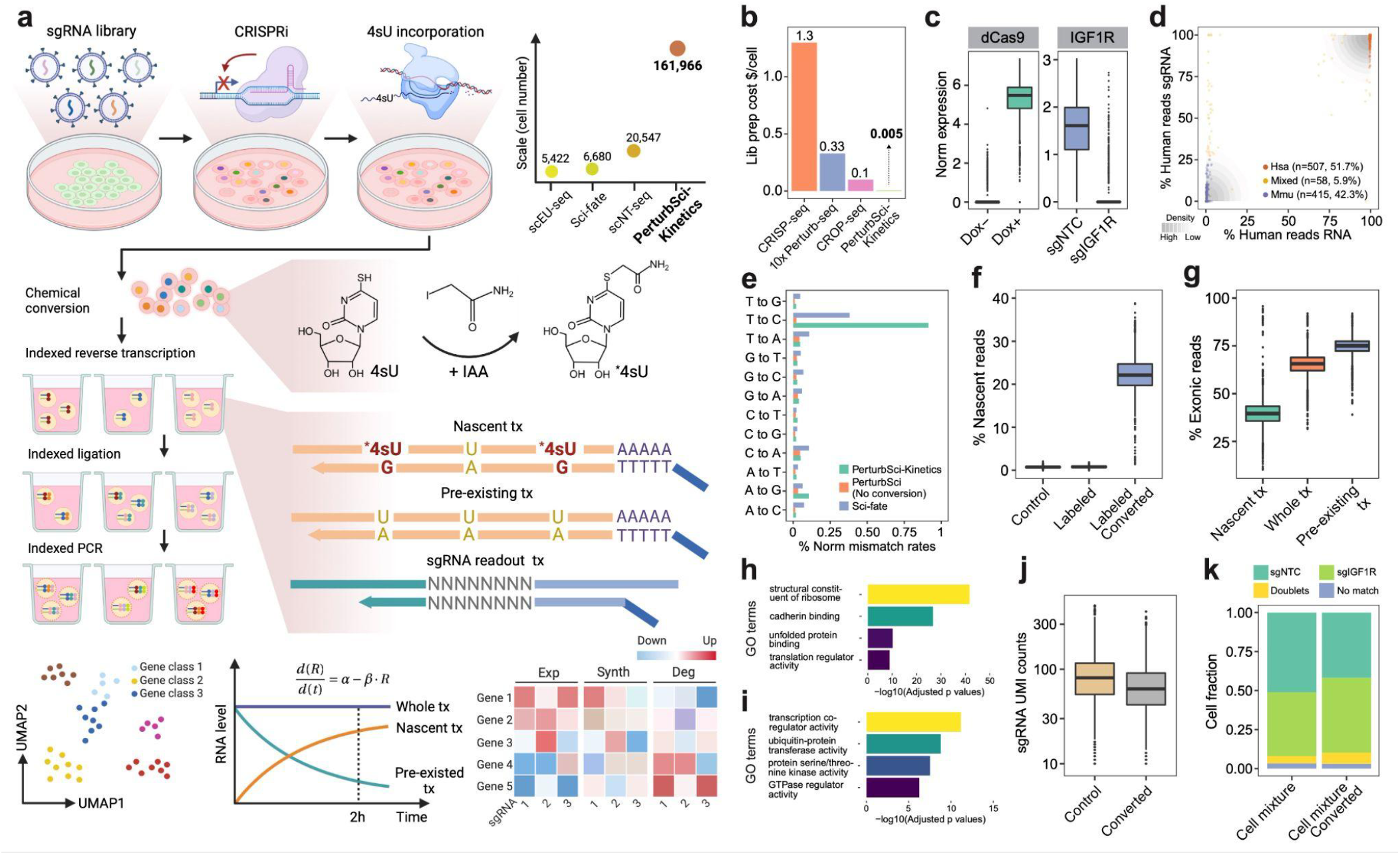
*PerturbSci-Kinetics* enables joint profiling of transcriptome dynamics and high-throughput gene perturbations by pooled CRISPR screens. **a.** Scheme of the experimental and computational strategy for *PerturbSci-Kinetics*. The dot plot on the upper right shows the number of cells profiled in this study for comparison with the published single-cell metabolic profiling datasets^19, 22, 23^. Scale, the highest number of cells profiled in a single experiment of each technique. IAA, iodoacetamide. *4sU, chemically modified 4sU. R, steady-state RNA level. α, RNA synthesis rate. β, RNA degradation rate. Exp, steady-state expression. Synth, synthesis rate. Deg, degradation rate. **b**. Bar plot showing the estimated library preparation cost for *PeturbSci-Kinetics* and other published techniques^25, 26^ for single-cell transcriptome analysis coupled with CRISPR screens. **c.** The left box plot shows the normalized expression of dCas9-KRAB-MeCP2 in untreated or Dox-induced HEK293-idCas9 cells. The right box plot shows the normalized expression of IGF1R in Dox-induced HEK293-idCas9 cells transduced with sgNTC or sgIGF1R. Gene counts of each single cell were normalized by the total gene count, multiplied by 1e4 and then log-transformed. **d.** An equal number of induced HEK293-idCas9-sgIGF1R cells and 3T3-CRISPRi-sgFto cells were mixed after cell collection and were profiled using *PerturbSci*. The scatter plot displays the percentage of single-cell transcriptome reads that were mapped to the human genome on the x-axis, and the percentage of single-cell sgRNA reads that were mapped to the human cell-specific sgRNA on the y-axis. The shading represents the density of dots. **e.** Bar plot showing the normalized percentage of all possible single base mismatches in reads from sci-fate (blue), and *PerturbSci-Kinetics* on chemically converted (green) or unconverted cells (orange). Normalized mismatch rates, the percentage of each type of mismatch in all sequencing bases. **f**. Box plot showing the fraction of recovered nascent reads in single-cell transcriptomes across conditions: no 4sU labeling + no chemical conversion, 4sU labeling + no chemical conversion, and 4sU labeling + chemical conversion. **g.** Box plot showing the ratio of reads mapped to exonic regions of the genome in nascent reads, pre-existing reads, and reads of the whole transcriptomes across single cells. **h-i.** Bar plots showing the significantly enriched Gene Ontology (GO) terms in the list of genes with low (h) or high (i) nascent reads fractions (**Methods**). **j.** Box plot showing the number of unique sgRNA transcripts detected per cell in cells with or without the chemical conversion. **k.** We performed *PerturbSci-Kinetics* experiment using converted/unconverted HEK293-idCas9 cells transduced with sgNTC/sgIGF1R. Stacked bar plot showing the fraction of converted/unconverted cells identified as sgNTC/sgIGF1R singlets, doublets, and cells with no sgRNA detected.

By integrating *PerturbSci* with 4-thiouridine (4sU) labeling method^12^, *PerturbSci-Kinetics* enables the capture of single-cell time-resolved nascent transcriptomes. Specifically, following 4sU labeling and *in-situ* thiol (SH)-linked alkylation reaction^12–18^ (referred to as ‘chemical conversion’), the nascent transcriptome and the whole transcriptome from the same cell can be distinguished by T to C conversions in reads mapping to mRNAs^19^. The kinetic rates of mRNA (*e.g.,* synthesis and degradation) were then inferred for each genetic perturbation (**Fig 1a**, **Methods**). Notebly, *PerturbSci-Kinetics* exhibits orders of magnitude higher throughput than previous metabolic labeling-coupled single-cell RNA-seq approaches (*e.g.*, scEU-seq, sci-fate, scNT-seq)^19–23^(**Fig 1a**). We extensively optimized the cell fixation condition to reduce the cell loss during cell permeabilization and chemical conversion (**Extended Data Fig 3a-e**). We also optimized the computational pipeline for nascent reads calling^19^, enabling the identification of single cell nascent transcriptomes with higher accuracy (**Extended Data Fig 4**). Statistical benchmarking with other published datasets^19^ further validated the data quality (**Extended Data Fig 3h-i**).

As a proof of concept, we generated a human HEK293 cell line with the inducible expression of dCas9-KRAB-MeCP2^11^ (HEK293-idCas9), then transduced cells with a non-target control (NTC) sgRNA (sgNTC) or a sgRNA targeting *IGF1R* (sgIGF1R). The induction of dCas9 expression after Dox treatment and the high knockdown efficiency on the target gene were validated by single-cell RNA-seq, RT-qPCR and flow cytometry (**Fig 1c, Extended Data Fig 5**). By profiling a 1:1 mixed cell pool consisting of human HEK293-idCas9-sgIGF1R cells and mouse 3T3-CRISPRi-sgFto cells, we recovered 94.1% of singlets with both species-specific whole transcriptomes and sgRNAs (**Fig 1d**, **Extended Data Fig 2l**), validating the purity of the single-cell transcriptomes and sgRNAs co-captured by *PerturbSci*.

We next sought to validate the ability of *PerturbSci-Kinetics* to capture three-layer readout (*i.e.,* whole transcriptomes, nascent transcriptomes, sgRNA identities) at the single-cell level. Following 4sU labeling (200uM for two hours), we mixed HEK293-idCas9-sgNTC cells and sgIGF1R cells for fixation and chemical conversion. A significant enrichment of T to C mismatches in mapped reads of the chemical conversion group was observed, similar to our previous study^19^ (**Fig 1e**). A median of 22.1% of newly synthesized reads was recovered in labeled and chemically converted cells, compared to only 0.8% in the control group (**Fig 1f**). Reassuringly, the proportion of reads mapped to exonic regions was significantly lower in nascent reads compared with pre-existing reads (p-value < 1e-20, Tukey’s test after ANOVA) (**Fig 1g**, **Extended Data Fig 3g**). Indeed, genes with a higher fraction of nascent reads were significantly enriched in highly dynamic biological processes such as transcription coregulator activity (FDR = 5.7e-12) and protein kinase activity (FDR = 2.6e-08)^24^ **(Fig 1h**). By contrast, genes with a lower fraction of nascent reads were strongly enriched for processes essential for cell vitality, such as the structural constituent of ribosome (FDR = 1.5e-42), unfolded protein binding (FDR = 4.5e-11), and translation regulator activity (FDR = 8.2e-10) (**Fig 1i**). Notably, the chemical conversion step is fully compatible with sgRNA detection at single-cell resolution: we recovered sgRNAs from 97% of chemically converted cells (a median of 62 sgRNA UMIs/cell), 92.6% of which were annotated as sgRNA singlets (**Fig 1j-k)**. These analyses demonstrate the capacity of *PerturbSci-Kinetics* to profile both transcriptome dynamics and the associated perturbation identities at the single-cell level.

To dissect the impact of key genetic regulators on transcriptome kinetics, we performed a *PerturbSci-Kinetics* screen on HEK293-idCas9 cells transduced with a library of 699 sgRNAs, which contains 15 NTC sgRNAs and sgRNAs targeting 228 genes involved in a variety of biological processes such as mRNA transcription, processing, degradation (**Fig 2a, Supplementary Table 1**). The cloning and lentiviral packaging were carried out in a pooled fashion, similar to the previous report^27^ (**Methods**). We then infected the HEK293-idCas9 cell line with the sgRNA lentiviral library at a low multiplicity of infection (MOI) (2 repeats at MOI = 0.1 and 2 repeats at MOI = 0.2) to ensure that most cells received only one sgRNA. After a 5-day puromycin selection to remove untransduced cells, we harvested a fraction of cells for bulk library preparation (‘day 0’ samples). The rest of cells were treated with 1ug/ml Doxycycline (Dox) to induce the dCas9-KRAB-MeCP2 expression for an additional seven days. At the end of the screen, we introduced 4sU labeling (200uM for two hours) and harvested samples for both bulk screen and single-cell *PerturbSci-Kinetics* library preparation (‘day 7’ samples). The time window for the screening period was chosen to ensure sufficient knockdown efficiency and establish new transcriptomic steady states^28^, as well as to minimize the effect of population dropout^8^ (**Methods**).

**Fig 2.**
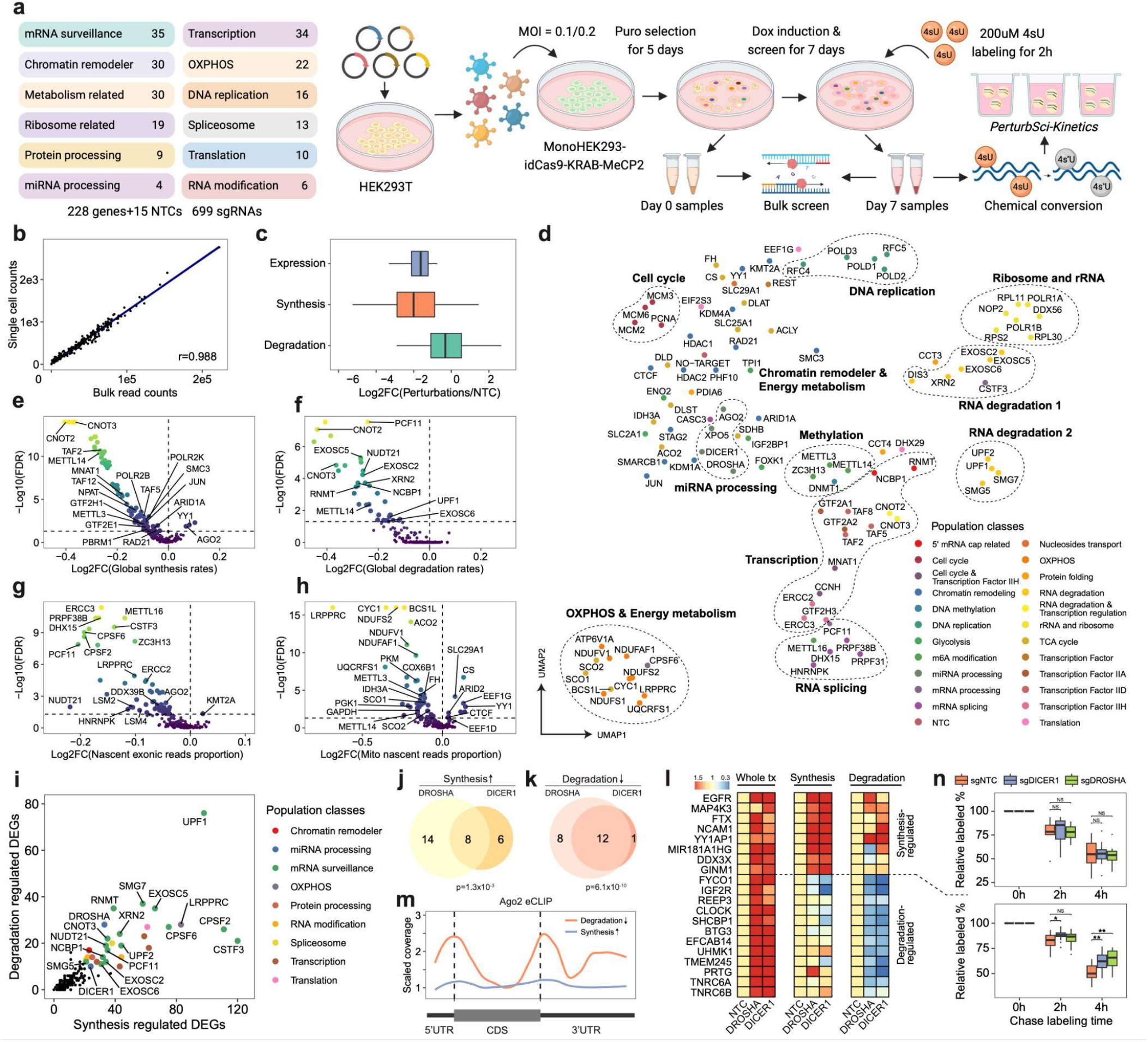
Characterizing the impact of genetic perturbations on gene-specific transcriptional and degradation dynamics with *PerturbSci-Kinetics*. **a.** Scheme of the experimental design of the *PerturbSci-Kinetics* screen. The main steps are described in the text. **b.** The scatter plot shows the correlation between perturbation-associated cell count from *PerturbSci-Kinetics* and sgRNA read counts from bulk screen libraries. **c.** Box plot showing the log2 transformed fold changes of gene expression, synthesis rates, and degradation rates of sgRNA-targeted genes in perturbed cells expressing the corresponding sgRNA, compared to NTC. **d.** UMAP visualization of perturbed pseudobulk whole transcriptomes profiled by *PerturbSci-Kinetics*. We aggregated single-cell transcriptomes in each perturbation, followed by dimension reduction using PCA and visualization using UMAP. Population classes, the functional categories of genes targeted in different perturbations. **e-h.** Scatter plots showing the extent and the significance of changes on the distributions of global synthesis (e), degradation (f), proportions of exonic reads in the nascent transcriptome (g), and proportions of mitochondrial nascent reads (h) upon perturbations compared to NTC cells. The fold changes were calculated by dividing the median values of each perturbation with that of NTC cells and were log2 transformed. **i.** Scatter plot showing the number of synthesis/degradation-regulated DEGs from different perturbations. nDEGs: the number of DEGs. **j-k.** Venn diagrams showing the number of merged DEGs with significantly enhanced synthesis (j) or impaired degradation (k) between *DROSHA* and *DICER1*. Based on statistical test results, merged DEGs of *DROSHA* and *DICER1* perturbations were classified into synthesis-regulated genes (*i.e.,* the upregulation of these genes was mainly driven by increased synthesis rates) and degradation-regulated genes (*i.e.,* the upregulation of these genes were mainly driven by reduced degradation rates). Merged DEGs with p-value <= 0.05 on synthesis increase/degradation decrease in at least one perturbation were included in the diagram, in which genes with p-value < 0.1 on synthesis increase/degradation decrease in both perturbations were regarded as shared hits between two perturbations. **l.** Heatmaps showing the steady-state expression, synthesis and degradation rate changes of genes sharing the same regulatory mechanism upon *DROSHA* and *DICER1* knockdown as shown in j-k. Tiles of each row were colored by fold changes of values of perturbations relative to NTC. **m.** Line plot showing the *Ago2* binding patterns on the transcript regions of protein-coding genes in Figure 2n and 2o. The transcript regions of genes were assembled by merging all exons, and were divided into 5’UTR, coding sequence (CDS), and 3’UTR based on coordinates of the 5’ most start codon and the 3’ most stop codon. Single-base coverage of *Ago2* eCLIP on each gene was calculated, binned, and scaled to 0-1. After merging and averaging scaled binned coverage of genes in the same group together, the lowest coverage value in the CDS was used to scale the averaged merged coverage again to visualize the *Ago2*/RISC binding pattern. **n.** Boxplots showing the relative proportion of labeled mRNA in a chase labeling experiment. HEK293-idCas9-sgNTC, sgDROSHA, and sgDICER1 cells were labeled with 100uM 4sU for 18 hours, followed by a 10mM uridine chase for 0h, 2h, and 4h. This tracked the degradation of labeled RNA in genes regulated by transcription (top) or degradation (bottom) upon *DROSHA* and *DICER1* knockdown (**Methods**). Student t-tests were performed between knockdown groups and the NTC group. **, p-value < 1e-2; *, p-value < 0.05; NS, no statistical significance.

As anticipated, activated CRISPRi significantly altered sgRNAs abundance in the pool, which was consistent between replicates and concordant with the previous study^29^ (**Extended Data Fig 6a-b**, **Supplementary Table 2, 3**). For example, sgRNAs-targeting genes involved in essential biological functions, such as DNA replication, ribosome assembly, and rRNA processing, were strongly depleted after the screen (**Extended Data Fig 6c**). Reassuringly, the sgRNA abundance recovered by *PerturbSci-Kinetics* significantly correlated with the bulk screen libraries (Pearson correlation r = 0.988, p-value < 2.2e-16) **(Fig 2b)**.

After filtering out low-quality cells, we recovered 161,966 labeled cells, 88% of which had matched sgRNAs (>= 10 sgRNA UMIs). A majority (78%) of these matched cells were annotated as sgRNA singlets (**Extended Data Fig 7a**). Despite the shallow sequencing (duplication rate of 17.9%), we obtained a median of 2,155 UMIs per cell. Most (698 out of 699) sgRNAs were recovered, with a median of 28 sgRNA UMIs detected per cell. sgRNAs with low knockdown efficiencies (<= 40% expression reduction of target genes compared with NTC) were further filtered out (**Extended Data** Fig 7b-e). Of note, the knockdown efficiencies of several individual sgRNAs were further examined using RT-qPCR, which showed high consistency with the pooled screen (**Extended Data** Fig 7f). Finally, 98,315 cells were retained for downstream analysis, corresponding to a median of 484 cells per gene perturbation with a median of 67.7% knockdown efficiency of target genes (**Fig 2c**).

Taking advantage of the ability of *PerturbSci-Kinetics* to capture multiple layers of information, we quantified gene-specific synthesis and degradation rates in each perturbation based on an ordinary differential equation^30^ (**Methods**). As CRISPRi is known to function through transcriptional repression^31, 32^, we first examined the kinetic changes of sgRNA target genes upon perturbation. Indeed, these genes exhibited strongly reduced synthesis rates while their degradation rates were only mildly affected (**Fig 2c**). As a further validation of the technique, we observed significantly higher correlations among sgRNAs targeting the same genes in multiple layers (*e.g.,* whole/nascent transcriptome, synthesis/degradation rates, **Extended Data Fig 8a**).

To further understand the impact of gene perturbations, we aggregated whole transcriptomes of each gene perturbation, followed by PCA for dimension reduction and UMAP visualization^33^(**Methods**). Indeed, perturbations targeting paralogous genes (*e.g., EXOSC5* and *EXOSC6*; *CNOT2* and *CNOT3*) or related biological processes (*e.g.,* RNA degradation, RNA splicing, oxidative phosphorylation (OXPHOS) and energy metabolism) were readily clustered together in the low dimension space (**Fig 2d**). Reassuringly, UMAP embedding on gene-specific synthesis/degradation rates also grouped perturbations by their functions (**Extended Data Fig 8b-c**).

We then investigated the impact of genetic perturbations on global transcriptome dynamics (*i.e.,* synthesis, splicing, and degradation) (**Fig 2e-g, Extended Data Fig 9, Methods, Supplementary Table 4, 5**). As expected, the knockdown of genes involved in transcription initiation (*e.g., GTF2E1*, *TAF2*, *MED21,* and *MNAT1*), mRNA synthesis (*e.g., POLR2B* and *POLR2K*), and chromatin remodeling (*e.g., SMC3, RAD21, CTCF, ARID1A*) significantly downregulated the global synthesis rates but not the degradation rates (**Fig 2e-f**). In contrast, perturbations targeting components of critical biological processes such as DNA replication (*e.g.*, *POLA2*, *POLD1*), ribosome synthesis and rRNA processing (*e.g.*, *POLR1A, POLR1B, RPL11, RPS15A*), mRNA and protein processing (*e.g.*, *CNOT2, CNOT3, CCT3, CCT4*) substantially reduced both RNA synthesis and degradation globally, indicating a compensatory mechanism for maintaining overall transcriptome homeostasis **(Fig 2e-f, Extended Data Fig 9a-b)**. Furthermore, we observed significantly reduced exonic reads fractions in nascent transcriptomes, an indicator of dysregulated splicing dynamics, following perturbations of genes involved in the main steps of RNA processing, including 5’ capping (*e.g., NCBP1*), RNA splicing (*e.g., LSM2, LSM4, PRPF38B, HNRNPK*), and 3’ cleavage/polyadenylation (*e.g., CPSF2, CPSF6, NUDT21, CSTF3*) **(Fig 2g, Supplementary Table 6)**. In addition, knocking down genes related to OXPHOS & energy metabolism (*e.g., GAPDH, NDUFS2, ACO2*) also significantly reduced the exonic reads ratio in nascent reads **(Fig 2g, Extended Data Fig 9c)**, potentially because the mRNA processing is highly energy-dependent^34–36^.

Interestingly, we found the knockdown of *AGO2*, a well-established post-transcriptional silencer^37^, resulted in an significant increase of the global synthesis (**Fig 2e)**, suggesting its direct involvement in global transcription regulation. To validate this finding, we first examined Ago2 ChIP-seq data and bulk RNA-seq data upon shRNA-mediated silencing of AGO2 from ENCODE^38, 39^. We observed a group of genes with strong Ago2 binding in close proximity to their transcription start sites (TSS) (**Extended Data Fig 10a**), and these genes were significantly upregulated upon AGO2 silencing, consistent with the transcription repressor role of AGO2 identified in our study (**Extended Data Fig 10b**). Moreover, we observed the enrichment of Ago2 binding right downstream of TSS (**Extended Data Fig 10a**), reflecting a potential role of Ago2 in regulating transcriptional pausing. To infer the gene-specific transcriptional pausing state, we analyzed another published GRO-seq dataset and quantified gene-specific Pausing Index (PI)^40, 41^ (**Methods**). Remarkably, a strong positive association between AGO2 TSS binding and PI across genes was observed (**Extended Data Fig 10c-d**). We sought to further validate this mechanism by tracking on-going transcription. Following the construction of HEK293-idCas9-sgAGO2 cell line and 7-day Dox induction, we performed 20min short-term 4sU labeling and full-coverage SLAM-seq^42^ (**Methods**). AGO2 knockdown significantly upregulated 78 highly-paused genes, and their nascent RNA exhibited increased 3’ end coverage upon AGO2 knockdown (**Extended Data Fig 10e-f**), indicating more efficient transcriptions. Together, our integrated analyses strongly supported the non-canonical role of AGO2 in transcription identified by *PerturbSci-Kinetics*.

We next sought to investigate regulators of mitochondrial RNA dynamics by quantifying the fraction of nascent read counts for mitochondrial genes (referred to as “mitochondrial transcriptome turnover”) (**Methods**). Notably, we observed a significantly reduced turnover of mitochondrial transcriptome following the perturbation of multiple metabolism-related genes (*e.g., GAPDH*, *FH*, *PKM* involved in glycolysis, *ACO2,* and *IDH3A* involved in the TCA cycle, *NDUFS2* and *COX6B1* involved in oxidative phosphorylation) **(Fig 2h, Extended Data Fig 9d, Supplementary Table 7)**. Furthermore, the knockdown of *LRPPRC* introduced the most substantial defect in the mitochondrial turnover and expression levels of all mitochondrial protein-coding genes **(Fig 2h, Extended Data Fig 11a**). By examining mitochondrial gene-specific kinetics, we identified 5 of 13 mitochondrial protein-coding genes, including *MT-CO1, MT-ATP8, MT-ND4, MT-CYB, and MT-ATP6*, to be significantly regulated by both decreased synthesis and increased degradation (**Extended Data Fig 11a, Supplementary Table 8**).

To further validate our findings, we examined the bulk RNA-seq and the RNase footprinting datasets from Lrpprc knockout mice^43^. While steady-state mitochondrial gene down-regulation showed great consistency across species, we further observed the strong relationship between mitochondrial mRNA stability and RNA secondary structure, potentially indicating the RNA stabilizing mechanism of LRPPRC (**Extended Data Fig 11b**). Moreover, the strong activation of integrated stress response (ISR) phenotype in LRPPRC knockdown cells enforced the essentiality of LRPPRC in maintaining mitochondrial mRNA homeostasis (**Extended Data Fig 11c-f**). By profiling HEK293-idCas9-sgLRPPRC cells following Dox induction and 4sU labeling, we further validated the robustness of kinetics changes of mitochondrial mRNA upon LRPPRC perturbation (**Extended Data Fig 10g-i**). It is worth noting that impaired mitochondrial gene expression and the functional defect of mitochondria in brown adipocytes specific LRPPRC knockout mice were reported in a recent study^44^. Overall, *PerturbSci-Kinetics* identified LRPPRC as a key regulator of mitochondrial RNA dynamics.

Extending the above analysis, we identified differentially-expressed genes (DEGs) across perturbations (**Supplementary Table 9**) and examined their dynamic rate changes (**Supplementary Table 10**) (**Methods**). Out of 14,618 perturbation-DEGs pairs, 22.9% exhibited significant dynamics rate changes: 15.1% showed synthesis rate changes only, 3.6% showed degradation rate changes only, and 4.2% showed changes on both, suggesting that RNA dynamics control is highly involved in gene regulation^45^. As expected, degradation-regulated DEGs were highly associated with perturbations on mRNA surveillance/processing genes (*e.g., UPF1, UPF2, SMG5, SMG7* in nonsense-mediated mRNA decay pathway; *EXOSC2, EXOSC5, EXOSC6* in RNA exosome; *CSTF3, CPSF2, CPSF6, NUDT21, XRN2* for 3’ polyadenylation; *RNMT, NCBP1* related to 5’ RNA capping) **(Fig 2i)**. For instance, our study identified a set of significantly overlapped DEGs upon knockdown of *DROSHA* and *DICER1*^46, 47^, two crucial regulators in the miRNA biogenesis pathway^48^. By concurrently profiling gene-specific expression and dynamics, we discovered that these differentially expressed genes were regulated through distinct mechanisms: some genes were regulated through decreased degradation (*e.g.,* miRNA-mediated silencing complex (RISC) components: TNRC6A and TNRC6B), while others are through increased transcription (*e.g.,* miRNA host genes: MIR181A1HG, FTX; genes involved in miRNA biogenesis: DDX3X) (**Fig 2j-l, Extended Data Fig 12a-c**). To reveal the direct involvement of miRNA-mediated RNA destabilization in regulating these degradation-regulated DEGs, we visualized the mRNA transcripts binding patterns of *Ago2*, a core component of RISC for targeted mRNA binding and degradation^49^. As expected, *Ago2* binding was strongly enriched in the 5’ and 3’ untranslated region (UTR) of transcripts from genes with reduced degradation but not in transcripts from genes with upregulated synthesis **(Fig 2m)**, consistent with prior studies that miRNA induces targeted RNA degradation and translation repression mainly through binding to UTRs^46, 50^.

As validation, we profiled HEK293-idCas9 cells transduced with sgRNA targeting each of the four miRNA biogenesis pathway genes (*XPO5, AGO2, DROSHA, DICER1*) after Dox induction and 4sU labeling. Both single-cell transcriptome UMAP embedding and RNA dynamics changes recapitulated our findings in the pooled screen (**Extended Data Fig 12d-f).** In addition, by using 4sU chase labeling and 3’end bulk SLAM-seq^42^, we tracked RNA degradation dynamics in *DROSHA* and *DICER1* knockdown cells and confirmed the enhanced stability of mRNA from degradation-regulated genes (*e.g., SHCBP1, PRTG*), compared to transcription-related genes (*e.g., FTX, YY1AP1*) (**Fig 2l, n, Extended Data Fig 12g**). These analyses further demonstrate the unique capacity of *PerturbSci-Kinetics* of deciphering the regulatory mechanisms (degradation vs. transcription) involved in gene expression changes.

Finally, to explore cell-state dependent RNA dynamics changes upon genetic perturbations at the sub-population level, we investigated the effects of perturbations on RNA dynamics throughout the cell cycle using our validation dataset, in which we profiled more cells with deeper sequencing. Using a combination of cell cycle-related genes^51^ and cell cycle-related transcription factors^19^ for dimension reduction and clustering, we separated cells into five clusters representing different cell cycle stages (**Extended Data Fig 13a-c**). Gene-specific synthesis and degradation rates in the physiological condition were inferred across cell cycle stages in NTC cells, and four gene clusters with different cell cycle-dependent synthesis dynamics were identified^52^ (**Extended Data Fig 13d**). Among clusters, only genes in cluster 1 exhibited a cell cycle stage-specific expression pattern. While their synthesis and degradation rates both increased with the cell cycle progression, their synthesis rates outpaced the degradation rates, driving the increase of their steady-state mRNA levels from S to G2M stage (**Extended Data Fig 13d**). GO term enrichment analysis confirmed the strong association between functions of these genes and cell division^53^ (**Extended Data Fig 13e**).

However, in DROSHA and DICER1 knockdown cells, while the steady-state expression pattern of cluster 1 genes resembled NTC along the cell cycle, degradation rates of these genes failed to respond, especially in the early phase of the cell cycle (**Extended Data Fig 13f**). Surprisingly, it was fully compensated by the reduced synthesis rates (**Extended Data Fig 13f**), suggesting the existence of synthesis-degradation feedback loops for gene regulation. As a result of this buffering effect, both DROSHA and DICER1 knockdown did not substantially affect the cell-cycle progression (**Extended Data Fig 13b**). In comparison, the knockdown of LRPPRC did not impair the RNA degradation dynamics of these genes throughout the cell cycle progression (**Extended Data Fig 13g**), which is consistent with our conclusion that LRPPRC primarily affects mitochondrial mRNA stability (**Extended Data Fig 10d**). Together, our findings reveal the highly coordinated RNA synthesis-degradation regulations and the existence of feedback loops between RNA synthesis and degradation throughout the cell cycle.

In summary, *PerturbSci-Kinetics* is the first method that allows for the quantitative analysis of the genome-wide mRNA kinetic rates across hundreds of genetic perturbations in a single experiment. We have provided step-by-step protocols and data processing pipelines in supplementary files (**Supplementary file 1-4**) to facilitate the broad applications of the technique. Of note, there are several potential limitations to consider: First, long-term 4sU labeling might alter cell states and hinder the identification of sgRNA sequences. We thus chose a relatively short-term (2 hours) treatment to minimize such effects. Second, RNA dynamics identified by *PerturbSci-Kinetics* may not directly indicate causality in gene regulation, partly due to the long duration of CRISPRi-based gene knockdown. This limitation could be mitigated by coupling the technique with large-scale chemical perturbations that allow short treatment and labeling time.

Despite these limitations, our results uncover the unique advantages of *PerturbSci-Kinetics* over conventional assays. Its multi-layer readout offers a comprehensive view of gene expression and RNA dynamics in response to genetic perturbations, facilitating high-throughput and parallel characterization of regulators of gene-specific transcription, splicing, and degradation. Moreover, the low-cost and the high sgRNA capture sensitivity of *PerturbSci* allows for systematic characterization of cell-type-specific gene regulatory networks in various biological contexts at an unprecedented scale and resolution both *in vitro* and *in vivo*.

## Endnotes

### Acknowledgments

We thank all members of the Cao lab for helpful discussions and feedback. We thank Dr. R. Satija (New York Genome Center) for insightful feedback related to this work. We thank the Tissue Culture facility of the University of California, Berkeley for the 3T3 cell line, and the Scott Keeney Lab at Memorial Sloan Kettering Cancer Center for the HEK293 cell line. We thank members of the Rockefeller University Flow Cytometry Resource Center and the Rockefeller University Genomics Resource Center for their extensive help with FACS sorting and sequencing experiments. We also thank members of the Information Technology and High-Performance Computing team at Rockefeller University, especially J. Banfelder and B. Jayaraman for their great support. We acknowledge that the research resulting in this publication was supported, in part, by The G. Harold and Leila Y. Mathers Charitable Foundation.

## Author contributions

J.C. and W.Z. conceptualized and supervised the project. Z.X. performed experiments, including technique development and optimization, with input from J.L.. Z.X. performed computational analyses, with input from A.S.. J.C., W.Z., and Z.X. wrote the manuscript with input and biological insight from all co-authors.

## Supplementary Figures

**Extended Data Fig. 1.**
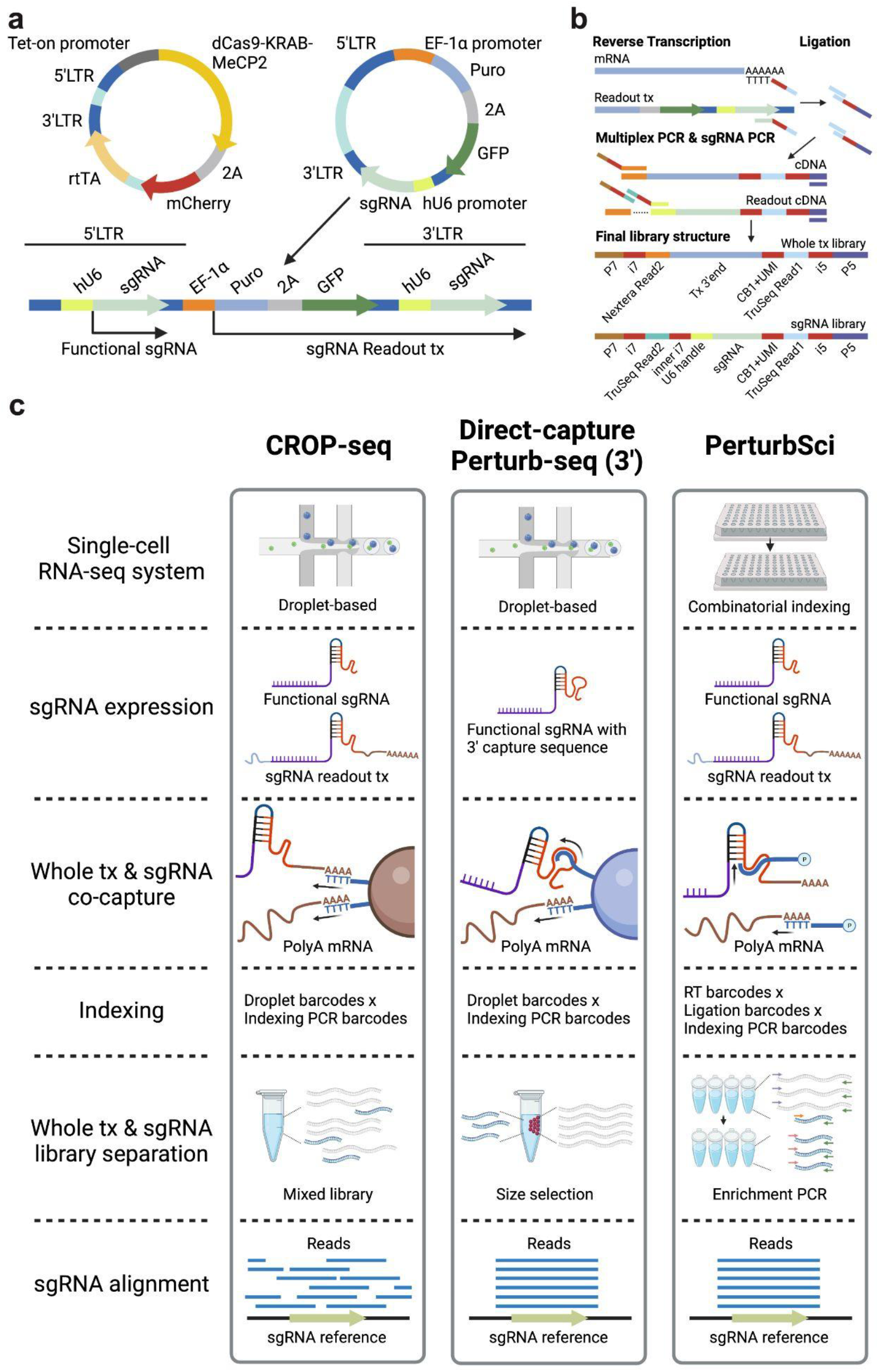
Scheme of plasmids and experiment procedures of *PerturbSci*. **a**. The vector system used in *PerturbSci* for dCas9 and sgRNA expression. The expression of the enhanced CRISPRi silencer dCas9-KRAB-MeCP2^11^ was controlled by the tetracycline responsive (Tet-on) promoter. A GFP sequence was added to the original CROP-seq-opti plasmid^6^ as an indicator of successful sgRNA transduction and for the lentivirus titer measurement. The CROP-seq vector utilizes the self-replication mechanism of lentivirus during the integration for amplifying the sgRNA expression cassette. In this lentiviral plasmid, the sgRNA expression cassette replaced the U3 region of the 3’LTR^5^. During the lentiviral integration, the shortened 5’LTR of reverse-transcribed lentiviral genomic DNA was extended by using its 3’LTR as a template, and the sgRNA expression cassette is self-replicated to the 5’LTR^54^. The self-replicated sgRNA expression cassette at 5’LTR generates functional sgRNA for guiding dCas9, and the original expression cassette at 3’LTR is transcribed as a part of the Puro-GFP fusion transcript driven by the EF-1α promoter. **b**. The library preparation scheme and the final library structures of *PerturbSci*, including a scalable combinatorial indexing strategy with direct sgRNA capture and enrichment that reduced the library preparation cost, enhanced the sensitivity of the sgRNA capture compared to the original CROP-seq^5^, and avoided the extensive barcodes swapping detected in Perturb-seq^6^. **c.** A schematic comparison of chemistries between *PerturbSci*, CROP-seq^5^, and Direct-capture Perturb-seq^7^.

**Extended Data Fig. 2.**
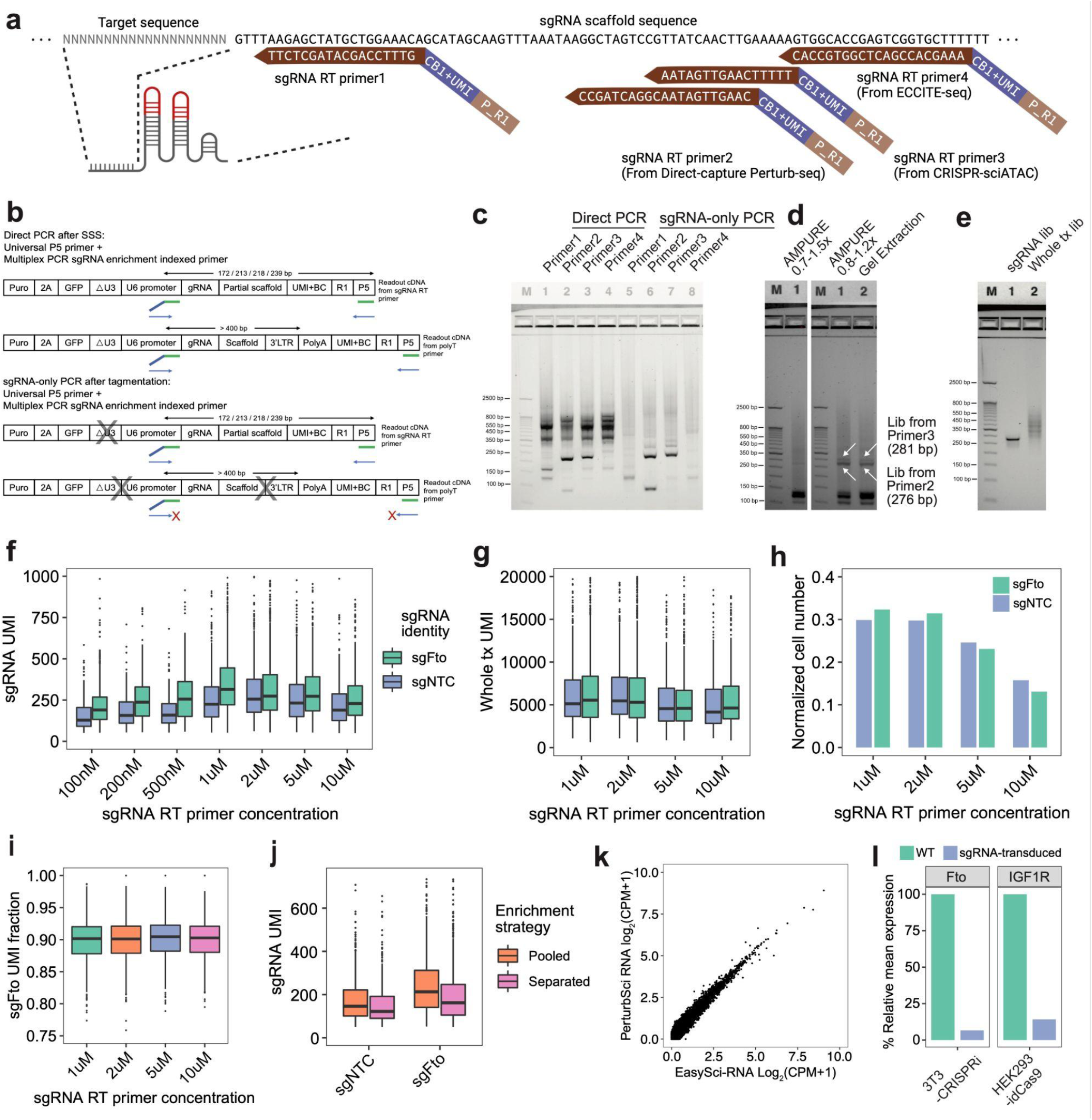
Representative optimizations on sgRNA enrichment of *PerturbSci*. **a**. Multiple RT primers targeting different sgRNA scaffold regions were mixed with polyT primers respectively and were used in our test experiment for targeted enrichment of sgRNA (RT primer 2-4 were modified from primers used in Direct-capture Perturb-seq^7^, CRISPR-sciATAC^55^, and ECCITE-seq^56^). CB, cell barcode. P_R1, partial TruSeq read1 sequence. **b-c.** A 96-well plate was divided into 4 parts and RT was performed using different combinations of sgRNA capture primers and shortdT primers. After ligation, cells were mixed and redistributed for SSS. We tested the capture efficiency of sgRNA by different RT primers in *PerturbSci* using “Direct PCR” and tested the efficiency of by-product removal by “sgRNA-only PCR” (Scheme shown in b) followed by gel electrophoresis for analyzing the PCR product (c). Crosses in b, potential Tn5 tagmentation sites. As shown in c, sgRNA primer 2 and 3 yielded strongest amplification signals following PCR, while primer1 and 4 recovered weak signals. In addition, tagmentation removed large by-products generated potentially from polyT priming (as shown in b). **d.** We tested different conditions in post-multiplex PCR purification to obtain the input for the sgRNA enrichment PCR that could maximize the recovery of the sgRNA library. Left lane: 0.7x-1.5x double-size AMPURE beads purification followed by the sgRNA enrichment PCR reaction. Middle lane: 0.8x-1.2x AMPURE beads purification followed by the sgRNA enrichment PCR reaction. Right lane: Gel extraction on multiplex PCR product within 175-275 bp range followed by the sgRNA enrichment PCR reaction. The recovered sgRNA libraries generated from gRNA primer2 and 3 were marked on the gel image. Based on the result, the sgRNA primer2 and the 0.8-1.2x AMPURE beads purification condition yielded the best performance. **e.** A representative gel image of the final libraries of *PerturbSci*, including the sgRNA library (Lane 1) and the whole transcriptome library (Lane 2). **f-i.** We tested different concentrations of sgRNA RT primers in the *PerturbSci* experiment using 3T3-L1-CRISPRi cells transduced with either sgFto and sgNTC. The box plots show the number of unique sgRNA transcripts (f) or mRNA transcripts (g) detected per cell, the cell recovery rate (h) and sgRNA capture purity (i) across different sgRNA RT primer concentrations. **j.** We performed *PerturbSci* experiment with 3T3-L1-CRISPRi cells transduced with sgFto and sgNTC. After multiplex PCR, PCR product was purified and amplified for sgRNA library enrichment in individual PCR wells or in a pooled manner. The box plot showing the number of unique sgRNA transcripts detected per cell across the two conditions. **k.** Scatter plot showing the correlation between log2-transformed counts per million (CPM) profiled by *PerturbSci* and *EasySci*^10^ in the mouse 3T3-L1-CRISPRi cell line. **l.** Barplots showing strong gene expression knockdown in both mouse 3T3-CRISPRi-sgFto cells and human HEK293-idCas9-sgIGF1R cells that we computationally assigned in the species-mixing experiment (Fig 1d), indicating the accuracy of single-cell sgRNA assignment and the compatibility of *PerturbSci* with diverse dCas9 effectors.

**Extended Data Fig. 3.**
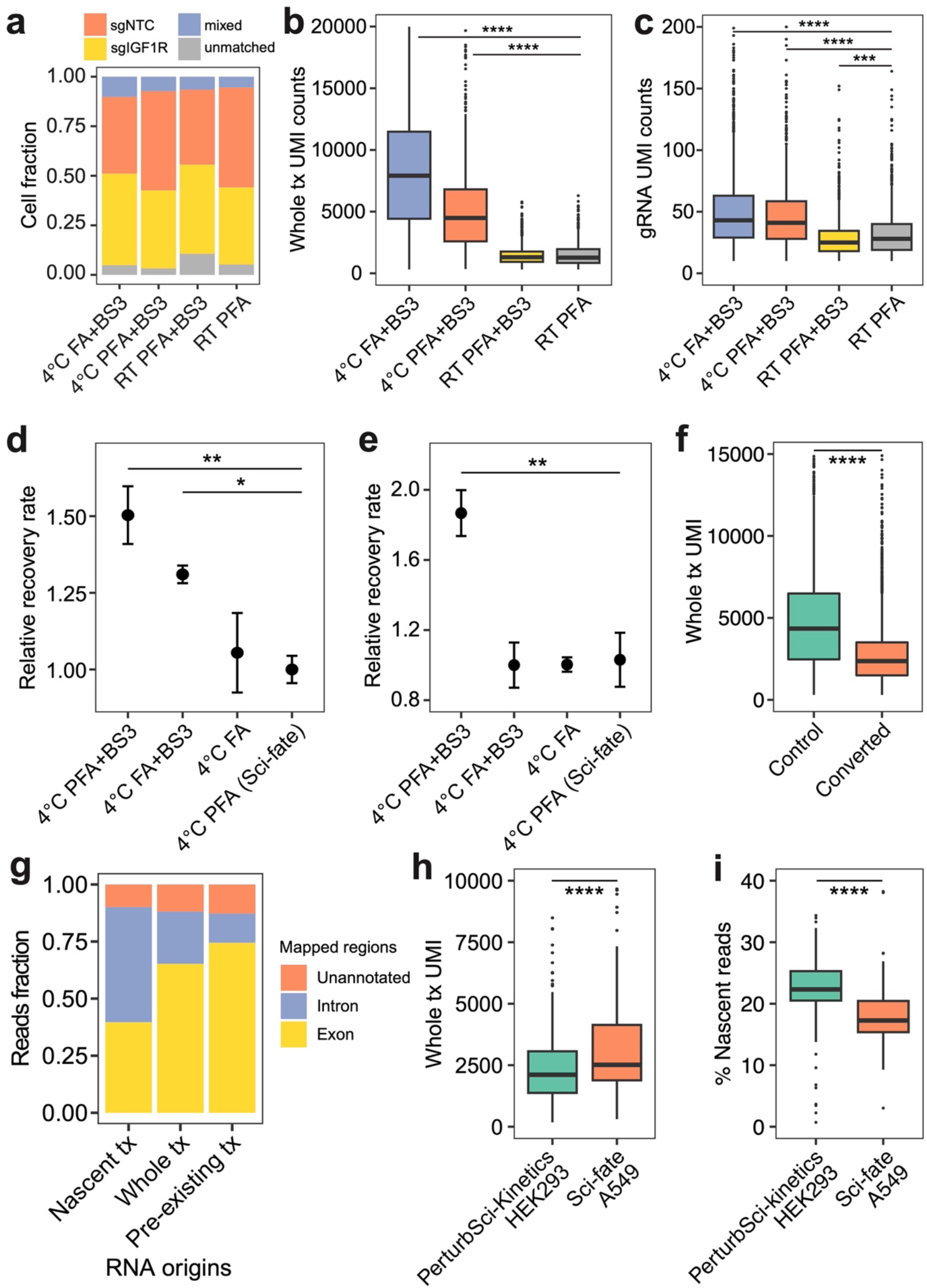
Representative optimizations on fixation conditions of *PerturbSci-Kinetics* and statistical benchmarking. We aimed to search for an optimal fixation condition that can i) minimize the cell loss during the fixation and chemical conversion, ii) reduce the RNA cross-contamination, iii) be compatible with *in-situ* combinatorial indexing of cellular transcriptomes. **a-c.** We tested different cell fixation conditions on HEK293-idCas9 cells followed by *PerturbSci* profiling and quantified the fraction of cells that were assigned to different groups (a), the number of unique sgRNA (b) and mRNAs (c) detected per cell. PFA fixation conditions at the room temperature (RT) were too strong to recover sufficient signals. FA fixation at 4°C yielded higher total UMI counts but showed stronger cross-contamination, indicating that under 4°C it was a milder fixative compared to 4% PFA. **d-e.** Dot plots showing the relative recovery rate (with standard error of the mean) of HEK293-idCas9 cells in different fixation conditions (n = 4) following HCl permeabilization (d) and chemical conversion (e). All values were normalized by the standard condition used in sci-fate (PFA fixation)^19^. **f.** Box plot showing the number of unique transcripts detected per cell with or without chemical conversion. Fixation conditions included in the plots: 4°C PFA+BS3: cells were fixed with 4% PFA in PBS for 15 minutes, and were further fixed by 2mM BS3 during and after Triton-X100 permeabilization (**Methods**). 4°C FA+BS3: cells were fixed with 1% Formaldehyde (FA) in PBS for 10 minutes, and were further fixed by 2mM BS3 during and after Triton-X100 permeabilization. 4°C FA: cells were only fixed once with 1% Formaldehyde (FA) in PBS for 10 minutes. 4°C PFA: cells were only fixed once with 4% PFA in PBS for 15 minutes as sci-fate^19^. **g.** Mapping statistics of reads of different origins from *PerturbSci-Kinetics*. Exon, reads were mapped to exonic regions of the genome. Intron, reads were mapped to intronic regions of the genome. Unannotated, reads were mapped to the genomic regions without annotation. For nascent RNA, we found an average of 40% reads mapped to exonic regions and 50% reads mapping to intronic regions. Around 0.9% of nascent reads were mapped to the antisense transcripts. We observed around 10% of the nascent reads that did not overlap with any annotations in a strand-specific manner, suggesting that they may represent unannotated transcripts from intergenic regions or antisense transcripts. Notably, the proportion of unannotated reads we observed is consistent with other sci-RNA-seq datasets^19, 57^, indicating that it is unlikely to be a technique-specific artifact. **h-i.** Downsampling comparison between sci-fate and *PerturbSci-Kinetics*. A subset of raw reads in sci-fate A549 dataset^19^ were randomly selected to generate a downsampled dataset with the same single-cell raw reads number distribution with *PerturbSci-Kinetics*, and both datasets were processed using the same pipeline. The single-cell whole transcriptome UMI counts (h) and the nascent reads percentages (i) between two datasets were compared. One-way ANOVA on all groups followed by Dunnett’s tests between control and other groups were performed in b-e. Wilcox tests were performed in f, h-i. ****, p-value < 1e-4; ***, p-value < 1e-3; **, p-value < 1e-2; *, p-value < 0.05. Comparisons with no statistical significance were not marked.

**Extended Data Fig. 4.**
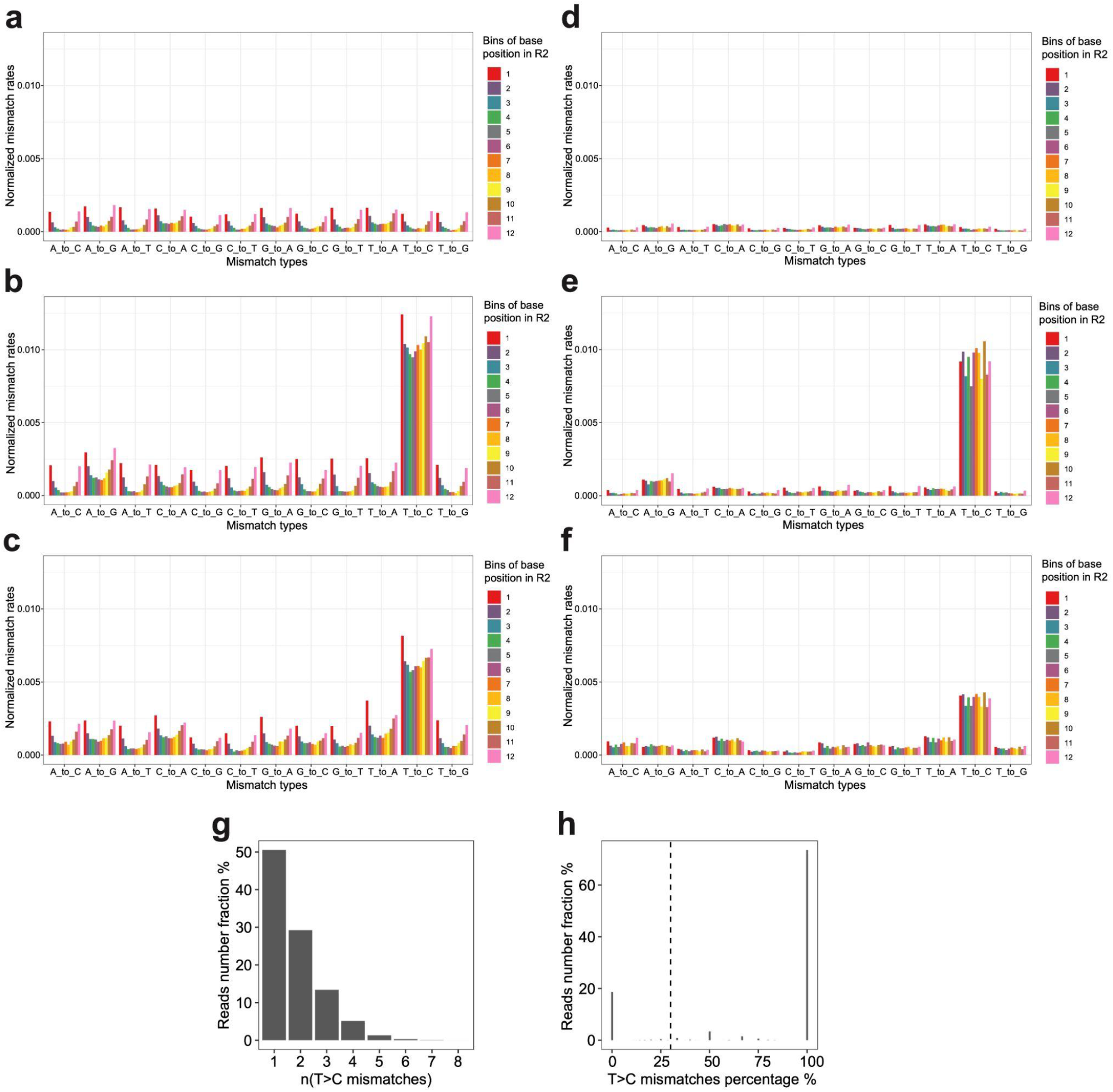
Optimization of the computational pipeline for nascent reads calling. **a-c.** Bar plots showing the normalized mismatch rates of all 12 mismatch types detected in unconverted cells (a), converted cells (b), and the original sci-fate A549 dataset^19^ (c) at different positions of the reads using the original sci-fate mutation calling pipeline^19^. **d-f.** Bar plots showing the normalized mismatch rates of all 12 mismatch types detected in unconverted cells (d), converted cells (e), and the original sci-fate A549 dataset^19^ (f) at different positions of the reads using the updated mutation calling pipeline. Considering the different sequencing lengths between the present dataset and sci-fate, the Read2 from sci-fate were trimmed to the same length as the present dataset before processing. Compared to the original pipeline, the updated pipeline further filtered the mismatch based on the CIGAR string and only mismatches with “CIGAR = M” were kept. As shown in the result, this optimized pipeline efficiently removed the unaligned mismatches enriched at the 5’ and 3’ end of reads. Normalized mismatch rates in each bin, the percentage of each type of mismatch in all sequencing bases within the bin. **g-h.** Statistics of T>C mutations in *PerturbSci-Kinetics* reads. Histogram showing the number of T>C mutations on reads that are identified to be from newly synthesized transcripts (g). For each read with high-quality mismatches identified, the fraction of mismatches from T>C mutations was calculated, which clearly separated the reads with background mutations and mutants introduced by 4sU in the plot (h). 30% was set as the cutoff to assign nascent reads as sci-fate^19^.

**Extended Data Fig. 5.**
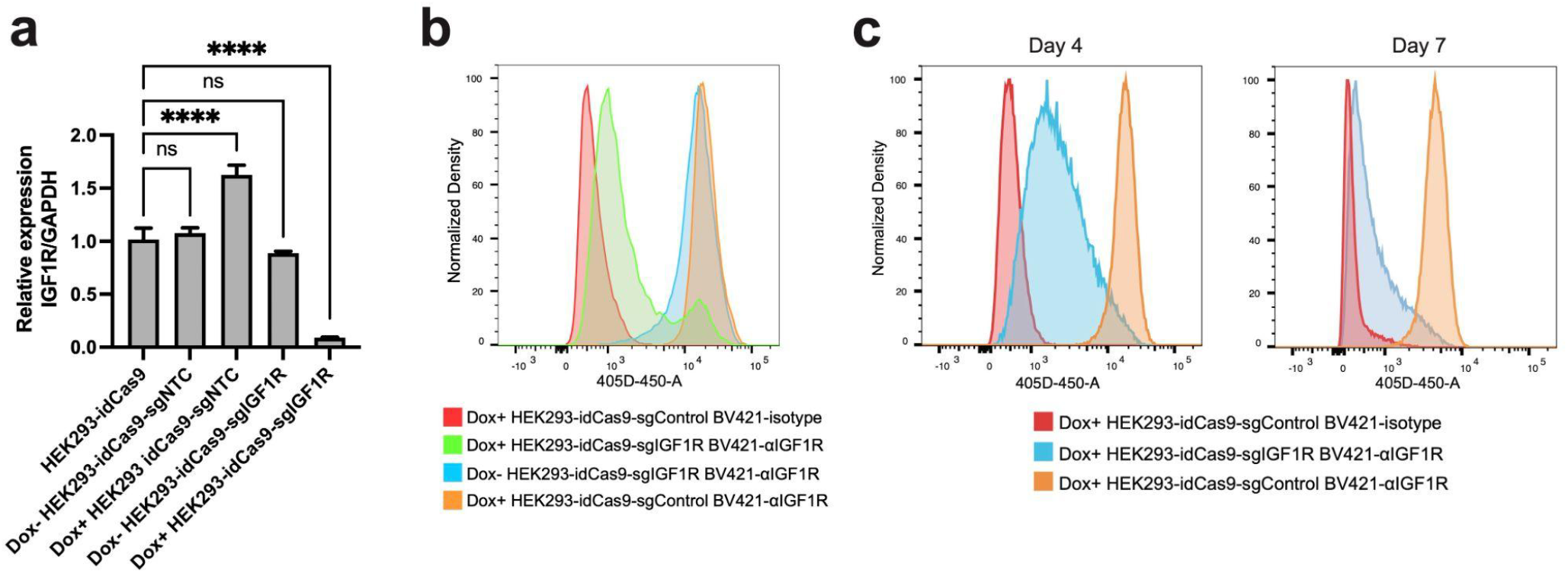
Validation of the CRISPRi performance. Inducible *IGF-1R* mRNA and protein knockdown in HEK293-idCas9-sgIGF1R cells were further validated by **a.** RT-qPCR after 3-day Dox induction (n=4, Dunnett’s test after one-way ANOVA was performed. ****, p-value < 1e-4; ns, no statistical significance.) and **b.** flow cytometry after 7-day Dox induction. Isotype, isotype control. αIGFIR, anti-IGF1R. **c.** Flow cytometry detection of IGF1R protein abundance after 4/7-day Dox induction in sgIGF1R and sgControl cells.

**Extended Data Fig. 6.**
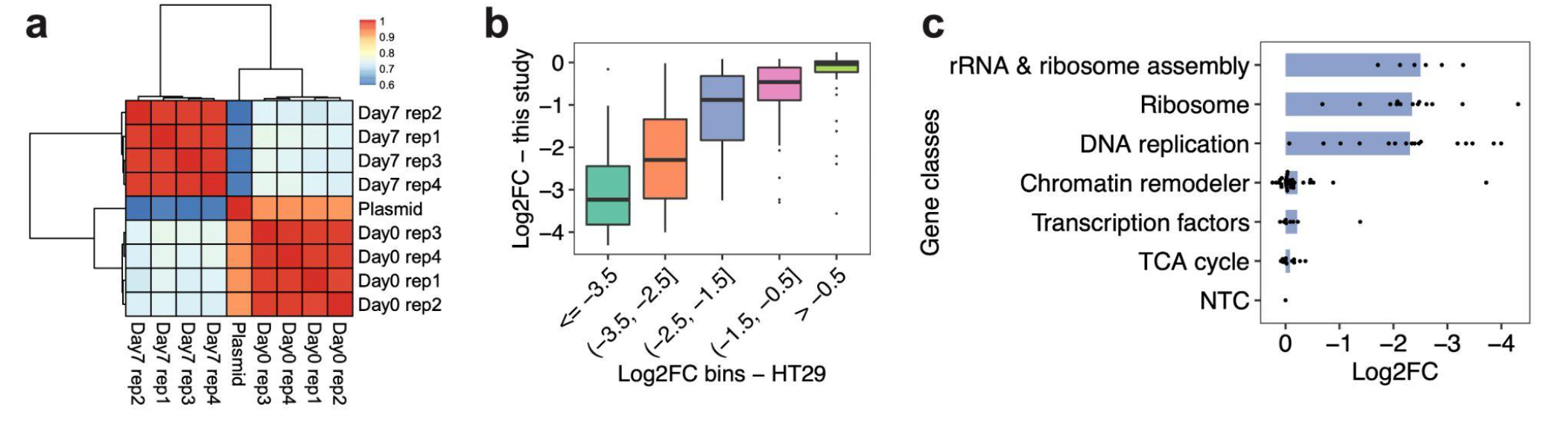
The changes in sgRNA abundance are consistent between replicates and previously published data. **a**. Heatmap showing the overall Pearson correlations of normalized sgRNA read counts between the plasmid library and bulk screen replicates at different sampling times. For each library, read counts of sgRNAs were normalized first by the sum of total counts and then by the counts of sgNTC. **b**. Box plot showing the reproducible trends of deletion upon CRISPRi between the present study and a prior report^29^. We calculated the fraction changes (After vs. before the CRISPRi induction) of sgRNAs for each gene, followed by log2 transformation. **c**. Bar plot showing the different extent of deletion of cells receiving sgRNAs targeting genes in different categories in the bulk screen. The knockdown on genes with higher essentiality caused stronger cell growth arrest.

**Extended Data Fig. 7.**
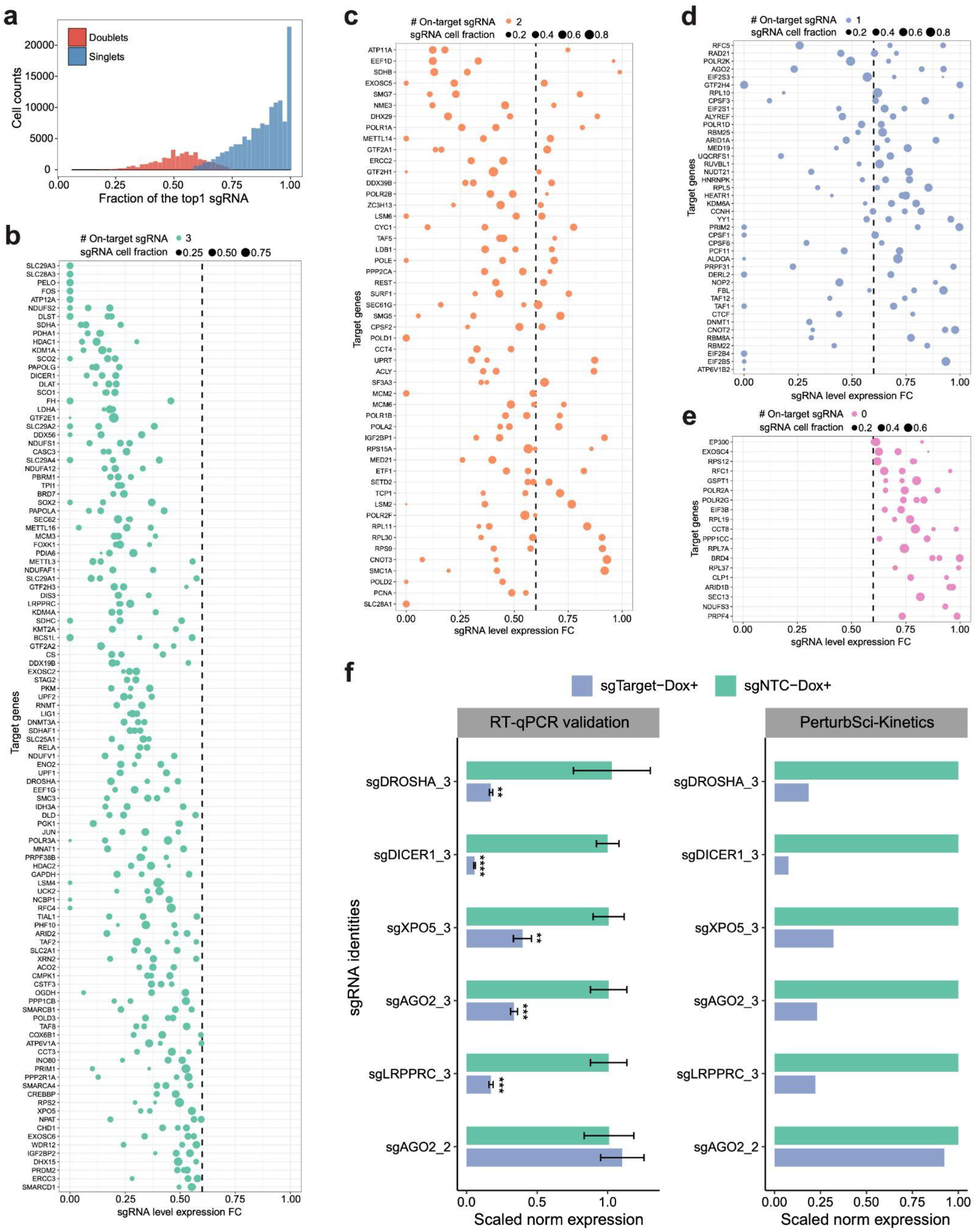
Quality control and sgRNA filtering on the *PerturbSci-Kinetics* library. **a**. We filtered out cells assigned to multiple gRNAs based on two criteria: the cell is defined as a sgRNA singlet if the most abundant sgRNA in the cell took >= 60% of total sgRNA counts and was at least 3-fold of the second most abundant sgRNA. The histogram shows the fraction distribution of the most abundant sgRNA in singlets (78%) and doublet cells (22%). **b-e**. Dotplots showing the expression fold changes of target genes upon CRISPRi induction compared to NTC. Each dot represents a sgRNA. Fold change < 0.6 was used for sgRNA filtering, and target genes with 3, 2, 1, 0 on-target sgRNA(s) were shown in b-e, respectively. FC, fold change. **f.** The accuracy of sgRNA targeting efficiency detected in *PerturbSci-Kinetics* was further confirmed by individual RT-qPCR validation (n=4). 5 sgRNAs with high efficiency and 1 off-target sgRNA were cloned to the modified CROP-seq-opti plasmid, and individual HEK293-idCas9 clones were established. RNA was extracted and RT-qPCR was conducted after 3-day Dox induction. ACTB was used as the internal reference in RT-qPCR. ****, p-value < 1e-4; ***, p-value < 1e-3; **, p-value < 1e-2. The comparison with no statistical significance was not marked. Mean expressions of target genes in NTC and corresponding cell populations in the original *PerturbSci-Kinetics* screen dataset were exhibited on the right for comparison.

**Extended Data Fig. 8.**
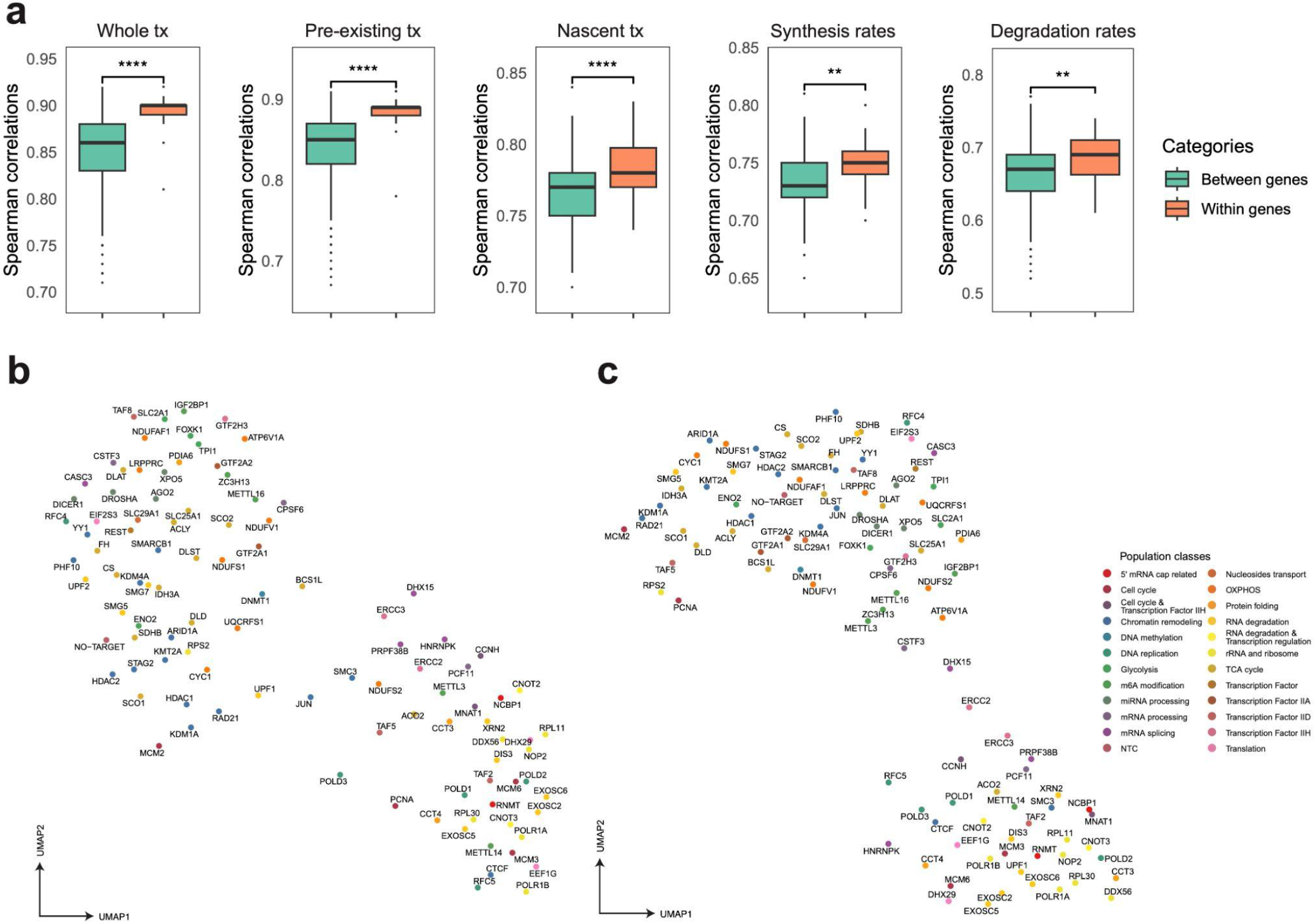
*PerturbSci-Kinetics* captures multi-layer transcriptome and RNA kinetics information upon perturbations. **a**. Boxplots showing the pairwise correlation coefficients of sgRNAs targeting the same gene or different genes, computed using aggregated whole transcriptomes, pre-existing transcriptomes, nascent transcriptomes, gene-specific synthesis rates and degradation rates. Considering the data sparsity and different cell numbers across perturbations, 150 cells per sgRNA were assembled into one pseudobulk for downstream analysis. Spearman correlation coefficients were calculated using DEGs between perturbations and NTC in the pooled screen. Compared with sgRNAs targeting different genes, the sgRNAs targeting the same genes showed significantly higher correlations in the aggregated whole RNA, preexist RNA, nascent RNA, synthesis rates, and degradation rates, confirming the data quality of our pooled perturbation. **b-c.** UMAP visualization of gene perturbations by inferred synthesis rates (b) or degradation rates (c). Differentially expressed genes between all perturbations-NTC pairs were combined, and their synthesis and degradation rates were calculated for each perturbation. For genes with no steady-state expression or no nascent counts, their synthesis and degradation rates were both assigned as 0. To denoise, only genes with inferred synthesis or degradation rate > 0 in at least 75% of pseudobulk cell populations were used for dimension reduction. The top 12 and 15 principal components from the synthesis and degradation rates matrix were used for UMAP visualization, respectively. These UMAPs still showed meaningful patterns. For example, RNA exosome genes (*e.g.,* EXOSC2, EXOSC5, EXOSC6), nonsense-mediated mRNA decay pathway members (*e.g.,* SMG5, SMG7), ribosomal biogenesis genes (*e.g.,* NOP2, RPL30, RPL11, POLR1A, POLR1B), miRNA biogenesis pathway members (*e.g.,*DICER1, DROSHA, XPO5, and AGO2) were in relative proximity in both UMAPs. Chromatin remodelers (*e.g.,* HDAC1, HDAC2, STAG2, RAD21, KMT2A, KDM1A, ARID1A) were closely clustered in synthesis rates-derived UMAP, while m6A regulators (*e.g.,* METTL3, METTL16, ZC3H13, IGF2BP1) and polyadenylation factors (*e.g.,* CPSF6, CSTF3) were closer to each other in degradation rates-derived UMAP.

**Extended Data Fig. 9.**
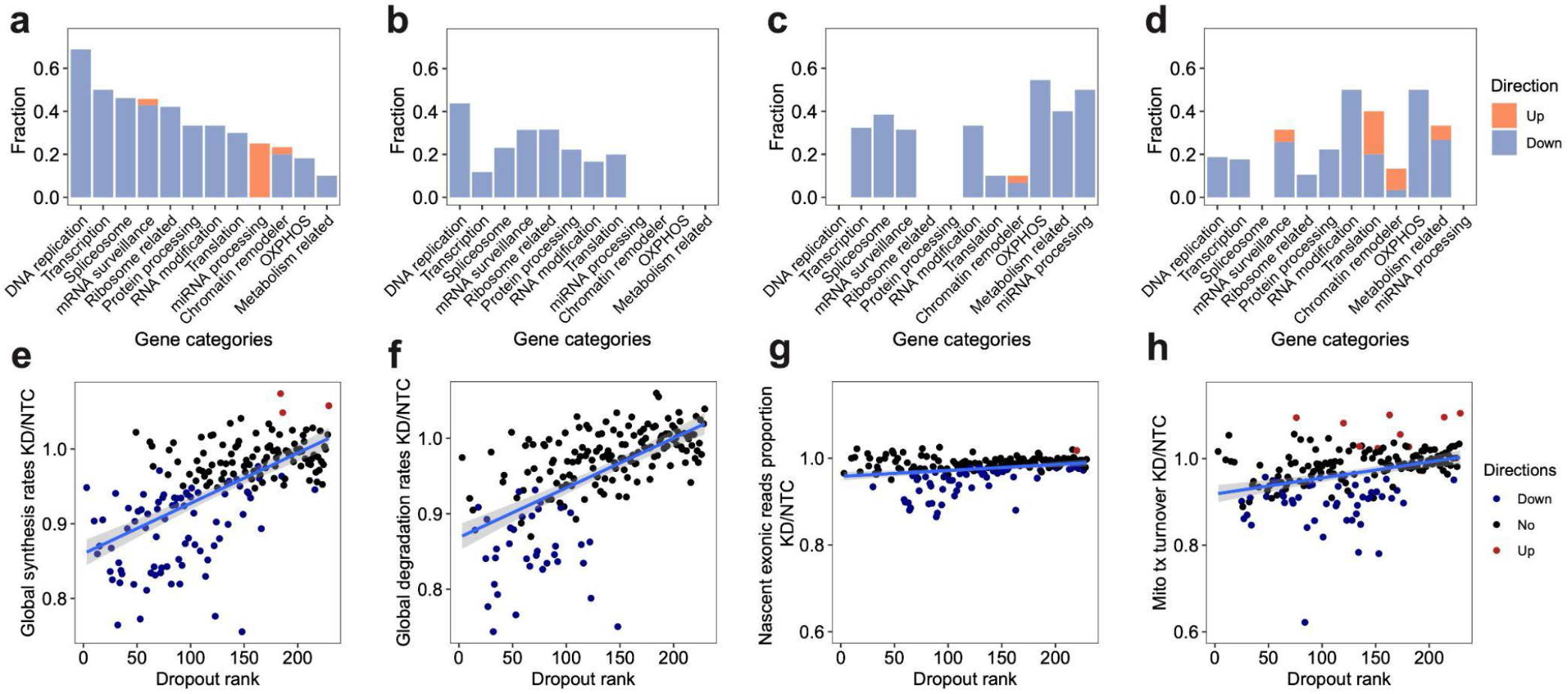
A systematic view of the effects of perturbations on global synthesis rates, global degradation rates, exonic reads ratio, and mitochondrial turnover rates. **a-d.** For each gene category, we calculated the fraction of genetic perturbations associated with significant changes in global synthesis rates (a), global degradation rates (b), proportions of exonic reads in the nascent transcriptome (c), and proportions of mitochondrial nascent reads (d). Overall global transcription could be affected by more genes than degradation. Perturbation on essential genes, such as DNA replication genes, could affect both global synthesis and degradation. Perturbations on chromatin remodelers only specifically impaired the global synthesis rates but not the degradation rates, supporting the established theory that gene expression is regulated by chromatin folding. In addition to the enrichment of genes in transcription, spliceosome and mRNA surveillance, perturbation on OXPHOS genes and metabolism-related genes also affected the RNA processing, consistent with the fact that 5’ capping, 3’ polyadenylation, and RNA splicing are highly energy-dependent processes. That knockdown of OXPHOS genes and metabolism-related genes could reduce the mitochondrial transcriptome dynamics and also supported the complex feedback mechanisms between energy metabolism and mitochondrial transcription^58^. **e-h.** Scatter plots showing the relationships between dropout effects and global synthesis rates (e), global degradation rates (f), proportions of exonic reads in the nascent transcriptome (g), and mitochondrial RNA turnover (h). Dropout rank, the ascending rank of gene-level sgRNA counts log2FC from the bulk screen. Directions were assigned as shown in Figure 2e-h. Both global synthesis and degradation rates showed strong negative correlations with dropout, indicating knocking out essential genes generally resulted in impaired global RNA synthesis and degradation. In contrast, proportions of exonic reads in the nascent transcriptome were much more stable across perturbations, and were only specifically affected by genes functioning in RNA processing. Proportions of mitochondrial nascent reads were also prone to be affected by genetic perturbation, but directions of changes depend more on the functions of perturbed genes than the essentiality of genes.

**Extended Data Fig. 10.**
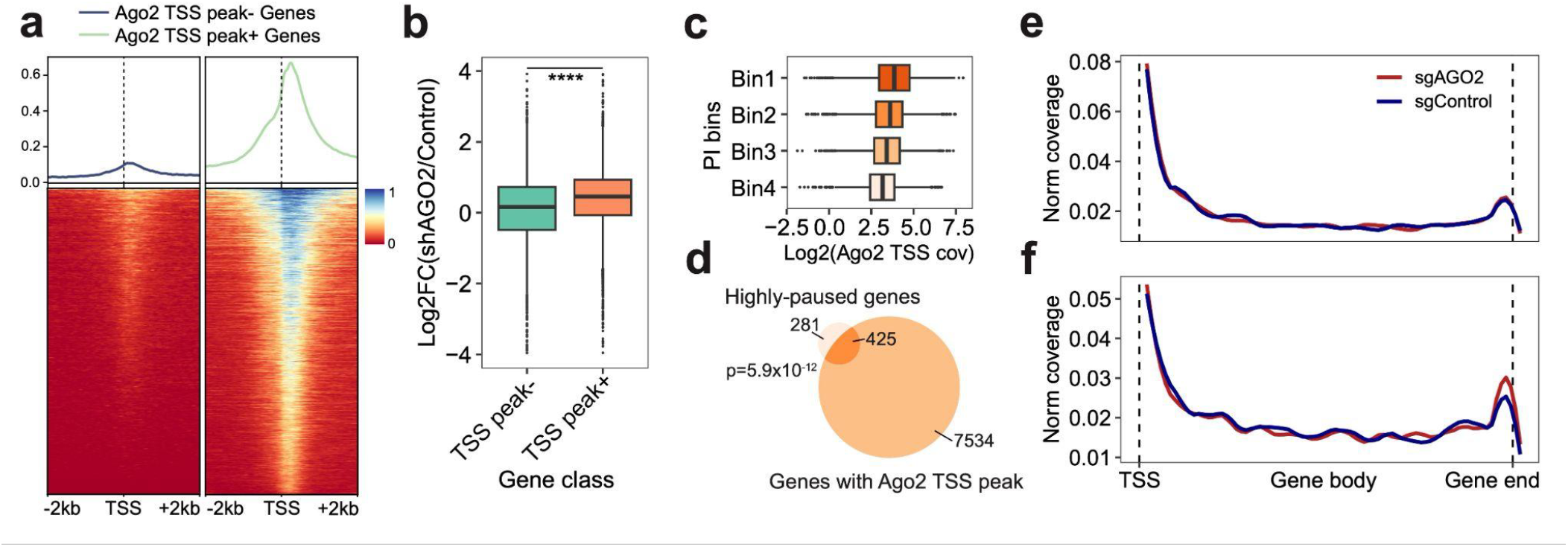
AGO2 functions as a transcriptional repressor by arresting transcription at the pausing status. **a**. The density plot (top) and heatmap (bottom) show the density of Ago2 ChIP-seq reads around TSS of genes with or without enriched Ago2 TSS binding peaks. **b.** We calculated the log2FC of gene expression between AGO2-silenced and control groups on genes with or without Ago2 TSS binding peaks. **c.** The box plot displays the positive correlation between PI of genes and normalized Ago2 ChIP-seq coverage within corresponding TSS regions. **c-d.** Genes were separated into 4 bins based on the average ranks of PI in two replicates (**Methods**). The Venn diagram highlights the significant association between Ago2 TSS binding and strong pausing status of genes. Highly-paused genes, genes with top 10% of average PI ranks. **e-f.** Highly-paused genes were split into two groups, 1) significantly-upregulated genes upon AGO2 knockdown or 2) genes without significant expression changes. We then calculated the nascent RNA coverage of these two groups of genes in sgNTC and sgAGO2 cells. Notably, only genes in group 1 displayed increased 3’ end enrichment upon AGO2 knockdown (f).

**Extended Data Fig. 11.**
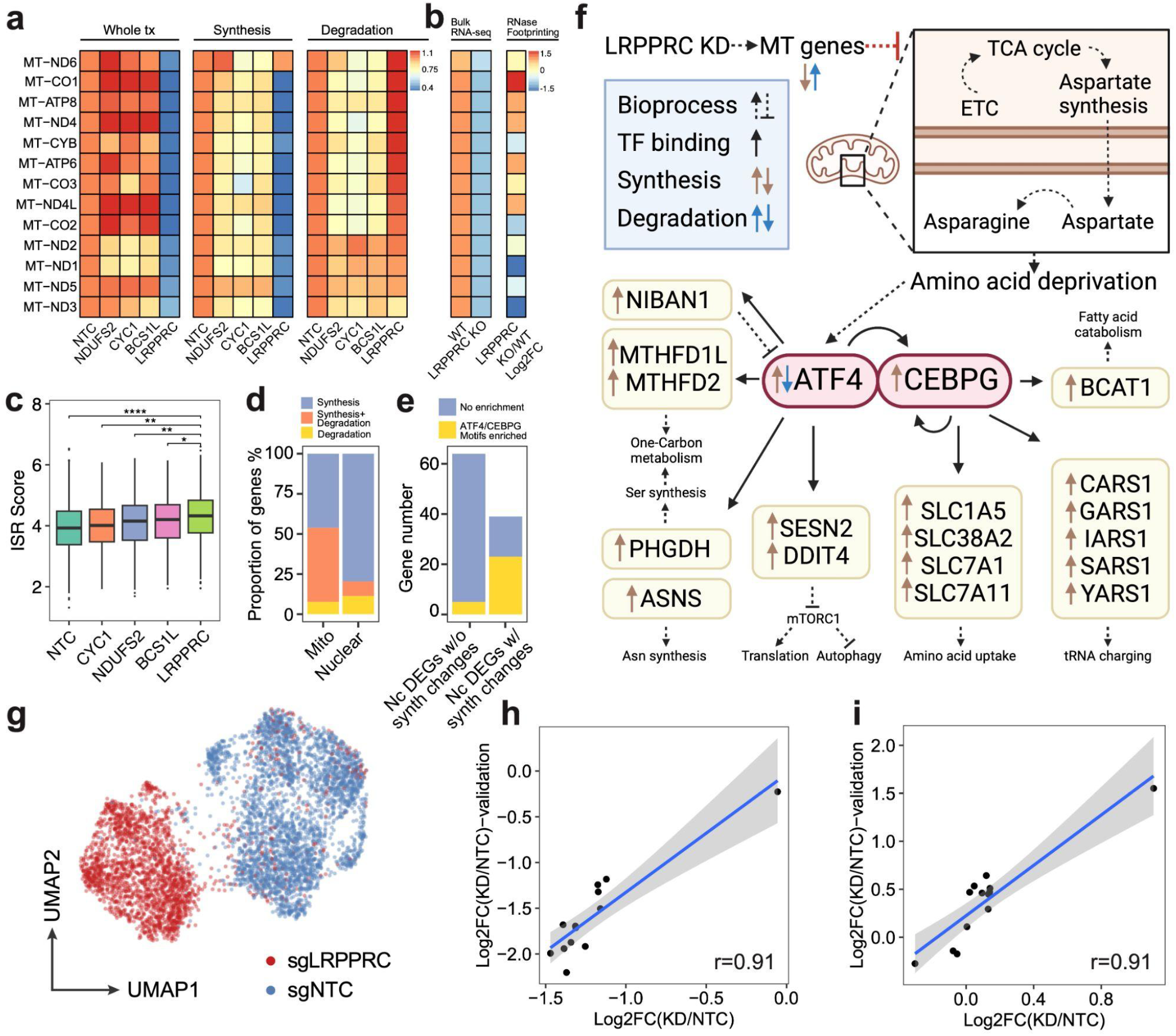
*PerturbSci-kinetics* identified LRPPRC as the master regulator of mitochondrial RNA dynamics. **a**. Heatmap showing the relative fold changes of gene expression, synthesis, and degradation rates of mitochondrial protein-coding genes upon NDUFS2, CYC1, BCS1L and LRPPRC knockdown compared to NTC cells. Perturbation on genes encoding electron transport chain components resulted in stable steady-state expression with impaired turnover. However, LRPPRC knockdown significantly disrupted the mitochondrial transcriptome dynamics by inhibiting the synthesis of almost all mitochondrial protein-coding genes and promoting the degradation of multiple genes including *MT-ND6, MT-CO1, MT-ATP8, MT-ND4, MT-CYB* and *MT-ATP6*. **b.** The heatmap on the left showed mean z-scored mitochondrial protein-coding gene expression changes between wild-type and *LRPPRC*-knockout mice heart tissue samples, as reported by Siira, S.J., et al. ^43^. The heatmap on the right showed the extent of the mRNA secondary structure increase upon *Lrpprc* knockdown observed in the same prior study^43^, which positively correlated with the elevated degradation rates of genes detected in our study (coefficient of Pearson correlation = 0.708, p-value = 6.8e-3). These results further validated the mRNA-stabilizing role of *Lrpprc* in regulating mitochondrial transcriptome. **c.** Boxplot showing the distribution of integrated stress response scores of single cells received different perturbations. ISR, integrated stress response. ISR score, average normalized expressions of genes within the ISR transcription program identified by Genome-wide Perturb-seq^8^. ****, p-value < 1e-4; **, p-value < 1e-2; *, p-value < 0.05. **d.** Bar plot showing the fraction of genes regulated by synthesis, degradation or both in mitochondrial-encoded and nuclear-encoded DEGs. **e.** Bar plot showing the enrichment of *ATF4/CEBPG* motifs at promoter regions of DEGs with or without significant synthesis changes. Nc DEGs w/o synth changes, Nuclear-encoded differentially expressed genes without synthesis changes. Nc DEGs w/ synth changes, Nuclear-encoded differentially expressed genes with synthesis changes. A large part of synthesis-regulated nuclear-encoded DEGs showed motif enrichment, suggesting the activation of an integrated stress response transcriptional program mediated by ATF4/CEBPG upon LRPPRC knockdown^59^. 5kb regions around transcription start sites of input genes were used for motif scanning and enrichment calculation using RcisTarget^60^. We identified two transcription factors (*ATF4* and *CEBPG*) that were i) significantly upregulated upon *LRPPRC* knockdown ii) significantly over-represented in the surroundings of the transcription start site of the synthesis-regulated nuclear-encoded DEGs (Normalized motif enrichment score of 16 for *ATF4* and 16.6 for *CEBPG*). **f.** The transcriptional regulatory network in *LRPPRC* perturbation inferred from our analysis. Notably, it was consistent with the prior study^59^ that *ATF4* was regulated at both transcriptional and post-transcriptional levels. **g.** Single-cell UMAP embedding of HEK293-idCas9-sgNTC and sgLRPPRC cells in the validation dataset. **h-i.** Correlations of synthesis rate and degradation rate changes of mitochondrial mRNA upon LRPPRC knockdown between the original screen and the validation dataset. r, coefficient of Pearson correlation.

**Extended Data Fig. 12.**
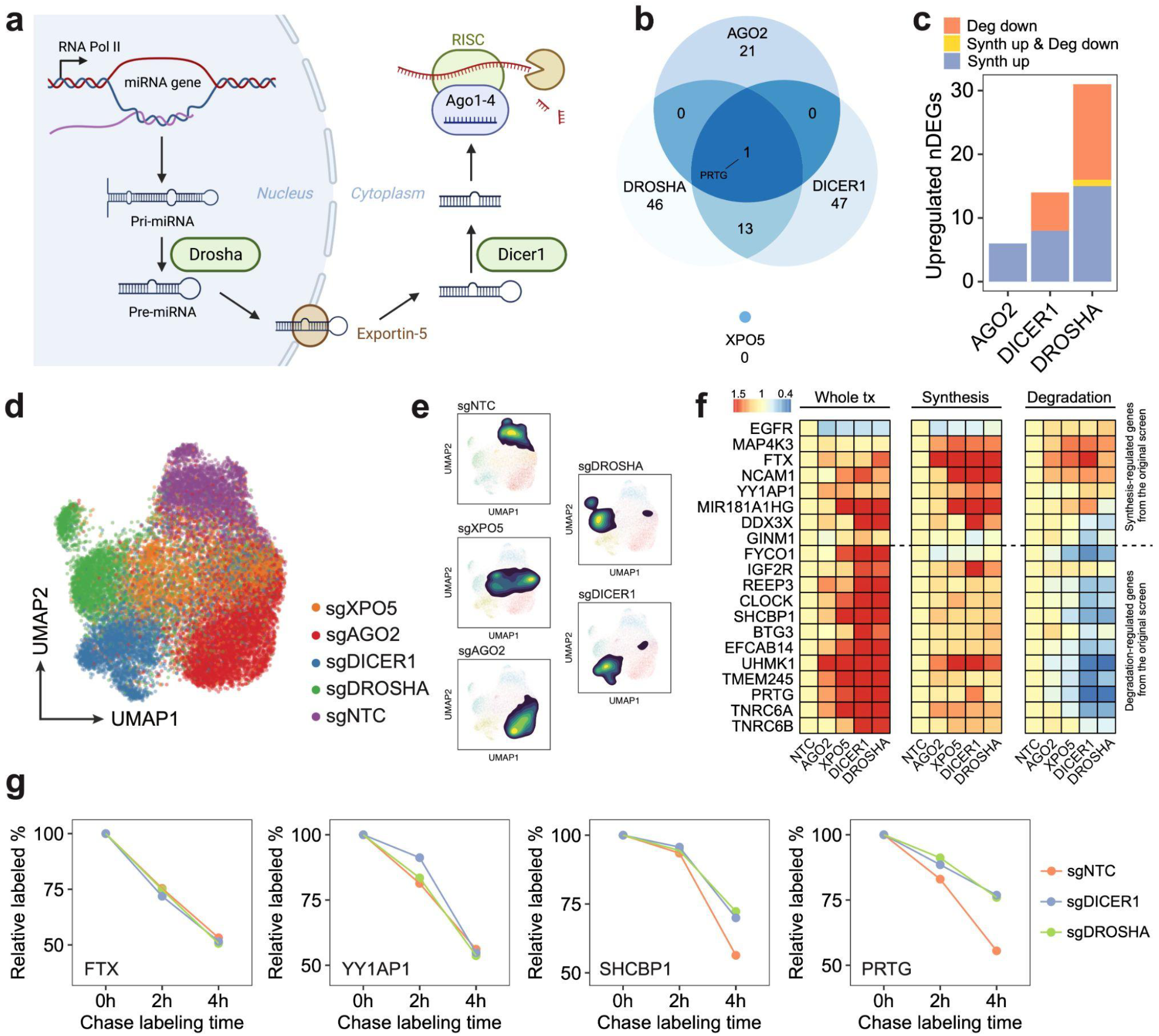
The overview of the miRNA biogenesis pathway and the validation on miRNA pathway perturbations. **a**. Illustration of the canonical miRNA biogenesis pathway. After the transcription of miRNA host genes, the primary miRNA (pri-miRNA) forms into a hairpin and is processed by *Drosha*. Processed precursor miRNA (pre-miRNA) is transported to the cytoplasm by Exportin-5. The stem loop is cleaved by *Dicer1*, and one strand of the double-stranded short RNA is selected and loaded into the RISC for targeting mRNA^46^. **b.** Venn diagram showing the overlap of upregulated DEGs across perturbations on four genes encoding main members of the miRNA pathway. The knockdown of two critical RNases in this pathway (*i.e., DROSHA* and *DICER1*) resulted in significantly overlapped DEGs (p-value = 2.2e-16, Fisher’s exact test). In contrast, *AGO2* knockdown resulted in more unique transcriptome features, and only 1 DEG (*PRTG*, identified to be mainly regulated by degradation and has been reported as a miRNA target^61^) overlapped with DEGs from *DROSHA* and *DICER1* knockdown, indicating the RNAi-independent roles of *AGO2*. Interestingly, *XPO5* knockdown showed no upregulated DEGs, which is consistent with a previous report in which *XPO5* silencing only minimally perturbed the miRNA biogenesis, indicating the existence of an alternative miRNA transportation pathway^47^. **c.** Bar plot showing the fraction of upregulated DEGs driven by synthesis changes and degradation changes upon *DROSHA*, *DICER1*, and *AGO2* perturbations. While *DROSHA* and *DICER1* knockdown resulted in increased synthesis and reduced degradation, *AGO2* knockdown only affected gene expression transcriptionally, which was consistent with the previous finding that *AGO2* knockdown resulted in a global increase of synthesis rates (Fig 2e), and further supported its roles in nuclear transcription regulation^62–64^. As *Drosha* is upstream of *Dicer1* in the pathway, we indeed observed stronger effects of *DROSHA* knockdown than *DICER1* knockdown, which was supported by the previous study^47^. **d-e.** UMAP embedding of NTC cells and single cells with individual miRNA biogenesis pathway genes knockdown (d). Perturbed cells exhibited distinct transcriptomic features and low-dimensional space distributions (e). **f.** Steady-state expression, synthesis rate, and degradation rate changes of synthesis/degradation-regulated genes showed high consistencies between the validation dataset and the original screen. **g.** Examples showing unchanged (transcription-regulated genes: FTX, YY1AP1) and enhanced (degradation-regulated genes: SHCBP1, PRTG) mRNA stability upon DROSHA/DICER knockdown. After long term 4sU labeling on Dox-induced HEK293-idCas9-sgNTC, sgDROSHA, sgDICER cells, chase labeling was performed. 3’end SLAM-seq was used to directly track the degradation of labeled mRNA. The fraction of labeled read counts of individual genes at each time point were divided by their labeled fractions at 0h for normalization.

**Extended Data Fig. 13.**
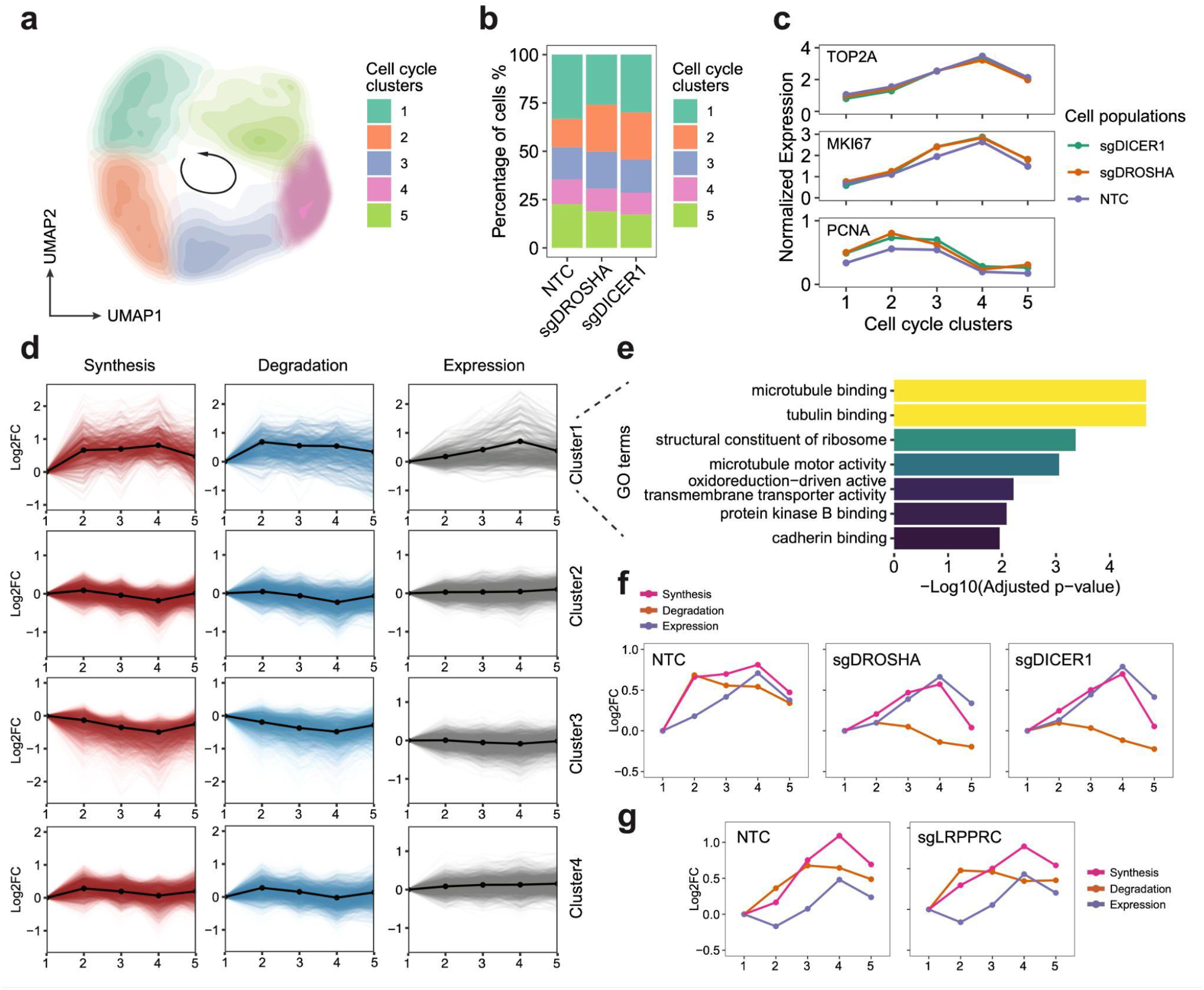
*PerturbSci-Kinetics* enables dissecting the effects of perturbations on cell cycle-dependent RNA dynamics. **a**. UMAP embedding of miRNA pathway genes-knockdown cells and NTC cells reflected the cell-cycle progression. **b.** Stacked barplot showing the cell cycle distribution of cells from each perturbation. **c.** The expression changes of cell cycle marker genes in cell cycle clusters. **d.** The cell cycle time-course synthesis rates, degradation rates, and expression levels of 4 gene clusters. Solid lines with dots, the mean values and the average trend of all genes within the cluster. **e.** Highly enriched GO terms of gene cluster 1 in GO enrichment analysis. **f.** Averaged trends of cell cycle time-course synthesis rates, degradation rates, and expression levels of cluster 1 genes in HEK293-idCas9-sgNTC, sgDROSHA, sgDICER1 cells. **g.** Averaged trends of cell cycle time-course synthesis rates, degradation rates, and expression levels of cluster 1 genes in NTC and sgLRPPRC HEK293-idCas9 cells. Considering potential strong batch effects from distinct genetic perturbation, cell cycle clustering analysis in (g) was performed independently of (a). Cell cycle clusters in (g) were not fully synchronized with clusters in (f).

## Materials and Methods

### Cell culture

The 3T3-L1-CRISPRi cell line was obtained from the Tissue Culture facility at the University of California, Berkeley. The HEK293 cell line was a gift from the Scott Keeney Lab at Memorial Sloan Kettering Cancer Center. The HEK293T cell line and the NIH/3T3 cell line were obtained from ATCC. All cells were maintained at 37 °C and 5% CO2 in high glucose DMEM medium supplemented with L-Glutamine and Sodium Pyruvate (Gibco 11995065) and 10% Fetal Bovine Serum (FBS; Sigma F4135). When generating a monoclonal cell line, the medium was supplemented with 1% Penicillin-Streptomycin (Gibco 15140163). In the screening experiment, sgRNA-transduced HEK293-idCas9 cells were cultured in high glucose DMEM medium supplemented with L-Glutamine (Gibco 11965092) and 10% FBS, following the induction of dCas9-KRAB-MeCP2 expression by 1ug/ml Dox (Sigma D5207),

### Cell lines generation

To generate HEK293 with Dox-inducible dCas9-KRAB-MeCP2 expression, the lentiviral plasmid Lenti-idCas9-KRAB-MeCP2-T2A-mCherry-Neo was constructed. A dCas9-KRAB-MeCP2-T2A insert was amplified from dCas9-KRAB-MeCP2 (Addgene #110821). A T2A-mCherry Gblock was synthesized by IDT. Gibson Assembly reaction (NEB E2611S) was performed at 50 °C with a mixture of Bsp119I-digested Lenti-Neo-iCas9 (Thermo FD0124; Addgene #85400), dCas9-KRAB-MeCP2-T2A amplicon, T2A-mCherry Gblock for 60 minutes to construct a dCas9-KRAB-MeCP2-T2A-mCherry plasmid. The reaction product was transformed into NEBstable competent cells (NEB C3040H), and colonies were inoculated and amplified in LB medium (Gibco 10855001) with 50ug/ml Sodium Ampicillin (Sigma A8351) at 37 °C overnight.

After plasmid extraction (QIAGEN No.27106) and sequencing validation, the plasmid was co-transfected with psPAX2 (Addgene #12260) and pMD2.G (Addgene #12259) into low-passage HEK293T cells in a 10cm dish using Polyjet (SignaGen SL100688) for 24 hours. Cells were gently washed twice with PBS, then cultured in a medium with 10mM Sodium Butyrate (Sigma TR-1008-G) for another 24 hours. The supernatant was collected, and cell debris was cleared by spinning down (5 minutes, 1000xg) and passed through a 0.45 μm filter. The lentivirus was concentrated 10x by the Lenti-X concentrator (TaKaRa 631231), and the virus suspension was flash frozen by Liquid Nitrogen and was stored at -80 °C.

The lentivirus titer was determined by examining the ratio of mCherry+ cells after 24 hours of transduction and 48 hours of Dox induction. Polybrene (Sigma TR-1003) at a final concentration of 8ug/ml was used to enhance the transduction efficiency. Then HEK293 cells were counted and transduced with lentivirus at MOI = 0.2 for 48 hours. Cells were treated with Dox for 48 hours, and the top 10% of cells with the strongest mCherry fluorescence were sorted to each well of a 96-well plate containing 100ul medium. After a 3-week expansion, monoclonal cells that survived were transferred to larger dishes for further expansion. We picked the clone with inducible homogeneous strong mCherry expression and normal morphology for the following experiment.

The polyclone 3T3-CRISPRi cell line was generated in a similar way. pHR-SFFV-dCas9-BFP-KRAB (Addgenes #46911) was co-transfected with psPAX2 and pMD2.G to generate dCas9-expressing lentivirus, and the transduction at MOI=0.2 was performed on 3T3 cells. BFP^hi^ cells (top 35% in BFP+population) were sorted and the sorting was repeated twice more after cell expansion to enrich cells with strong dCas9 expression.

### Gene Knockdown and efficacy examination

To simplify the lentiviral titer measurement, CROP-seq-opti-Puro-T2A-GFP was assembled by adding a T2A-GFP downstream of Puromycin resistant protein coding sequence on the CROP-seq-opti plasmid (Addgene #106280). Flanking MluI and CsiI digestion sites were added to the GFP Gblock (IDT) by PCR. Both amplicon and CROP-seq-opti vector were digested using MluI (Thermo, FD0564) and CsiI (Thermo, FD2114) at 37 °C for 30 minutes, and were ligated at room temperature for 20 minutes using the Blunt/TA Ligase Master Mix (NEB M0367S). Transformation, clone amplification, and sequencing validation were done as stated above.

Oligos corresponding to individual guides for ligation were ordered as standard DNA oligos from IDT with the following design:

Plus strand: 5’-CACCG[20bp sgRNA plus strand sequence]-3’

Minus strand: 5’-AAAC[20bp sgRNA minus strand sequence]C-3’

Oligos were reconstituted into 100uM and were mixed and phosphorylated using T4 PNK (NEB M0201S) by incubating at 37 °C for 30 minutes. The reaction was heated at 95 °C for 5 minutes and then ramped down to 25 °C by -0.1 °C/second to anneal oligos into a double-stranded duplex. The CROP-seq-opti-Puro-T2A-GFP was digested by Esp3I (NEB R0734L) at 37 °C for 30 minutes, then the linearized backbone and the annealed duplex were ligated at room temperature for 20 minutes using the Blunt/TA Ligase Master Mix (NEB M0367S). Transformation, clone amplification, sequencing validation, lentivirus generation, and titer measurement were done as stated above.

Mouse 3T3-L1-CRISPRi cells and 3T3-CRISPRi cells were transduced with the lentivirus expressing non-target control (NTC) sgRNA or sgRNA targeting *Fto*. Human HEK293-idCas9 cells were transduced with lentivirus expressing NTC sgRNA or sgRNA targeting *IGF1R* during technique development, and HEK293-idCas9-sgXPO5, sgAGO2, sgDROSHA, sgDICER1, sgLRPPRC cell lines were later established for validating significant hits from the screen. Transduction was carried out at MOI = 0.2 with 8ug/ml of Polybrene for 48 hours. Based on our puromycin titration experiments, sgRNA-transduced 3T3-L1-CRISPRi cells were selected by 2.5ug/ml Puromycin for 2 days and 2ug/ml Puromycin for 3 days, and sgRNA-transduced HEK293-idCas9 cells were selected by 1.5ug/ml Puromycin for 3 days and 1ug/ml Puromycin for 2 days. sgRNA-transduced 3T3-CRISPRi cells were directly sorted by gating on GFP fluorescence.

As dCas9-BFP-KRAB was constitutively expressed in 3T3-L1-CRISPRi cells and 3T3-CRISPRi cells, target genes started being silenced once sgRNA lentivirus was introduced. For HEK293-idCas9 cells, Dox treatment for a minimum of 72 hours was required before examining the knockdown effect.

For RT-qPCR validation, primer pairs targeting IGF1R, AGO2, XPO5, DROSHA, DICER1, LRPPRC, ACTB were selected from PrimerBank (https://pga.mgh.harvard.edu/primerbank/) and were synthesized from IDT. Total RNA in 1e6 cells of each sample was extracted using the RNeasy Mini kit (QIAGEN 74104) and the concentrations were measured by Nanodrop. 1ug total RNA was then reverse-transcribed into the first strand cDNA by SuperScript VILO Master Mix (Thermo 11755050). PowerTrack SYBR Green Master Mix (Thermo A46109) and PowerUp™ SYBR™ Green Master Mix (Thermo A25742) were used for RT-qPCR following the manufacturer’s instructions.

For flow cytometry validation, 1e6 cells of each sample were harvested and resuspended in 100ul of PBS-0.1% sodium azide-2% FBS. BV421 Mouse Anti-Human CD221 (BD 565966) and BV421 Mouse IgG1 k Isotype Control (BD 562438) at the final concentration of 10 ug/ml were added, and reactions were incubated at 4 °C in the dark with rotation for 30 minutes. Cells were then washed twice using PBS-0.1% sodium azide-2% FBS, and fluorescence signals were recorded.

### Construction of pooled sgRNA library

Genes to be included in our sgRNA library were carefully selected based on following considerations: 1) both essential and non-essential genes were included for comparison. These genes were identified using the bulk CRISPR screen data from the publication introducing the optimized CRISPRi sgRNA library^29^ and Depmap^65^. For example, knocking down ribosomal genes is usually fatal. 2) To validate the ability of *PerturbSci-kinetics* to characterize gene-specific RNA dynamics, we selected genes involved in transcription, chromatin remodeling, RNA processing, and mRNA decay based on Gene Ontology terms^66^ and KEGG pathways^67^. For instance, we included *CNOT2* and *CNOT3*, which are components of the key deadenylase complex for global mRNA degradation. 3) We ensured that all selected genes were expressed in the cell line to be used in our study. An in-house HEK293 EasySci-RNA dataset was used to select expressing genes that met criteria 1 and 2.

sgRNA sequences targeting genes of interest were obtained from an established optimized CRISPRi sgRNA library (only sgRNA set A was considered)^29^. Finally, 684 sgRNAs targeting 228 genes (3 sgRNAs/gene) and 15 non-targeting controls were included in the present study.

The single-stranded sgRNA library was synthesized in a pooled manner by IDT in the following format: 5’-GGCTTTATATATCTTGTGGAAAGGACGAAACACCG[20bp sgRNA plus strand sequence]GTTTAAGAGCTATGCTGGAAACAGCATAGCAAGTT-3’

100ng of oligo pool was amplified by PCR using primers targeting 5’ homology arm (HA) and 3’ HA with limited cycles (x12) to avoid introducing amplification biases. The PCR product was purified, and double-stranded library amplicons were extracted by DNA electrophoresis and gel extraction. Then the insert was cloned into Esp3I-digested CROP-seq-opti-Puro-T2A-GFP by Gibson Assembly (50 °C for 60 minutes). In parallel, a control Gibson Assembly reaction containing only the backbone was set. Both reactions were cleaned up by 0.75x AMPURE beads (Beckman Coulter A63882) and eluted in 5uL EB buffer (QIAGEN 19086), then were transformed into Endura Electrocompetent Cells (Lucigen, 602422) by electroporation (Gene Pulser Xcell Electroporation System, Bio-Rad, 1652662). After 1 hour of recovery at 250rpm, 37 °C, each reaction was spread onto an in-house 245 mm Square agarose plate (Corning, 431111) with 100ug/ml of Carbenicillin (Thermo, 10177012) and was then grown at 32 °C for 13 hours to minimize potential recombination and growth biases. All colonies from each reaction were scraped from the plate and the CROP-seq-opti-Puro-T2A-GFP-sgRNA plasmid library was extracted using ZymoPURE II Plasmid Midiprep Kit (Zymo, D4200). The lentiviral library was generated as stated above with extended virus production time. The step-by-step protocol is included in the supplementary materials.

### The pooled *PerturbSci-Kinetics* screen experiment

For each replicate, 7e6 uninduced HEK293-idCas9 cells were seeded. After 12 hours, two replicates were transduced at MOI=0.1 (1000x coverage/sgRNA) and another two replicates were transduced at MOI=0.2 (2000x coverage/sgRNA) with 8ug/ml of Polybrene for 24 hours. Then we replaced the culture medium with the virus-free medium and culture cells for another 24 hours. Transduced cells were selected by 1.5ug/ml of Puromycin for 3 days and 1ug/ml of Puromycin for 2 days. During the selection, we passed cells every 2 or 3 days to ensure at least 1000x coverage. At the end of the drug selection, we harvested 1.4e6 cells in each replicate (2000x coverage/sgRNA) as day0 samples of the bulk screen and pellet down at 500xg, 4 °C for 5 minutes. Cell pellets were stored at -80 °C for genomic DNA extraction later. Then the dCas9-KRAB-MeCP2 expression was induced by adding Dox at the final concentration of 1ug/ml, and L-glutamine+, sodium pyruvate-, high glucose DMEM was used to sensitize cells to perturbations on energy metabolism genes. Cells were cultured in this condition for additional 7 days and were passed every other day with 4000x coverage/sgRNA. On day7, 6ml of the original media from each plate was mixed with 6uL of 200mM 4sU (Sigma T4509-25MG) dissolved in DMSO (VWR 97063-136) and was put back for nascent RNA metabolic labeling. After 2 hours of treatment, 1.4e6 cells in each replicate were harvested as day7 samples of the bulk screen, and the rest of the cells were fixed and stored for single-cell *PerturbSci-Kinetics* profiling (see the next section).

Genomic DNA of bulk screen samples was extracted using Quick-DNA Miniprep Plus Kit (Zymo, D4068T) following the manufacturer’s instructions and quantified by Nanodrop. All genomic DNA was used for PCR to ensure coverage. The primer targeting the U6 promoter region with P5-i5-Read1 overhang and the primer targeting the sgRNA scaffold region with P7-i7-Read2 overhang was used for generating the bulk screen libraries for sequencing.

### Library preparation for the *PerturbSci-Kinetics*

After trypsinization, cells in each 10cm dish were collected into a 15ml falcon tube and kept on ice. Cells were spun down at 300xg for 5 minutes (4 °C) and washed once in 3ml ice-cold PBS. Cells were fixed with 5ml ice-cold 4% Paraformaldehyde (PFA) in PBS (Santa Cruz Biotechnology sc-281692) for 15 minutes on ice. PFA was then quenched by adding 250ul 2.5M Glycine (Sigma 50046-50G), and cells were pelleted at 500xg for 5 minutes (4 °C). Fixed cells were washed once with 1ml PBSR (PBS, 0.% SUPERase In (Thermo AM2696), and 10mM dithiothreitol (DTT; Thermo R0861)), and were then resuspended, permeabilized, and further fixed in 1ml PBSR-triton-BS3 (PBS, 0.1% SUPERase In, 0.2% Triton-X100 (Sigma X100-500ML), 2mM bis(sulfosuccinimidyl)suberate (BS3; Thermo, PG82083), 10mM DTT) for 5 minutes. Additional 4ml of PBS-BS3 (PBS, 2mM BS3, 10mM DTT) was then added to dilute Triton-X100 while keeping the concentration of BS3, and cells were incubated on ice for 15 minutes. Cells were pelleted at 500xg, 4 °C for 5 minutes and resuspended in 500ul nuclease-free water (Corning 46-000-CM) supplemented with 0.1% SUPERase In and 10mM DTT. 3ml of 0.05N HCl (Fisher Chemical, SA54-1) was added for further permeabilization. After 3 minutes of incubation on ice, 3.5ml Tris-HCl, pH 8.0 (Thermo 15568025), and 35ul of 10% Triton X-100 were added to each tube to neutralize the HCl. After spinning down at 4 °C, 500xg for 5 minutes, cells were finally resuspended in 400ul PSB-DTT at the concentration of ∼2e6 cells/100ul (PBS, 1% SUPERase In, 1% BSA (NEB B90000S), 1mM DTT), mixed with 10% DMSO, and were slow-frozen and stored in -80 °C.

The chemical conversion was performed before the library preparation. Cells were thawed with shaking in the 37 °C water bath and spun down, then were washed once with 400ul PSB without DTT. Next, cells were resuspended in 100ul PSB, mixed with 40ul Sodium Phosphate buffer (PH 8.0, 500mM), 40ul IAA (100mM, Sigma I1149-5G), 20ul nuclease-free water, and 200ul DMSO with the order. The reaction was incubated at 50 °C for 15 minutes and was quenched by adding 8ul 1M DTT. Then cells were washed with PBS and were filtered through a 20um strainer (Pluriselect 43-10020-60). Cells were finally resuspended in 100ul PSB.

For library preparation, a step-by-step protocol is included as a supplementary file. It is worth noting that *PerturbSci-Kinetics* is based on three levels indexing and can generate more than 5.6 million barcodes in a single full-scale experiment. In contrast, sci-fate is a two-level combinatorial indexing technique with a capacity of processing ∼5,000 cells in a single experiment^19^. scEU-seq is a "one-cell one-well" solution^23^ with a theoretical throughput of n plates x 96/384 wells, and Drop-seq-based scNT-seq^22^ normally processes ∼10,000 cells in each run. In the present proof-of-concept screen experiment, 576,000 cells in total were loaded for reverse transcription. After the second round of indexing (ligation), 69.5% - 74.5% of cells were recovered (413,980 cells in total). ∼248,388 cells were loaded for the third round of indexing, and finally 161,966 cells were computationally recovered after sequencing. The cell recovery rate of *PerturbSci-Kinetics* was significantly improved compared to 7% observed in sci-RNA-seq3^57^.

### 4sU pulse/chase labeling and SLAM-seq

HEK293-idCas9-sgAGO2 and sgNTC cells were induced with Dox for 7 days in 10cm dishes, during which the culture medium was replaced every 2 days to keep the stability of induction. Before harvesting cells, cells were labeled with 600uM 4sU for 20 minutes. The culture medium was then aspirated and cells were lysed by adding 600ul buffer RLT from RNeasy Mini kit with 1:100 V/V β-mercaptoethanol (Sigma M3148-250ML) to the dish. Lysate was scraped using cell scrapers (VMR 10062-906) and was then collected to a 1.5ml tube. Total RNA extraction was then done using the RNeasy Mini kit.

HEK293-idCas9-sgDROSHA, sgDICER1, and sgNTC cells were induced with Dox for 4 days in 10cm dishes. After seeded to a 6-well plate, cells were treated for another 3 days. By the end of the induction, the culture medium was replaced with Dox+ medium containing 100uM 4sU and cells were labeled for 18h. The medium was refreshed every 6h to keep the 4sU concentration stable. Then cells in each well were washed with PBS carefully once and the fresh medium containing 10mM uridine (Sigma U3750-1G) was added. Following 2h and 4h incubation, the medium was aspirated and cells were lysed with 250ul buffer RLT RNeasy Mini kit with 1:100 V/V β-mercaptoethanol. Total RNA was extracted using the RNeasy Mini kit, and RNA concentrations were measured by Nanodrop. Samples were stored at -80°C before further processing.

2-5 ug of total RNA from each sample was used for chemical conversion. RNA was diluted into 15ul, and mixed with 5ul of 100mM IAA, 5ul of NaPO4 (pH 8.0, 500mM) buffer, and 25ul of DMSO. The reaction was incubated at 50 °C for 15 minutes and was then quenched with 1ul 1M DTT. The RNA was purified using the Monarch RNA Cleanup Kit (NEB T2030L) and was eluted in 10ul of EB buffer. RNA concentrations were measured again using the Qubit RNA HS kit (Thermo Q32852) and samples were immediately used for library construction.

Full-length and 3’end bulk SLAM-seq were used for different experimental purposes. For full-length bulk SLAM-seq library construction, the CRISPRclean Stranded Total RNA Prep with rRNA Depletion Kit (Jumpcode Genomics KIT1014) was used. For 3’end bulk SLAM-seq library construction, an in-house 3’end library preparation workflow was used. In brief, 250-500ng total mRNA was mixed with 1ul 100uM oligodT primer (ACGACGCTCTTCCGATCTNNNNNNNNNNTTTTTTTTTTTTTTT), 1ul 10mM each dNTP mix, 0.5ul SUPERase In and the volume was adjusted to 15ul with molecular biology grade water. The reaction was Incubated at 55C for 5min and was then cooled on ice. 4ul 5xRT buffer and 1ul Maxima H Minus Reverse Transcriptase (Thermo EP0753) were added to the reaction, and reverse transcription was performed under the following program: 25 °C for 10min, then 50 °C for 15min. After 0.6x AMPURE beads purification, 20ul eluate was mixed with 14ul water, 4ul second strand synthesis (SSS) buffer, and 2ul SSS enzyme (NEB, E6111L). SSS was carried out by 1h incubation at 16 °C, then cDNA was extracted using another round of 0.6x AMPURE purification and the concentration was measured using Qubit dsDNA HS kit (Thermo Q32851). Read2 tagmentation was performed by mixing 10ng cDNA with 2xTD buffer (**Supplementary file 1**) containing 1:20 V/V Nextera Read2-Tn5 (**Supplementary file 1**) and incubating at 55 °C for 10 minutes. The reaction was quenched, and the final PCR was conducted in the same way as EasySci-RNA ^10^. The final library was purified using 0.8x AMPURE beads.

### Reads processing

For bulk CRISPR screen libraries, bcl files were demultiplexed into fastq files based on index 7 barcodes. Reads for each sample were further extracted by index 5 barcode matching. Then every read pair was matched against two constant sequences (Read1: 11-25bp, Read2: 11-25bp) to remove reads generated from the PCR by-product. For all matching steps, a maximum of 1 mismatch was allowed. Finally, sgRNA sequences were extracted from filtered read pairs (at 26-45bp of R1), assigned to sgRNA identities with no mismatch allowed, and read counts matrices at sgRNA and gene levels were quantified.

For *PerturbSci-Kinetics* transcriptome reads processing and whole-transcriptome/nascent transcriptome gene counting, the pipeline was developed based on *EasySci*^10^ and *Sci-fate*^19^ with minor modifications. After demultiplexing on index 7, Read1 were matched against a constant sequence on the sgRNA capture primer to remove unspecific priming, and cell barcodes and UMI sequences sequenced in Read1 were added to the headers of the fastq files of Read2, which were retained for further processing. After potential polyA sequences and low-quality bases were trimmed from Read2 by Trim Galore (0.6.7)^68^, reads were aligned to a customized reference genome consisting of a complete hg38 reference genome and the dCas9-KRAB-MeCP2 sequence from Lenti-idCas9-KRAB-MeCP2-T2A-mCherry-Neo using STAR (2.7.9a)^69^. Unmapped reads and reads with mapping score < 30 were filtered by samtools (1.13)^70^. Then deduplication at the single-cell level was performed based on the UMI sequences and the alignment location, and retained reads were split into SAM files per cell. These single-cell sam files were converted into alignment tsv files using the sam2tsv function in jvarkit (d29b24f)^71^. To minimize the impact of sequencing errors, we set thresholds on both the quality and the quantity of mismatches. First, we only considered mismatches with the CIGAR string “M”, and soft-clipped mismatches that failed to map to the reference genome in the head or at the end of reads were removed. After mismatches overlapping with intrinsic SNPs were removed, only mismatches with quality scores > 45 were used for 4sU mutation calling, as the probability of these mismatches to be sequencing errors were lower than 10^-4.5^ defined by the sequencer. Referred to sci-fate^19^, only reads with > 30% of T>C mutations among all mismatches were identified as nascent reads, and the list of reads was extracted from single-cell whole transcriptome sam files by the Picard MarkDuplicates program (2.27.4)^72^. Finally, single-cell whole transcriptome gene x cell count matrix and nascent transcriptome gene x cell count matrix were constructed by assigning reads to genes if the aligned coordinates overlapped with the gene locations on the genome. At the same time, single cell exonic/intronic read numbers were also counted by checking whether reads were mapped to the exonic or the intronic regions of genes. To quantify dCas9-KRAB-MeCP2 expression, a customized gtf file consisting of the complete hg38 genomic annotations and additional annotations for dCas9 was used in this step.

Read1 and read2 of *PerturbSci-Kinetics* sgRNA libraries were matched against constant sequences respectively, allowing a maximum of 1 mismatch. For each filtered read pair, cell barcode, sgRNA sequence, and UMI were extracted from designed positions. Extracted sgRNA sequences with a maximum of 1 mismatch from the sgRNA library were accepted and corrected, and the corresponding UMI was used for deduplication. De-duplication was performed by collapsing identical UMI sequences of each individual corrected sgRNA under a unique cell barcode. Cells with overall sgRNA UMI counts higher than 10 were maintained and the sgRNA x cell count matrix was constructed.

SLAM-seq reads were processed in a similar way with *PerturbSci-Kinetics*. In brief, for 3’end SLAM-seq, UMI sequences in Read1 were extracted and were attached to the headers of Read2 by UMI-tools (1.1.2)^73^, and only read2 were further processed. After polyA and low quality base trimming by Trim Galore, reads were aligned to the hg38 reference genome by STAR. To align reads from samples with high-concentration 4sU labeling, more loose alignment parameters were used (-- outFilterMatchNminOverLread 0.2 --outFilterScoreMinOverLread 0.2). Unmapped reads and reads with mapping score < 30 were filtered by samtools, and PCR duplicates in passed reads were further removed by UMI-tools. Nascent reads were identified and extracted, and gene counting on both whole transcriptome and nascent transcriptome were performed as mentioned above but at the sample level. For full-length SLAM-seq, reads were processed similarly but paired-end reads were retained. FeatureCounts (v2.0.1)^74^ was used for gene counting on these paired-end strand-specific libraries. A full list of software and their versions were included in the **Supplementary file 4**.

### Bulk screen sgRNA counts analysis

For each bulk screen library, read counts of sgRNAs were normalized first by the sum of total counts to remove the biases from sequencing depth, and then the abundance of each sgRNA relative to the sum of sgNTC was calculated, assuming the NTC cells had no selection pressure during the screen. The Pearson correlations across replicates were calculated based on the relative abundances. Then the fraction changes (After vs. before the CRISPRi induction) of sgRNAs were calculated within each replicate, and the mean fold changes across replicates were log2 transformed. The raw counts of another external bulk CRISPRi screen dataset^29^ was processed as stated above and the log2 mean relative abundance was compared to the current study.

### sgRNA singlets identification and off-target sgRNA removal

In the cell mixture experiments, cells with at least 200 whole transcriptome UMIs and 200 genes detected, and unannotated reads ratio < 40% were kept. If the count of the most abundant sgRNA was at least 3-fold of the second most abundant sgRNA within this single cell, then this cell was identified as a sgRNA singlet.

In the screen dataset, cells with at least 300 whole transcriptome UMIs and 200 genes detected, and unannotated reads ratio < 40% were kept. sgRNA identities of cells were assigned and doublets were removed based on the following criteria: the cell is assigned to a single sgRNA if the most abundant sgRNA in the cell took >= 60% of total sgRNA counts and was at least 3-fold of the second most abundant sgRNA. Then whole transcriptomes and sgRNA profiles of single cells were integrated with the matched nascent transcriptomes.

Target genes with the number of cells perturbed >= 50 were kept for further filtering. The knockdown efficiency was calculated at the individual sgRNA level to remove potential off-target or inefficient sgRNAs: whole transcriptome counts of all cells receiving the same sgRNA were merged, normalized by the total counts, and scaled using 1e6 as the scale factor, then the fold changes of the target gene expressions were calculated by comparing the normalized expression levels between corresponding perturbations and NTC. sgRNAs with >= 40% of target gene expression reduction relative to NTC were regarded as “effective sgRNAs”, and singlets receiving these sgRNAs were kept as “on-target cells”. Downstream analyses were done at the target gene level by analyzing all cells receiving different sgRNAs targeting the same gene together.

### Gene Ontology analysis of genes with high or low nascent reads ratio

To validate the specificity of 4sU labeling and the computational identification of nascent reads, we identified features of gene groups with different turnover rates. Single cells were split into nascent transcriptomes and pre-existing transcriptomes, and were loaded into Seurat^33^. Nascent transcriptomes and pre-existing transcriptomes were normalized, scaled independently, and DEGs between the two groups were identified by FindMarkers function^33^ with default parameters. Then GO enrichment analyses were performed using ClusterProfiler^75^ on upregulated genes (genes with significantly higher fraction of nascent counts, FDR of 0.05) and downregulated genes (genes with significantly lower fraction of nascent counts, FDR of 0.05) respectively.

### UMAP embedding on pseudo-cells

The count matrix of the “on-target” cells described above was loaded into Seurat^33^, and DEGs of each perturbation (compared to NTC) were retrieved by FindMarkers function^33^ with default parameters. Cells from perturbations with over one DEGs (by FindMarkers function^33^) were selected. We also included cells from genetic perturbations involved in similar pathways of the top perturbations. The fold changes of the normalized gene expression between perturbations and NTC were calculated, and were binned based on the gene-specific expression levels in NTC. The top 3% of genes showing the highest fold changes within each bin were selected and merged as features for Principal Component Analysis (PCA). The top 9 PCs were used as input for Uniform Manifold Approximation and Projection (UMAP) embedding (min.dist = 0.3, n.neighbors = 10).

### Differential expression analysis

In *PerturbSci-Kinetics*, pairwise differential expression analyses between each perturbation and NTC cells were performed by the differentialGeneTest() function of Monocle 2^76^. To identify DEGs with rate changes, we selected significant hits (FDR of 5%, likelihood) with a >= 1.5-fold expression difference and counts per million (CPM) >= 5 in at least one of the tested cell pairs. To showcase LRPPRC and miRNA pathway perturbations, more stringent criteria were used to obtain DEGs with high confidence: significant hits (FDR of 5%, likelihood) with a >= 1.5-fold expression difference and CPM >= 50 in at least one of the tested cell pairs were kept. EdgeR was used for bulk RNAseq DEGs analysis. Genes were firstly filtered by following thresholds: 1) a minimum of 10 raw counts in at least one sample; 2) genes were expressed in at least 50% of samples in each group. Then library sizes were normalized, and differential expression analysis was conducted. P-values were corrected by the Benjamini-Hochberg method and significant hits were selected at FDR < 5% level.

### Synthesis and degradation rates calculation

After the induction of CRISPRi for 7 days, we assumed new transcriptomic steady states had been established at the perturbation level before the 4sU labeling, and the labeling didn’t disturb these new transcriptomic steady states. The following RNA dynamics differential equation is used for synthesis and degradation rates calculation similar to the previous study^30^:

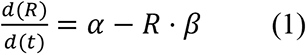

In which is the mRNA abundance of each gene, is the synthesis rate of this gene, and *β* is the degradation rate of this gene. Since the RNA synthesis follows the zero-order kinetics and RNA degradation follows the first-order kinetics in cells, 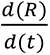 is determined by α and *R* ⋅ β.

As steady states had been established, the mRNA level of each gene didn’t change. We can get:

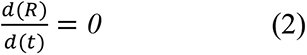

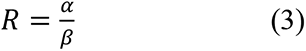

Under the assumption that the labeling efficiency was 100%, all nascent RNA were labeled during the 4sU incubation, and pre-existing RNA would only degrade. So, for nascent RNA (*R*_*n*_), *R*_*n*_(*t* = *0*) = *0* and *α*_*n*_ = *α*. For pre-existing RNA 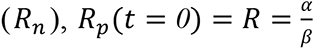 and *α*_*p*_ = *0*. Based on these boundary conditions, we could further solve the differential equation above on nascent RNA and pre-existing RNA of each gene.

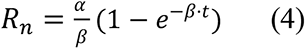

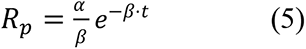

As both *R* and *R*_*n*_ were directly measured in *PerturbSci-Kinetics*, and cells were labeled by 4sU for 2 hours (*t* = *2*), *β* can be calculated from equation 3 and 4. Then *α* can be solved by equation 3.

Due to the shallow sequencing and the sparsity of the single cell expression data, synthesis and degradation rates of DEGs were calculated at the pseudo-cell level. We aggregated the expression profiles of all cells with the same target gene knockdown, normalized the expressions of genes by the sum of gene counts, and scaled the size of the total counts to 1e6. Synthesis and degradation rates of DEGs in the corresponding perturbed pseudo-cell were calculated as stated above. DEGs with only nascent counts or degradation counts were excluded from further examination since their rates couldn’t be estimated.

To examine the significance of synthesis and degradation rate changes upon perturbation, regarding the different cell sizes across different perturbations and NTC, which could affect the robustness of rate calculation, randomization tests were adopted. Only perturbations with cell number >= 50 were examined. For each DEG belonging to each perturbation, background distributions of the synthesis and degradation rate were generated: a subset of cells with the same size as the corresponding perturbed cells was randomly sampled from a mixed pool consisting of corresponding perturbed cells and NTC cells, then these cells were aggregated into a background pseudo-cell, and synthesis and degradation rates of the gene for testing were calculated as stated above, and the process was repeated for 500 times. Rates = 0 were assigned if only nascent counts or degradation counts were sampled during the process (referred to as invalid samplings), but only genes with less than 50 (10%) “invalid samplings” were kept for p-value calculation. The two-sided empirical p-values for the synthesis and degradation rate changes were calculated respectively by examining the occurrence of extreme values in background distributions compared to the rates from perturbed pseudo-cell. Rate changes with p-value <= 0.05 were regarded as significant, and the directions of the rate changes were determined by comparing the rates from the perturbed pseudo-cell with the background mean values. The fold changes of rates for each significant gene were calculated as follows: only NTC cells were sampled at the same size as perturbed cells and aggregated, and the background rates were calculated at the pseudo-cell level. After resampling for 200 times, these gene-specific rates were averaged. Fold changes of the rates = rates in perturbed pseudo-cell / mean rates from the NTC-only background.

Of note, in *PerturbSci-Kinetics*, we used oligo-dT primers for whole/nascent transcriptome capture, which only capture 3’ ends of transcripts and allow for gene-level feature counting, and most introns within gene bodies cannot be covered. Besides, the random tagmentation at the 3’ ends of the cDNA during the library preparation further confounds the 3’end exon-intron fraction, making it difficult to accurately infer the gene-specific splicing status. Additionally, splicing timings vary between different splicing sites within the same transcript^77^, and isoforms could have distinct splicing kinetics. Moreover, incorporating the splicing rate into our current rate estimation model is technically challenging, as the one-shot metabolic labeling design used in *PerturbSci-Kinetics* is incompatible with estimating the splicing rate mathematically. In fact, at least three labeling time points are needed to solve the differential equation for splicing rate estimation^30^.

### Global changes of key statistics upon perturbations

For global synthesis and degradation rate changes, considering the noise from lowly-expressed genes, we selected top1000 highly expressed genes from NTC cells, then calculated their synthesis rates and degradation rates in NTC cells and all perturbations with cell number >= 50. KS tests were performed to compare rate distributions between each perturbation and NTC cells.

During the reads processing, the number of reads aligned to exonic/intronic regions were counted at the single cell level. Then the distributions of exonic reads percentage in nascent reads from single cells with the same target gene knockdown and NTC cells were compared using the KS tests to identify genes affecting RNA processing.

The ratio of nascent mitochondrial read counts to total mitochondrial read counts was calculated in each single cell, and the distributions of the ratio from single cells with the same target gene knockdown and NTC cells were compared using the KS tests to identify the master regulator of mitochondrial mRNA dynamics.

In all global statistics examinations, the p-values were corrected from multiple comparisons, and comparisons with FDR <= 0.05 were considered as significant. The median value from each perturbation and NTC cells were compared to determine the direction of significant changes.

### Coverage analysis

To identify the potential different RISC binding patterns between synthesis/degradation-regulated DEGs in *DROSHA* and *DICER1* perturbations, we reprocessed the raw data of *Ago2* eCLIP obtained from Hela cells (two replicates, SRR7240709 and SRR7240710) from Zhang, K et, al^78^. Potential adapters at 3’ ends of reads were trimmed by Cutadapt^79^, and the first 6-base UMI were extracted and attached to headers of the reads. After STAR alignment^69^ and samtools filtering^70^, only uniquely aligned reads were kept and deduplication was performed based on the UMI and mapping coordinates using UMI-tools^73^. Then bam files were transformed to the single-base coverage by BEDtools^80^. The transcript regions of genes-of-interest were reconstructed based on the hg38 genome annotation gtf file from GENCODE. Briefly, for each gene, the exonic regions were extracted and were redivided into 5’UTR, CDS, and 3’UTR by the 5’most start codon and the 3’most stop codon annotated in the gtf. The *Ago2* binding coverages of these designated regions were obtained by intersection and were binned. A small background (0.1/base) was added for smoothing. The gene-specific signal in each bin was normalized by the number of bases in each bin, and the binned coverage of each gene was scaled to be within 0-1. After aggregating scaled coverages of synthesis/degradation-regulated genes respectively, the second scaling was performed to visualize the relative enrichment of *Ago2* binding at UTR compared to the CDS: fold changes of the scaled binned coverage relative to the lowest coverage value in the CDS along the aggregated transcript were calculated.

Meta-gene coverage analysis was conducted to visualize the gene body distribution of newly transcribed RNA in NTC and AGO2-knockdown samples. Genomic coordinates of protein coding genes on chromosome 1-22 and chromosome X were retrieved from the hg38 genome annotation gtf file from GENCODE in R. Gene bodies were binned into 50 bins by the tile() function in the GenomicRanges package, and an additional 200 bp bin downstream the end of genes were attached. These bins were ordered from TSS to gene ends and were exported as bed files separately by strandness using the export.bed() function in the rtracklayer package^81^. For input reads, two nascent reads BAM files per group from the pulse-labeling full-coverage SLAM-seq were merged using samtools, then reads with FLAG = 83 and 163 were extracted for coverage calculation of genes on the plus strand, and reads with FLAG = 99 and 147 were extracted for coverage calculation of genes on the minus strand. The gene-specific binned coverages were counted using the bedtools intersect command. Binned counts of each gene were normalized by total counts in the gene body, and the coverage of any group of genes was finally drawn by averaging the normalized signals across genes.

### Public ChIP-seq, shRNA RNA-seq, GRO-seq data analysis

Genes with detectable expression were identified from shControl/shAGO2 bulk RNA-seq in ENCODE. Processed gene counts quantification tables were downloaded from the ENCODE portal (ENCSR495YSS, ENCSR898NWE). Only genes with mean transcript per million (TPM) > 1 across 4 samples and with detected expression in at least 3 of 4 samples were included. Log2 fold changes of each gene upon AGO2 silencing were calculated by dividing the mean TPM in the shAGO2 group with the mean TPM in the shControl group.

Ago2 ChIP-seq bam and narrow peak files from ENCODE (ENCSR151NQL) were merged and were then used to identify TSS binding of Ago2. TSS regions of genes with detectable expression (defined as upstream 2kb to downstream 2kb around TSS) were retrieved by the promoters() function in the GenomicRanges package^82^. Genes were classified into Ago TSS peak+/- genes based on the overlap between their TSS regions with merged Ago2 ChIP-seq narrow peaks identified by findOverlaps() function. The Ago2 binding patterns of these two groups of genes were visualized using the computeMatrix function in deepTools^83^, and bed files containing coordinates of corresponding TSS regions and the merged ChIP-seq bam file were used as input.

GRO-seq data was downloaded from GEO (GSM2551016, GSM2551017) and were reprocessed to depict the transcriptional pausing status of genes. 3’end of reads were trimmed against polyA by Cutadapt^79^, and reads were then aligned to the hg38 reference genome using Bowtie2^84^. After filtering out unmapped reads using samtools, bam files were imported to R. TSS proximal regions and transcriptional elongation regions of protein coding genes with gene lengths >= 1kb were extracted, and the getPausingIndices() function from the BRGenomics package^85^ was used to calculate the pausing indices of genes. Genes detected in both replicates were ranked by the pausing index within the replicate, and an averaged rank was used to study the association with Ago2 TSS binding.

## Data Availability

The data generated by this study can be downloaded in raw and processed forms from the NCBI Gene Expression Omnibus (GSE218566, reviewers’ token: itqlgacczrgxpmb).

## Code Availability

The computation scripts for processing *PerturbSci-Kinetics* were included as supplementary files. Scripts and the user manual are available for open access in GitHub: https://github.com/JunyueCaoLab/PerturbSci_Kinetics

## Supplementary Tables (provided as Microsoft Excel files)

**Supplementary Table 1:** Genes and sgRNAs included in the study. Each gene (“gene_symbol”) has 3 sgRNAs, and they were named in the format “Gene_number” (“names”). sgRNA sequences were included in “sgRNA_seq”. The “gene_class” is the functional category of each gene.

**Supplementary Table 2:** Raw sgRNA counts of the bulk screen samples collected at different time points. Read counts of each sgRNA (“sgRNA_name”) from 4 replicates at day 0 and day 7 were included.

**Supplementary Table 3:** Relative sgRNA abundance fold changes between day 7 and day 0. The “Day7_vs_Day0_repX” is the fold changes of relative sgRNA abundance at the gene level (**Methods**).

**Supplementary Table 4:** Information about perturbations that showed significant global synthesis rate changes. The “adj.p” is the false discovery rate adjusted for multiple comparisons. The “direction” is the direction of the changes on the global synthesis rates distributions comparing perturbed cells to the NTC cells, and the “KD_median/NTC_median” is the quantitative measurement of the changes. The “gene_class” is the functional category of target genes (“Perturbations”).

**Supplementary Table 5:** Information about perturbations that showed significant global degradation rate changes. The “adj.p” is the false discovery rate adjusted for multiple comparisons. The “direction” is the direction of the changes on the global degradation rates distributions comparing perturbed cells to the NTC cells, and the “KD_median/NTC_median” is the quantitative measurement of the changes. The “gene_class” is the functional category of target genes (“Perturbations”).

**Supplementary Table 6:** Information about perturbations that showed significant nascent exonic reads ratio changes. The “adj.p” is the false discovery rate adjusted for multiple comparisons. The “direction” is the direction of the changes on the nascent exonic reads ratio distributions comparing perturbed cells to the NTC cells, and the “KD_median/NTC_median” is the quantitative measurement of the changes. The “gene_class” is the functional category of target genes (“Perturbations”).

**Supplementary Table 7:** Information about perturbations that showed significant mitochondrial RNA turnover changes. The “adj.p” is the false discovery rate adjusted for multiple comparisons. The “direction” is the direction of the changes in the distributions of mitochondrial nascent/total reads ratio comparing perturbed cells to the NTC cells, and the “KD_median/NTC_median” is the quantitative measurement of the changes. The “gene_class” is the functional category of target genes (“Perturbations”).

**Supplementary Table 8:** Steady-state expression and synthesis/degradation dynamics of mitochondrial genes upon *LRPPRC*, *NDUFS2*, *CYC1*, *BCS1L* perturbations. The “synth_rate”, “synth_FC”, “synth_pval”, “synth_direction” are the synthesis rate of the gene in the perturbed cells, the fold change of the synthesis rate of the gene in the perturbed cells compared to the NTC cells, the significance of the synthesis rate change, and the direction of the synthesis rate changes. The “deg_rate”, “deg_FC”, “deg_pval”, “deg_direction” are the degradation rate of the gene in the perturbed cells, the fold change of the degradation rate of the gene in the perturbed cells compared to the NTC cells, the significance of the degradation rate change, and the direction of the degradation rate changes. The “DEG_qval” and “DEG_fold.change” are the multiple comparison-corrected FDR and the fold change of the steady-state gene expression change in perturbed cells compared to the NTC cells.

**Supplementary Table 9:** Filtered differentially expressed genes between perturbations with cell number >= 50 and NTC. For each gene (“Gene_symbol”), the “perturbation” is the target gene in perturbed cells. The “DEGs_direction” is the direction of gene expression changes comparing perturbed cells to the NTC cells, and the “DEGs_FC” is the fold change of the gene expression changes comparing perturbed cells to the NTC cells. The “max.CPM.between.KD.NTC” and “min.CPM.between.KD.NTC” are the pseudobulk expression levels of the gene that showed higher and lower expression in perturbed cells or the NTC cells. The expression level was quantified by counts per million. The “qval” is the false discovery rate (one-sided likelihood ratio test with adjustment for multiple comparisons).

**Supplementary Table 10:** Differentially expressed genes with significant synthesis and/or degradation changes. The “perturbations” is the target gene of the perturbed cells, and the “Gene_symbols” is the symbols of DEGs with significant synthesis and/or degradation rate changes in corresponding perturbations. The type of significant rate change of each gene is included in the “Regulation_type”. The “Synth_deg_FC”, the “Synth_deg_direction”, and the “Synth_deg_pval” reflect the fold change, the direction of the change, and the randomization test p-value of the rate indicated in the “Regulation_type”. “DEGs_FC”, “DEGs_direction”, and “max.expr.between.KD.NTC” are the fold changes of gene expression, the direction of the change, and the maximum pseudobulk CPM between the corresponding perturbation and the NTC cells.

**Supplementary Table 11:** Steady-state expression and synthesis/degradation dynamics of merged DEGs upon *DROSHA* and *DICER1* perturbations. The “synth_rate”, “synth_FC”, “synth_pval”, “synth_direction” are the synthesis rate of the gene in the perturbed cells, the fold change of the synthesis rate of the gene in the perturbed cells compared to the NTC cells, the significance of the synthesis rate change, and the direction of the synthesis rate changes. The “deg_rate”, “deg_FC”, “deg_pval”, “deg_direction” are the degradation rate of the gene in the perturbed cells, the fold change of the degradation rate of the gene in the perturbed cells compared to the NTC cells, the significance of the degradation rate change, and the direction of the degradation rate changes. The “DEG_fold.change” and “DEG_qval” are the fold change of the steady-state gene expression change in perturbed cells compared to the NTC cells and the multiple comparison-corrected FDR.

## Supplementary files

**Supplementary file 1:** Detailed experiment protocols for *PerturbSci-Kinetics*, including all materials and equipment needed, step-by-step descriptions, and representative gel images.

**Supplementary file 2:** Primer sequences used in the *PerturbSci-Kinetics* experiment. The design principles and sequences of the oligo pool library, bulk screen sequencing primer, shortdT RT primers, sgRNA capture primers, ligation primers, sgRNA inner i7 primers, and P5/P7 primers were included. The columns indicate the positions on the 96-well plate (Well positions), an identifier of the sequence (Names), the full primer sequence (Sequences), and the barcode sequence (Barcodes).

**Supplementary file 3:** The overall costs for *PerturbSci-Kinetics* library preparation. Reagents used in each step were included, and the costs were calculated based on the scale of the real experiment.

**Supplementary file 4:** Computational pipeline scripts and notes for processing *PerturbSci-Kinetics* data, from sequencer-generated files to single-cell gene count matrix.

