## Supplementary file1 for "*PerturbSci-Kinetics*: Dissecting key regulators of transcriptome kinetics through scalable single-cell RNA profiling of pooled CRISPR screens"

Zihan Xu 2022/11/30

### PerturbSci-Kinetics library preparation protocol

#### Equipment:

- Centrifuge (Eppendorf 5702 RH)
- DynaMag-96 Side Skirted Magnet (Invitrogen, 12027)
- 12-tube Magnetic Separation Rack (NEB, S1509S)
- Eppendorf Mastercycler (4x)
- Freezer (-20C, -80C) and Refrigerator (4C)
- Gel Box
- Gel Imager
- Ice Buckets
- Microscope
- Multichannel Pipettes (2-20μL, 20-200μL) (Rainin Instruments)
- NextSeq 1000 Platform (Illumina)
- FreezeCell Cell Freezing Container (GeneSeeSci, catalog number: 27-802)
- Eppendorf ThermoMixer C (5382000023)

#### Primers and oligos:

The sequences of primers used are included in the supplementary material. All primers were ordered from IDT. Indexed primer plates (shortdT primer plate, sgRNA capture primer plate, ligation primer plate, EasySci-RNA/EasySci-ATAC P7 primer plates) were ordered in the solution form (100uM in TE buffer). Other primers are dissolved and diluted as stated in the protocol.

#### Materials:

- Nuclease free water (Corning, 46-000-CM)
- PBS 1X (Corning, 21-040-CV)
- 10cm cell culture dish (Genesee, 25-202)
- PluriStrainer Mini 40um (PluriSelect 43-10040-70)
- PluriStrainer Mini 20um (PluriSelect 43-10020-70)
- 4% PFA in PBS (Santa Cruz Biotechnology, sc-281692)
- SUPERase In RNase Inhibitor 20 U/uL (Thermo Fisher Scientific, AM2696)
- BSA 20 mg/ml (NEB, B9000S)
- Sodium phosphate monobasic dihydrate (Sigma, 71505-250G)
- Sodium phosphate dibasic dihydrate (Sigma, 71643-250G)
- 1M Tris-HCl, pH 7.5 (Thermo Fisher Scientific, 15567027)
- 1M Tris-HCl, pH 8.0 (Thermo Fisher Scientific, 15568025)
- 5M NaCl (Thermo Fisher Scientific, AM9759)
- 1M MgCl2 (Thermo Fisher Scientific, AM9530G)
- Dimethylformamide, 99.8% (Fisher Scientific, AC327175000)
- 4-thiouridine (Sigma, T4509-25MG)
- Glycine (Sigma, 50046-50G)
- Glycerol (Thermo Fisher Scientific, 15514011)
- DTT (dithiothreitol) (Thermo Fisher Scientific, R0861)
- Iodoacetamide (Sigma, I1149-5G)
- Dimethyl Sulfoxide (VWR, 97063-136)
- Triton X-100 for molecular biology (Sigma, 93443-100ML)
- 10mM dNTP (Thermo Fisher Scientific, R0192)
- Bis(sulfosuccinimidyl)suberate (Thermo Fisher Scientific, PG82083)

- 0.1N HCl (Fisher Chemical, SA54-1)
- Maxima H Minus Reverse Transcriptase with Buffer (Thermo Fisher Scientific, EP0753)
- SUPERase In (Thermo Fisher Scientific, AM2696)
- T4 DNA Ligase (NEB, M0202L)
- EDTA 0.5M Solution (VWR, 97062-656)
- Elution buffer (Qiagen, 19086)
- DAPI (10mg, Thermo Fisher Scientific, D1306)
- Falcon Tubes, 15 ml (VWR, 21008-936)
- Falcon Tubes, 50 ml (VWR, 21008-940)
- NEBNext® Ultra II Non-Directional RNA Second Strand Synthesis Module (NEB, E7550S)
- DNA binding buffer (Zymo Research, D4004-1-L)
- AMPure XP beads (Beckman Coulter, A63882)
- SDS, 20% Solution, RNase Free (Thermo Fisher Scientific, AM9820)
- Tween 20 (Sigma, P9416-100ML)
- Ethanol (Sigma, 459844-4L)
- NEBNext High-Fidelity 2X PCR Master Mix (NEB, M0541L)
- Qubit dsDNA HS kit (Invitrogen, Q32854)
- Qubit tubes (Invitrogen, Q32856)
- E-Gel EX Agarose Gel, 2% (Thermo Fisher Scientific, G402002)
- E-Gel 50bp DNA Ladder (Thermo Fisher Scientific, 10488099)
- 4-Chip Disposable Hemacytometers, Bulldog Bio (VWR, 102966-632)
- DNA LoBind Tube 1.5 ml, PCR clean (Eppendorf North America, 22431021)
- 1.0mL Self-Standing Cryovial (GeneSeeSci, catalog number: 24-200P)
- LoBind clear, 96-well PCR Plate (Eppendorf North America, 30129512)
- 0.2mL 8-Strip Tubes with Individual Caps (PCR Tubes) (Genesee, 27-125U)
- Reagent reservoirs (Fisher Scientific, 07-200-127)
- Falcon® 5mL Round Bottom w/ Cell Strainer (Fisher Scientific, 352235)
- eXTReme FoilSeal Film (Genesee, 12-156)
- eXTReme Clear Sealing Film (Genesee, 12-157)
- DNA Clean & Concentrator-5 (Zymo Research, D4014)
- Zymoclean Gel DNA Recovery Kit (Zymo Research, D4008)

#### Buffer and reagents preparation

##### DAPI stock preparation (Stored -20C)

- Dissolve 10mg DAPI in 2ml of nuclease-free water with a final concentration of 5mg/ml Split the DAPI solution into multiple tubes (100ul per tube).
- Take out one tube (100ul, 5mg/ml DAPI), add 1.9ml nuclease-free water. Split the diluted DAPI solution into multiple tubes (100ul per tube, 0.25mg/ml DAPI).
- Store the stock in the dark in -20C.

##### Sodium phosphate buffer (pH=8.0, 500mM) (Stored at RT)

- To prepare the stock solutions (500mM $NaH_2PO_4$ and 500mM $Na_2HPO_4$), dissolve 3.9g of $NaH_2PO_4$•$2H_2O$ (monobasic, MW = 156) in sufficient nuclease-free water to make a final volume of 50ml and dissolve 4.45g of $Na_2HPO_4$•$2H_2O$ (dibasic, MW = 178) in sufficient nuclease-free water to make a final volume of 50ml.
- Filter the buffer through the 0.2um strainer to new 50mL tubes using the syringe.
- Mix 9.5ml of $Na_2HPO_4$ with 500ul of $NaH_2PO_4$ to make 10mL of 500mM sodium phosphate buffer with pH=8.0.
- Take 2mL of the buffer for pH value measurement (My measurement is 7.95).

- Filter the remaining buffer (~8mL) through the 0.2um strainer to a new 15mL tube using the syringe. Store at room temperature.

**1M DTT (Stored at -20C)**
- Dissolve 1.54 g of DTT (MW=154) in 8 mL of $H_2O$.
- Adjust the total volume to 10 mL, dispense into 1-mL aliquots, and store them in the dark in -20C.

**10% (volume) Triton-X-100 in nuclease-free water (Stored at 4C)**
- Add 2mL Triton X-100 to 18mL nuclease-free water.
- Then rigorously vortex until a uniform solution is formed. The mix can be stored in 4C for up to 1 year.

**10% (volume) Tween-20 in nuclease-free water (Stored at 4C)**
- Add 2mL Tween-20 to 18mL nuclease-free water.
- Then rigorously vortex until a uniform solution is formed. The mix can be stored in 4C for up to 1 year.

**2x Tagmentation Buffer (Stored in -20C)**

| Reagents | Volume |
|---|---|
| 1M Tris HCl (pH 7.5) | 4ml |
| 1M MgCl2 | 2ml |
| DMF | 40ml |
| Nuclease-free water | 154ml |

- Aliquot the solution into 15mL or 1.5mL tubes and store at -20C.

**50x BS3 stock (Stored in -80C)**
- Dissolve 100mg BS3 (MW=572.43) into 1746uL nuclease-free water (~100mM, 50x).
- Split into 16 PCR tubes (~100ul/tube, can be used for fixing one sample) and store them in -80C for further use.

**Preparation of RT primer plate**
- Dilute sgRNA capture primers to 10uM: Add 45ul EB buffer to each well of a new 96-well plate. Then transfer 5ul 100uM sgRNA capture primers in the TE buffer to each well of the plate. Gently mix for several times and avoid any contamination during the pipetting.
- Add 8ul EB buffer to each well of another 96-well plate.
- Add 2ul 10uM sgRNA capture primer to each well.
- Add 10ul 100uM shortdT primers in TE buffer to each well. Gently mix for several time and avoid any contamination during the pipetting. Make sure barcodes of sgRNA capture primer and shortdT primer in each well correspond to each other.
- The final concentration of the primers: 1uM sgRNA capture primer + 50uM shortdT primer.
- The primer plate could be stored in 4C for 6 months or be stored in -20C for later use.

**Preparation of ligation adapter plate**
- In each well of an empty 96-well plate, add 5μL of 100uM Ligation Adaptor Primer and 5ul 100μM Barcoded Ligation Primers. Make sure to add the Barcoded Ligation Primers to their correct wells.

- Anneal the adaptor and ligation primers together by running the following thermocycler program:

| Temperature | Time |
|---|---|
| 95°C | 2 minutes |
| Ramp down to 20°C at the rate of -1°C/minute | \ |
| 4°C | Hold |

- The concentration of the annealed primer in each well is 50uM.
- Dilute the primers to 3.125uM by adding 150ul of EB buffer to each well. The resulting product is in stable, double-stranded form and can be stored at 4C or frozen. In 4C, the annealed primers should be stable for roughly three months and are suitable for short-term testing experiments.

**Preparation of Nextera-R2 Tn5**

- Protocol Derived from Hennig et al. 2018, Large-Scale Low-Cost NGS Library PreparationUsing a Robust Tn5 Purification and Tagmentation Protocol - purified Tn5 protein is also from this publication.
- We first mixed 150ul of 100uM Tn5-ME-B oligo (5'-GTCTCGTGGGCTCGGAGATGTGTATAAGAGACAG-3', in TE buffer) with 150ul of 100uM Tn5MErev oligo (-/5'Phos/CTGTCTCTTATACACATCT-3', in TE buffer) reaching a final concentration of 50uM.
- Then, we split the mixture into aliquots and performed the following thermocycler conditions: 95C for 5 minutes, slowly cooled to 65C (-0.1C/sec), 65C for 5 minutes, slowly cooled to 4C (-0.1C/sec).
- We further diluted the mixture to 35uM by mixing 10ul of the oligo mixture with 4.28ul of TE buffer. Then, we combined 1ul of the Tn5 enzyme at 4mg/mL with 19ul of Tn5 Dilution Buffer (25mM Tris pH 7.5, 800mM NaCl, 0.1mM EDTA, 1mM DTT and 50% glycerol) and 2ul of the 35uM Tn5-ME-B/Tn5-MErev oligo mixture.
- This solution was placed on a thermomixer at 23C for 30 minutes and diluted with 22ul of glycerol and stored at -20C for future usage.

#### Sample harvesting and cell fixation:

- After 4sU labeling, suction the culture media. Wash cells with 2ml PBS gently, then trypsinize cells for 3 minutes at 37C.
- Neutralize trypsin by adding 8ml DMEM+10%FBS. Separate cells well by pipetting for several times using serological pipette. Then quickly transfer cell suspension to a 15ml conical tube and keep the tube on ice.
- Spin cells down at 4C, 300g for 5 minutes.
- Aspirate the supernatant. Wash cells by resuspending cells in 3ml ice-cold PBS.
- Spin cells down at 4C, 300g for 5 minutes.
- Add 5ml of ice-cold 4%PFA in PBS to tubes.
- Resuspend cells thoroughly using P1000 pipette. Fix on ice for 15 minutes.
- Add 250ul 2.5M glycine to each tube. Mix well by pipetting for several times. Then spin down at 4C, 500g for 5 minutes.
- Aspirate the supernatant. Resuspend cells in 1ml PBS+0.1% SUPERase In+10mM DTT for washing. Then spin down at 4C, 500g for 5 minutes.

- After aspirating the supernatant, resuspend cells in 980uL PBS+0.1% SUPERase In+2mM BS3+10mM DTT, then quickly add 20ul 10% Triton-X100 into the tube, pipette for several times to mix well. Incubate on ice for 5 minutes for cell permeabilization.
- Add 4ml PBS+2mM BS3+10mM DTT to the tube, mix well by inverting the tube for several times, then put on ice for another 15 minutes for secondary fixation.
- Spin down at 500g, 4C for 5 minutes.
- Resuspend cells in 500uL nuclease free water+0.1% SUPERase In+10mM DTT.
- Add 3ml 0.05N HCl quickly using serological pipette, invert the tube several times, then incubate on ice for 3min.
- Add 3.5ml 1M Tris-HCl pH7.5 to each tube using serological pipette, pipette up and down for several times to neutralize HCl well. (V=7mL per tube)
- Add 35uL 10% Triton-X100 to each tube. Use wide-bore pipette tips to mix well.
- Spin down at 500g, 4C for 5 minutes.
- Resuspend cells in PSB with DTT (PBS+1% BSA + 1% SUPERase In+1mM DTT, DTT is not used in PSB for other parts of the protocol), then aliquot ~ 1-2*10^6 cells to each 1.5ml tube.
- **Stop point:** cells could be stored in -80C for a month. To freeze cells, add 1/10 volume of the DMSO and mix gently using the wide-bore pipette. Then transfer tubes to a freezing container and perform slow-freeze in -80C.

#### Single cell Perturb-Kinetics library preparation

##### Chemical conversion

- Thaw frozen cells: Take a tube of cells frozen in -80C, and thaw cells in the 37C water bath with shaking (V=~100ul).
- Add 400ul ice-cold PBS+1% BSA, mix well, then spin down at 4C, 500g for 5 minutes.
- Wash with 500ul ice-cold PSB once more to remove residual DTT. Spin down at 4C, 500g for 5 minutes. Resuspend cells in 100ul PSB.
- Prepare 100mM IAA: Dissolve 18.5 mg IAA (MW=184) in 1 mL of nuclease-free water. Keep away from the light until use. Prepare freshly before every experiment.
- Add 40ul Sodium phosphate buffer (pH 8.0, 500mM), 40ul IAA (100mM), 20ul nuclease-free water sequentially to the 100ul cell suspension. Mix gently.
- Mix 200ul DMSO with cells (V = 400ul), pipette gently to mix the reaction well.
- Incubate at 50C for 15 minutes. After the incubation, quickly transfer cells on ice.
- Quench the reaction by adding 8ul 1M DTT and mix gently.
- Transfer the mixture into 8.5ml ice cold PBS. Then rinse the wall of the reaction tube with PBS-cell suspension once. Combine it with the rest of the suspension. Add 45ul 10% Triton-X100 to the cell suspension, mix by inverting the tube for several times. Pellet the cells at 4C, 500xg for 5 minutes. Aspirate the supernatant.
- Re-suspend chemically converted cells with 500ul PBS+1% BSA to remove the residual chemicals. Pellet the cells at 4C, 500xg for 5 minutes and aspirate the supernatant.
- **Optional:** Resuspend cells in 500ul PBS+1% BSA, then filter cells through the strainer (20um-40um) by short centrifugation at 4C for 15 seconds. Wash the strainer once with 250ul PBS+1% BSA. Then pellet cells at 4C, 500xg for 5 minutes. Aspirate the supernatant.
- Resuspend the cells with 100ul PSB.
- Measure the cell concentration:
  - prepare the DAPI working solution: mix 1ul 0.25mg/ml DAPI with 500ul PBS (1:500 dilution). Keep it on ice and keep away from light.
  - Take 1ul cell suspension, mix with 9ul DAPI working solution (1:10 dilution).

- Load onto the hemocytometer and count the cell number under the fluorescent microscope.

#### Reverse transcription

- All operations are done on ice.
- Dilute cells to and mix with dNTP:
    - Adjust the concentration of cells to 1500 cells/ul in PSB.
    - For each well of the 96-well plate: 3000 cells in 2ul PSB + 0.25ul 10mM dNTP.
    - Premix the cells with dNTP based on the number of wells designed for samples. Aliquot cell suspension into an 8-well PCR strip.
    - Transfer 2.25ul to each well of a 96-well plate using the multi-channel pipette.
- Add 1ul premixed shortdT primers (50uM)+sgRNA capture RT primer (1uM) to each well of the 96-well plate using the multichannel pipette.
- After preheating the PCR thermocycler, incubate the plates at 55C for 5 min. After the reaction is cooled down to 4C, put the RT plate back on ice.
- Prepare the reverse transcription reaction mix during the incubation.

| Reagent | 1 well | For a 96-well plate (110 wells) |
|---|---|---|
| 5xMaxima H- RT Buffer | 1uL | 110 uL |
| Maxima H- RTase | 0.125uL | 13.8 uL |
| SUPERase In | 0.125 uL | 13.8 uL |
| Nuclease-free water | 0.5 uL | 54.9 uL |
| Final Volume | 1.75 uL | 192.5 uL |

- Split 24ul (in reality 23.6uL considering pipetting error) to each well of an 8-well PCR strip. Add 1.75uL to each well of the 96-well plate using a multichannel pipette and mix by pipetting gently for 3 times.
- Seal the plate with a plastic film. Then start the RT reaction using the following program:

| Temperature | Time |
|---|---|
| 20 °C | 4 minutes |
| 30 °C | 2 minutes |
| 40 °C | 2 minutes |
| 50 °C | 2 minutes |
| 55 °C | 15 minutes |

- After RT, put the RT plate back on ice.

#### Pool and wash cells

- For each 96-well plate, make 3mL PBS wash buffer with 0.05% Triton-X100 (PBB 0.05):

| Reagents | Volume |
|---|---|
| PBS | 3ml |
| BSA | 30ul |
| 10% Triton-X100 | 15ul |

- Also make 2mL PBS wash buffer with 0.1% Triton-X100 (PBB 0.1):

| Reagents | Volume |
|---|---|

| | |
|---|---|
| PBS | 2ml |
| BSA | 20ul |
| 10% Triton-X100 | 20ul |

- Prepare 40mM EDTA in PBB 0.1 (EDTA-PBB): add 56ul 500mM EDTA to 644ul PBB 0.1, mix well.
- Transfer EDTA-PBB into a basin, then add 5uL into each well of the RT plate using the multichannel pipette.
- Pool cells into 1 1.5ml tube using the wide-bore multichannel pipette.
- Pellet cells at 4C, 1000g for 3min. Aspirate the supernatant carefully.
- Resuspend cells in 500ul PBB 0.05.
- Pellet cells at 4C, 1000g for 3min. Aspirate the supernatant carefully.

#### Ligation

- Resuspend cells in 260uL PBB 0.05.
- Then split 32.5ul of cell suspension into each well of an 8-well PCR strip (in fact 32.2uL considering pipetting error).
- Distribute 2.5ul cell suspension to each well of a 96-well plate using the multichannel pipette.
- Take out the ligation adapter 96-well plate from 4C and keep it on ice, add 1μL of pre-annealed DNA ligation adapters (3.125uM) to each well of the ligation plate using the multichannel pipette.
- Prepare the ligation reaction mix for a 96 well plate:

| Reagent | 1 well | For a 96-well plate (110 wells) |
|---|---|---|
| 10x T4 ligation buffer | 0.5 uL | 55 uL |
| SUPERase In | 0.05 uL | 5.5 uL |
| T4 ligase | 0.5 uL | 55 uL |
| Nuclease-free water | 0.45 uL | 49.5 uL |
| Final Volume | 1.5 uL | 165 uL |

- Split 20.625ul (in fact 20.2uL considering error) to each tube of the 8-well PCR strip, and finally add 1.5uL into each well using a multichannel pipette with small tips.
- Incubate with gentle shaking (50rpm) on the shaker at room temperature for 30 minutes.
- Prepare EDTA-PBB before the end of the incubation: add 56ul 500mM EDTA to 644ul PBB 0.1, mix well.
- Transfer EDTA-PBB into a basin, then add 5uL into each well of the ligation plate.

#### Pool and wash cells

- Pool cells into 1 1.5ml tube using the wide-bore multichannel pipette.
- Pellet cells at 4C, 1000g for 3min. Aspirate the supernatant carefully.
- Resuspend cells in 500ul PBB 0.05.
- Pellet cells at 4C, 1000g for 3min. Aspirate the supernatant carefully.
- **Optional:** Resuspend cells in 500ul PBB 0.05, then filter cells through the strainer (20um) by a short spin down at 4C for 15 seconds. Wash the strainer once with 500ul PBB 0.05. Then pellet cells at 4C, 1000g for 3min. Aspirate the supernatant.
- Resuspend cells in 50ul PBB 0.05.
- Measure the cell concentration:
    - Take 1ul cell suspension, mix with 9ul DAPI working solution (1:10 dilution).

- Load onto the hemocytometer and count the cell number under the fluorescent microscope.

##### Distribute cells for Second Strand Synthesis

- Adjust cell concentration to 500 cells/ul using PBB 0.05.
- Distribute 4ul (2000 cells/well) to each well of multiple PCR strips.
- **Stop point:** store PCR strips in -80C if immediate processing is not needed. Post-ligation cells could be stored in -80C for 1 month.

##### Second Strand Synthesis

- Prepare second strand synthesis mix:

| Reagents | 1 well | For each strip |
|---|---|---|
| second strand synthesis buffer | 2/3 uL | 6 uL |
| second strand synthesis enzyme | 1/3 uL | 3 uL |
| Final Volume | 1 uL | 9 uL |

- Distribute 1 uL SSS reaction mix to each well of the strip. Gently vortex the strip for 1s, then briefly centrifuge using the benchtop centrifuge. Quickly put strips back on ice. (V=5uL)
- Perform second strand synthesis at 16C with shaking at 300rpm for 1 hour and 30 minutes.

##### 1x AMPURE beads purification

- Add 5ul nuclease-free water to each well, then add 10ul DNA binding buffer to each well, gently vortex for 1 second and then spin down. Incubate at room temperature for 5 minutes.
- Add 20ul AMPURE beads to each well, gently vortex and incubate at room temperature for 5 minutes.
- Put strips on the magnetic stand for 5min, then aspirate the supernatant carefully.
- Add 50uL 80% EtOH to each well while keeping the strips on the magnetic stand. Wait for 30 seconds, then aspirate the ethanol.
- Repeat the EtOH wash once more.
- Shortly spin down using the benchtop centrifuge, then put the strip back to the magnetic stand. Aspirate residual EtOH from the bottom of the strips.
- Resuspend beads of each well in 6.6uL of nuclease-free water.
- Incubate at room temperature for 5 minutes. Then put strips back to the magnetic stand for an extra 2 minutes.
- Transfer 6.6uL of the eluate from each well to new PCR strips. Put them back on ice for tagmentation reaction preparation.

##### Tagmentation

- Prepare the tagmentation mix on ice and add 6.6uL of the mix to each well.

| Reagents | For each strip |
|---|---|
| 2x tagmentation buffer | 60uL |
| Nextera-R2 Tn5 | 0.6uL |
| Final Volume | 60.6uL |

- Mix well by gentle vortexing for 1 second. After a brief spindown, incubate strips in the thermocycler at 55C for 5min.
- Prepare quenching buffer during the tagmentation:

| Reagents | For each strip |
|---|---|
| 1% SDS | 4uL |
| 20mg/ml BSA | 4uL |
| Final Volume | 8uL |

- After tagmentation, quickly put strips back onto the ice. Add 0.8uL of quenching buffer to each well. Mix by gentle vortexing for 1 second.

##### Multiplex PCR

- Add 1ul universal P5 primer (20uM) to each well.
- Incubate at 55C for 15 minutes.
- Add 2uL of 10% tween-20 to each well to quench SDS. Mix by gentle vortexing for 1 second and then briefly spinning down.
- Add 2uL of indexed EasySci-RNA P7 primer (with Nextera R2, 10uM) to each well for whole transcriptome indexing and amplification. Record the indices used in each well.
- Add 1uL of indexed sgRNA inner i7 primer (20uM) to each well of the strip for sgRNA library indexing and amplification. (Well 1-8 of each strip corresponds to the inner i7 barcode 1-8.)
- Add 20uL 2x NEBnext PCR master mix to each well. Mix by gentle vortexing for 1 second and a brief spindown. (V=40ul/reaction)
- Run the PCR program:

| Temperature | Time | Cycles |
|---|---|---|
| 72 °C | 5 minutes | |
| 98 °C | 30 seconds | |
| 98 °C | 10 seconds | 14 cycles |
| 65 °C | 30 seconds | 14 cycles |
| 72 °C | 30 seconds | 14 cycles |
| 72 °C | 2 minutes | |
| 4 °C | - | |

- After PCR, transfer PCR strips on ice.

##### sgRNA enrichment PCR

- Take 5uL multiplex PCR product from each well and combine the PCR product from 8 wells of each strip together (V=5ul/well*8 wells = 40ul/strip) respectively. Don't combine PCR products from different strips now since the indexing on sgRNA libraries hasn't completed yet.
- Purify each 40ul combined PCR product with double size selection using 0.8-1.2x AMPURE beads (Basically, mix 40ul combined PCR product with 32uL beads. Take the

supernatant, and mix it with another 16uL beads, and finally elute cDNA on beads). Elute combined cDNA of each strip in 16uL EB buffer respectively.

- For sgRNA enrichment PCR:
    - Add 2uL 10uM P5 universal primer + 2uL indexed EasySci-ATAC P7 primers (with TruSeq R2, 10uM) + 20uL 2x NEBnext PCR master mix to each well. Record the indices used in each well.
- Mix the reaction as stated above, then run the following PCR program:

| Temperature | Time | Cycles |
| --- | --- | --- |
| 98 °C | 1 min | |
| 98 °C | 10 seconds | 13 cycles |
| 66 °C | 30 seconds | 13 cycles |
| 72 °C | 20 seconds | 13 cycles |
| 72 °C | 2 minutes | |
| 4 °C | Hold | |

#### sgRNA library purification

- Combine PCR product of every 2 wells (contains sgRNA library of 2 strips, ~32000 cells, V=80ul).
- Perform the column purification using the Zymo DNA-concentrator5 kit. Elute each column in 20ul EB buffer.
- Load all eluate to the 2% e-gel, then perform the electrophoresis to check the sgRNA library amplification.
- If a strongest band at 276bp could be seen on the gel, perform gel extraction on this band. Finally elute sgRNA libraries from every two strips (~32000 cells/lib) in 20ul EB buffer.

#### Whole transcriptome library purification

- If the amplification of sgRNA libraries is successful, combine the rest of the PCR product.
- Take 200ul of combined PCR product, perform 0.8x AMPURE beads purification.
- Finally elute the whole transcriptome library containing cDNA from all single cells in the 20ul EB buffer.
- Dilute the library and check the size distribution using a 2% e-gel. The typical median size of the library should be ~350 bp.
- A typical gel image of the whole transcriptome library (lane 6) and the sgRNA library ready for gel extraction (lane 7):

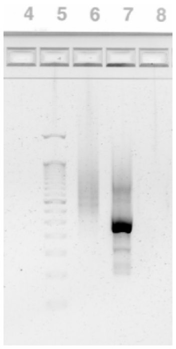


#### Concentration measurement and sequencing

- Measure the concentrations of libraries using Qubit.
- Dilute each library to 2nM for illumina NextSeq 1000 sequencing.
