## Supplementary file4 for "*PerturbSci-Kinetics*: Dissecting key regulators of transcriptome kinetics through scalable single-cell RNA profiling of pooled CRISPR screens": PE_bulkRNAseq_trim.sh

```
input_folder=$1
sample=$2
output_folder=$3

echo Trimming sample: $sample
trim_galore --paired $input_folder/$sample*.R1.fastq.gz $input_folder/$sample*.R2.fastq.gz -o $output_folder
echo Trimming $sample done.
```
