## Supplementary file4 for "*PerturbSci-Kinetics*: Dissecting key regulators of transcriptome kinetics through scalable single-cell RNA profiling of pooled CRISPR screens": SNP_calling_processing.sh

```
#! /bin/bash

#############################set up the directories and parameters####################################
bashrc_location=/asziraki/original_pipeline/sci-RNA-seq3_pipeline/.bashrc
# 'original_pipeline' conda environment is necessary to run the pipeline

# define the fastq folder including all fastq files for bg SNP calling 
fastq_folder="/zxu/ref_HEK_vcf/ref_fq"

# define the PCR group sample id for each fastq file
sample_ID="/zxu/ref_HEK_vcf/sampleID.txt"

# define the output folder
all_output_folder="/zxu/ref_HEK_vcf"

# define the core number for parallele processing
core=16 # for most steps
samtools_core=4 # for reads filtering and sorting

# define the location of index files for reads alignment with STAR
index="/zxu/gencode_ref_anno/hg38_STAR_index"
reference_fa="/zxu/gencode_ref_anno/GRCh38.primary_assembly.genome.fa"
varscan_zx="/zxu/tools/varscan/VarScan.v2.3.9.jar"
java_zx="/zxu/tools/openjdk_v11/jdk-11.0.2/bin/java"

# Define the location of the sub script folder
script_folder="/zxu/scripts/bulk_PERNAseq"

# define the location of the R script for multi-core processing
R_script=$script_folder/sci3_bash_input_ID_output_core.R

script_path=$script_folder
######################################################################################################

now=$(date)
echo "Current time : $now"

#################load the env
conda deactivate
source $bashrc_location
conda activate original_pipeline

################# Trimming the read2
echo
echo "Start trimming the read2 file..."
echo $(date)

raw_fastq=$fastq_folder
trimmed_fastq=$all_output_folder/trimmed_fastq
bash_script=$script_path/PE_bulkRNAseq_trim.sh

mkdir -p $trimmed_fastq

Rscript $R_script $bash_script $raw_fastq $sample_ID $trimmed_fastq $core

############align the reads with STAR, filter the reads based on q > 30, and remove duplicates based on UMI sequence and tagmentation site

#define the output folder for mapping
input_folder=$trimmed_fastq
STAR_output_folder=$all_output_folder/STAR_alignment
filtered_sam_folder=$all_output_folder/filtered_sam

#align read2 to the index file using STAR
echo "Start alignment using STAR..."
echo input folder: $input_folder
echo sample ID file: $sample_ID
echo index file: $index
echo output_folder: $STAR_output_folder

#make the output folder
mkdir -p $STAR_output_folder

#remove the index from the memory
STAR --genomeDir $index --genomeLoad Remove
#start the alignment
for sample in $(cat $sample_ID); do echo Aligning $sample;STAR --runThreadN $core --outSAMstrandField intronMotif --genomeDir $index --readFilesCommand zcat --readFilesIn $input_folder/$sample*R1.fastq.gz $input_folder/$sample*R2.fastq.gz --outFileNamePrefix $STAR_output_folder/$sample --genomeLoad LoadAndKeep --outFilterMultimapNmax 1; done
#remove the index from the memory
STAR --genomeDir $index --genomeLoad Remove
echo "All alignment done."

#Filter and sort the sam file
echo
echo "Start filter and sort the sam files..."
echo input folder: $STAR_output_folder
echo output folder: $filtered_sam_folder
mkdir -p $filtered_sam_folder

for sample in $(cat $sample_ID); do
samtools view -bh -q 30 -@ $core $STAR_output_folder/$sample*.sam|samtools sort -@ $core -|samtools view -bh -@ $core ->$filtered_sam_folder/$sample.bam
samtools index -@ $core $filtered_sam_folder/$sample.bam
done

# Call SNPs from the control files
### Generate a script with: (1) input folder (2) sample list (3) output folder, merge the single cell sam files in the
# input folder, and then call SNPs from the merged sequences
# first generate a tmp folder for all sorted singe cell bam files

sample_list=$sample_ID
input_folder=$filtered_sam_folder
output_folder=$all_output_folder/merged_bam_call_SNP

mkdir -p $output_folder

echo "Converting and merging bam files."
samtools merge -@ $core -f $output_folder/merged.no_sort.bam $input_folder/*.bam
samtools sort -@ $core $output_folder/merged.no_sort.bam > $output_folder/merged.bam

echo "Calling SNPs."
samtools mpileup -B -f $reference_fa $output_folder/merged.bam > $output_folder/output.mpileup
$java_zx -jar $varscan_zx mpileup2snp $output_folder/output.mpileup --strand-filter 0 > $output_folder/HEK293_ref_SNP.vcf

echo "analysis done, removing temp files...."
echo "All analysis done."

conda deactivate
```
