## Supplementary file4 for "*PerturbSci-Kinetics*: Dissecting key regulators of transcriptome kinetics through scalable single-cell RNA profiling of pooled CRISPR screens": key_functions.R

```
############load packages############
library(dplyr)
library(Matrix)
library(ggplot2)
library(rtracklayer)
library(clusterProfiler)
library(umap)
library(Seurat)
library(rhdf5)
library(viridis)
library(parallel)
library(rtracklayer)
library(GenomicRanges)

############load the R object and filter cells############
filter_dT_cells <- function(file_directory, condition_table_directory, UMI_threshold, gene_threshold, unmatched_rate_threshold){

  #load the object
  load(file_directory)
  df_cell$sample <- as.character(df_cell$sample)
  df_gene$gene_id <- as.character(df_gene$gene_id)
  rownames(gene_count_all) <- df_gene$gene_id
  colnames(gene_count_all) <- df_cell$sample
  
  #load condition table
  condition_table <- read.csv(condition_table_directory, header = T)
  
  #filter cells based on UMI-cutoff, gene-cutoff, and unmatched rate
  binarized_gene_count_all <- gene_count_all
  binarized_gene_count_all@x[binarized_gene_count_all@x > 0] <- 1
  genes_per_cell <- Matrix::colSums(binarized_gene_count_all)
  gene_based_whitelist <- names(genes_per_cell)[genes_per_cell > gene_threshold]
  UMI_based_whitelist <- df_cell[df_cell$UMI_count > UMI_threshold, "sample"]
  unmatched_rate_based_whitelist <- df_cell[df_cell$unmatched_rate < unmatched_rate_threshold, "sample"]
  
  whitelist <- intersect(gene_based_whitelist, intersect(UMI_based_whitelist, unmatched_rate_based_whitelist))
  
  whitelisted_df_cell <- df_cell[df_cell$sample %in% whitelist, ]
  whitelisted_gene_count_all <- gene_count_all[, whitelist]
  
  #split the cell names for recognition of different conditions
  separated_CB_df <- tidyr::separate(data = data.frame(full_name = whitelist), col = "full_name", into = c('PCR_group', 'RT_lig_CB'), sep = '\\.', remove = F)
  separated_CB_df$lig_cb <- substr(separated_CB_df$RT_lig_CB, start = 1, stop = 10)
  separated_CB_df$RT_cb <- substr(separated_CB_df$RT_lig_CB, start = 11, stop = 20)
  
  #separate cells in different groups
  UMI_counts <- c()
  conditions <- c()
  cell_names <- c()
  for (each_condition in unique(condition_table$Conditions)){
    available_RT_bc <- condition_table[condition_table$Conditions == each_condition, "barcode"]
    cells_in_this_condition <- separated_CB_df[separated_CB_df$RT_cb %in% available_RT_bc, "full_name"]
    UMI_counts <- append(UMI_counts, whitelisted_df_cell[whitelisted_df_cell$sample %in% cells_in_this_condition, "UMI_count"])
    conditions <- append(conditions, rep(each_condition, length(cells_in_this_condition)))
    cell_names <- append(cell_names, whitelisted_df_cell[whitelisted_df_cell$sample %in% cells_in_this_condition, "sample"])
  }
  UMI_per_group_df <- data.frame(cell_names = cell_names, UMI_counts = UMI_counts, conditions = conditions)
  p <- ggplot(UMI_per_group_df, aes(x=conditions, y=UMI_counts, fill=conditions))+geom_violin()+geom_boxplot(width=0.2)
  
  list(filtered_merged_matrix = whitelisted_gene_count_all,
       filtered_merged_cells_bc_tb = UMI_per_group_df,
       UMI_per_condition_plot = p)
}

############convert gene ids to gene symbols############
gene_id2gene_names <- function(input_gene_id_vector, gtf_dir){
  #load the gtf file
  gtf <- rtracklayer::import(gtf_dir, format = 'gtf')
  gtf <- gtf[gtf$type == "gene"]
  id_conversion_table <- data.frame(gene_id = gtf$gene_id, symbol = gtf$gene_name)
  remove(gtf)
  
  id_conversion_table[match(input_gene_id_vector, id_conversion_table$gene_id), "symbol"]
}

############load sgRNA expression matrix and reformat sgRNA cell names############
gRNA_cell_reformatting <- function(input_gRNA_summary_rdata_dir, reformatting_df){
  
  load(input_gRNA_summary_rdata_dir)
  input_gRNA_count_mat <- gRNA_count
  
  #the first step: unify cell name format between gRNA lib and whole txme lib
  gRNA_cell_names_df <- data.frame(gRNA_cell_names=colnames(input_gRNA_count_mat)) %>% tidyr::separate(col="gRNA_cell_names", into=c("gRNA_PCR_group", "bc"), sep="\\.", remove=FALSE)

  gRNA_cell_names_df <- dplyr::left_join(x = gRNA_cell_names_df, y = reformatting_df, by = "gRNA_PCR_group")
  new_gRNA_count_mat <- input_gRNA_count_mat[,gRNA_cell_names_df$gRNA_cell_names]

  new_cell_names <- paste(gRNA_cell_names_df$whole_txme_PCR_group, gRNA_cell_names_df$bc, sep = ".")
  colnames(new_gRNA_count_mat) <- new_cell_names
  new_gRNA_count_mat
}


############integrate whole tx cells with sgRNA cells############
match_whole_nascent_txme_with_gRNA <- function(whole_txme_obj, nascent_txme_obj, gRNA_expr_mat, max_gRNA_ratio_cutoff = 0.6, first_to_second_ratio=3, gtf_for_gene_name_conversion="/zxu/gencode_ref_anno/human_v38.primary_assembly.gtf"){
    
    matched_cells_name <- intersect(colnames(gRNA_expr_mat), whole_txme_obj$filtered_merged_cells_bc_tb$cell_names)
    matched_condition_table <- whole_txme_obj$filtered_merged_cells_bc_tb[match(matched_cells_name, whole_txme_obj$filtered_merged_cells_bc_tb$cell_names),]
    rownames(matched_condition_table) <- matched_condition_table$cell_names
    
    matched_whole_txme_expr_mat <- whole_txme_obj$filtered_merged_matrix[,matched_condition_table$cell_names]
    matched_nascent_txme_expr_mat <- nascent_txme_obj$filtered_merged_matrix[,matched_condition_table$cell_names]
    matched_gRNA_expr_mat <- gRNA_expr_mat[,matched_condition_table$cell_names]
    
    ###change the gene id of whole txme to gene names
    gene_names <- gene_id2gene_names(rownames(matched_whole_txme_expr_mat), gtf_dir = gtf_for_gene_name_conversion)
    matched_whole_txme_expr_mat <- matched_whole_txme_expr_mat[(!is.na(gene_names))&(!duplicated(gene_names)),]
    gene_names <- gene_names[(!is.na(gene_names))&(!duplicated(gene_names))]
    rownames(matched_whole_txme_expr_mat) <- gene_names
    
    ###change the gene id of nascent txme to gene names
    gene_names <- gene_id2gene_names(rownames(matched_nascent_txme_expr_mat), gtf_dir = gtf_for_gene_name_conversion)
    matched_nascent_txme_expr_mat <- matched_nascent_txme_expr_mat[(!is.na(gene_names))&(!duplicated(gene_names)),]
    gene_names <- gene_names[(!is.na(gene_names))&(!duplicated(gene_names))]
    rownames(matched_nascent_txme_expr_mat) <- gene_names
    
    ###calculate nascent UMI counts and ratio of single cells
    matched_condition_table$nascent_UMI_counts <- 0
    matched_condition_table$nascent_ratio <- 0
    matched_condition_table[colnames(matched_nascent_txme_expr_mat), "nascent_UMI_counts"] <- colSums(matched_nascent_txme_expr_mat)
    matched_condition_table[colnames(matched_nascent_txme_expr_mat), "nascent_ratio"] <- matched_condition_table$nascent_UMI_counts/matched_condition_table$UMI_counts
    
    ###identify gRNA target of single cells
    matched_condition_table$target <- "mixed"
    matched_condition_table$target_genes <- "mixed"
    matched_condition_table$binarized_target_genes <- "mixed"
    
    cells_max_gRNA <- apply(X = matched_gRNA_expr_mat, MARGIN = 2, FUN = max)
    cells_gRNA_sum <- colSums(matched_gRNA_expr_mat)
    cells_gRNA_ratio <- cells_max_gRNA/cells_gRNA_sum
    
    #add gRNA UMI counts to the condition table
    matched_condition_table$gRNA_UMI_counts <- 0
    matched_condition_table[colnames(matched_gRNA_expr_mat), "gRNA_UMI_counts"] <- cells_gRNA_sum
    
    #identify gRNA-based singlets
    ###the first layer of filtering based on the ratio of the most abundant gRNA
    singlets_gRNA_expr_mat <- matched_gRNA_expr_mat[,cells_gRNA_ratio >= max_gRNA_ratio_cutoff]
    singlets_cell_names <- colnames(singlets_gRNA_expr_mat)
    all_targets <- rownames(matched_gRNA_expr_mat)
    targets <- sapply(singlets_cell_names, function(x){
        gRNA_sorted <- sort(singlets_gRNA_expr_mat[,x], decreasing = T)
        first_gRNA <- gRNA_sorted[1]
        second_gRNA <- gRNA_sorted[2]
        
        ###the second layer of filtering based on the relative ratio between top2 gRNAs
        if(first_gRNA >= second_gRNA*first_to_second_ratio){
            return(names(first_gRNA))
        }else{
            return("mixed")
        }
    })
    target_genes <- sapply(strsplit(targets, split = "_"), function(x) x[[1]])
    
    #add target gRNA and gene names of singlets to the condition table
    matched_condition_table[singlets_cell_names, "target"] <- targets
    matched_condition_table[singlets_cell_names, "target_genes"] <- target_genes
    matched_condition_table[matched_condition_table$target_genes != "mixed", "binarized_target_genes"] <- "singlets"                       
    
    ###return a new obj containing all info
    list(matched_condition_table=matched_condition_table, 
         matched_whole_txme_expr_mat=matched_whole_txme_expr_mat,
         matched_nascent_txme_expr_mat=matched_nascent_txme_expr_mat,
         matched_gRNA_expr_mat=matched_gRNA_expr_mat)                       
}


############rates calculation#############
synth_deg_bootstrapping_NTC_vs_KD <- function(matched_obj, NTC_population_name, KD_population_name, resampling_time=500, target_gene_colname = "target_genes", genes_0_percentage_filtering_threshold = 0.1, random_seed = 10, significance_p_cutoff = 0.05){
    
    set.seed(random_seed)
    
    ###get cell names of KD population
    KD_cell_names <- matched_obj$matched_condition_table[matched_obj$matched_condition_table[,target_gene_colname] == KD_population_name, "cell_names"]
    KD_cell_num <- length(KD_cell_names)
    
    ###get cell names of NTC
    NTC_cell_names <- matched_obj$matched_condition_table[matched_obj$matched_condition_table[,target_gene_colname] == NTC_population_name, "cell_names"]
    NTC_cell_num <- length(NTC_cell_names)
    
    ###define several functions to calculate synth and deg rates
    normalization_for_rates_calc <- function(sampled_pseudobulk_whole_txme, sampled_pseudobulk_nascent_txme){
        norm_pseudobulk_whole_cpm <- sampled_pseudobulk_whole_txme*1e6/sum(sampled_pseudobulk_whole_txme)
        norm_pseudobulk_nascent_cpm <- sampled_pseudobulk_nascent_txme*1e6/sum(sampled_pseudobulk_whole_txme)
        norm_pseudobulk_preexisted_cpm <- norm_pseudobulk_whole_cpm - norm_pseudobulk_nascent_cpm
        
        list(norm_pseudobulk_whole_cpm=norm_pseudobulk_whole_cpm, 
             norm_pseudobulk_nascent_cpm=norm_pseudobulk_nascent_cpm, 
             norm_pseudobulk_preexisted_cpm=norm_pseudobulk_preexisted_cpm)
    }
    
    synth_deg_calculation <- function(norm_cpm_list){
        
        ###for genes w/o preexisting counts, it's impossible to infer the deg rate
        non_zero_old_genes <- names(norm_cpm_list$norm_pseudobulk_preexisted_cpm)[norm_cpm_list$norm_pseudobulk_preexisted_cpm > 0]
        deg_rate <- -log(norm_cpm_list$norm_pseudobulk_preexisted_cpm[non_zero_old_genes]/norm_cpm_list$norm_pseudobulk_whole_cpm[non_zero_old_genes])/2
        synth_rate <- norm_cpm_list$norm_pseudobulk_whole_cpm[non_zero_old_genes] * deg_rate
        
        ###for synth rate, we can assume it's 0 if we can't get any count from the nascent part
        synth_mat <- Matrix::Matrix(data = 0, nrow = length(norm_cpm_list$norm_pseudobulk_whole_cpm), ncol = 1, sparse = T)
        synth_mat[match(non_zero_old_genes, names(norm_cpm_list$norm_pseudobulk_whole_cpm)), 1] <- synth_rate
        
        ###for deg rate, the 0 means invalid
        deg_mat <- Matrix::Matrix(data = 0, nrow = length(norm_cpm_list$norm_pseudobulk_whole_cpm), ncol = 1, sparse = T)
        deg_mat[match(non_zero_old_genes, names(norm_cpm_list$norm_pseudobulk_whole_cpm)), 1] <- deg_rate
        
        list(synth_mat=synth_mat, deg_mat=deg_mat)
    }
    
    ###resample equal number of NTC cells with KD cells for multiple times and form the background distribution
    NTC_target_combined_synth_deg_resampling_list <- lapply(1:resampling_time, function(x){
        ###in this function, we assume the cell number of the KD population is much smaller than NTC
        sampled_NTC_target_combined_cell_names <- sample(x = c(NTC_cell_names, KD_cell_names), size = KD_cell_num, replace = F)
        sampled_pseudobulk_whole_txme <- Matrix::rowSums(matched_obj$matched_whole_txme_expr_mat[,sampled_NTC_target_combined_cell_names])
        sampled_pseudobulk_nascent_txme <- Matrix::rowSums(matched_obj$matched_nascent_txme_expr_mat[,sampled_NTC_target_combined_cell_names])
        
        ###1. calculate the normalized pseudobulk expression
        norm_sampled_NTC_target_combined_txme_list <- normalization_for_rates_calc(sampled_pseudobulk_whole_txme, sampled_pseudobulk_nascent_txme)
        
        ###2. calculate rates and return two matrices
        sampled_rates_matrices_list <- synth_deg_calculation(norm_sampled_NTC_target_combined_txme_list)
        sampled_rates_matrices_list
    })
    
    ###merge synth and deg matrices separately
    resampled_NTC_target_combined_synth_mat_list <- lapply(NTC_target_combined_synth_deg_resampling_list, function(x) x[["synth_mat"]])
    resampled_NTC_target_combined_deg_mat_list <- lapply(NTC_target_combined_synth_deg_resampling_list, function(x) x[["deg_mat"]])
    
    resampled_NTC_target_combined_synth_mat <- do.call(cbind, resampled_NTC_target_combined_synth_mat_list)                                
    resampled_NTC_target_combined_deg_mat <- do.call(cbind, resampled_NTC_target_combined_deg_mat_list)
                                         
    rownames(resampled_NTC_target_combined_synth_mat) <- rownames(matched_obj$matched_whole_txme_expr_mat)
    rownames(resampled_NTC_target_combined_deg_mat) <- rownames(matched_obj$matched_whole_txme_expr_mat)
                                         
    ###choose genes with confident rates calculation                                     
    binarized_resampled_NTC_target_combined_deg_mat <- resampled_NTC_target_combined_deg_mat                                   
    binarized_resampled_NTC_target_combined_deg_mat@x[binarized_resampled_NTC_target_combined_deg_mat@x > 0] <- 1
    zero_percentage <- Matrix::rowSums(binarized_resampled_NTC_target_combined_deg_mat)/resampling_time                                    
    confident_genes_in_background <- rownames(binarized_resampled_NTC_target_combined_deg_mat)[zero_percentage >= (1-genes_0_percentage_filtering_threshold)]                                                                        
                                         
    ###calculate rates of KD cells
    KD_pseudobulk_whole_txme <- Matrix::rowSums(matched_obj$matched_whole_txme_expr_mat[,KD_cell_names])
    KD_pseudobulk_nascent_txme <- Matrix::rowSums(matched_obj$matched_nascent_txme_expr_mat[,KD_cell_names])
    
    norm_KD_txme_list <- normalization_for_rates_calc(KD_pseudobulk_whole_txme, KD_pseudobulk_nascent_txme)
    KD_rates_list <- synth_deg_calculation(norm_KD_txme_list)
    rownames(KD_rates_list[["synth_mat"]]) <- rownames(matched_obj$matched_whole_txme_expr_mat)
    rownames(KD_rates_list[["deg_mat"]]) <- rownames(matched_obj$matched_whole_txme_expr_mat)                                     
    
    ###choose confident genes of KD cells (deg rate > 0)
    confident_KD_genes <- rownames(KD_rates_list$deg_mat)[KD_rates_list$deg_mat[,1] > 0]                                     

    ###export the matrices and we can check genes of interest on ourselves
    ###merge confident genes from NTC and KD together to capture on-and-off regulation
    genes_to_export <- unique(c(confident_KD_genes, confident_genes_in_background))                                     
    output_list <- list(KD_synth_vector = KD_rates_list[["synth_mat"]][genes_to_export,], KD_deg_vector = KD_rates_list[["deg_mat"]][genes_to_export,],
                         resampled_bg_synth_mat = resampled_NTC_target_combined_synth_mat[genes_to_export,], resampled_bg_deg_mat = resampled_NTC_target_combined_deg_mat[genes_to_export,])
    
    ###examine the significance of each gene
    ##synthesis
    syn_pval_list <- lapply(genes_to_export, function(x){
        bg_syn_mean <- mean(output_list$resampled_bg_synth_mat[x,])
        gene_x_syn <- output_list$KD_synth_vector[x]
        if(gene_x_syn <= bg_syn_mean){
            two_sided_p <- (sum(output_list$resampled_bg_synth_mat[x,] <= gene_x_syn)+sum(output_list$resampled_bg_synth_mat[x,] >= 2*bg_syn_mean-gene_x_syn))/resampling_time
            export_df <- data.frame(synth.two.sided.p = two_sided_p, sig_synth_direction="Down")
        }else{
            two_sided_p <- (sum(output_list$resampled_bg_synth_mat[x,] >= gene_x_syn)+sum(output_list$resampled_bg_synth_mat[x,] <= 2*bg_syn_mean-gene_x_syn))/resampling_time
            export_df <- data.frame(synth.two.sided.p = two_sided_p, sig_synth_direction="Up")
        }
        
        ##modify the direction if the pval is not significant
        if(export_df$synth.two.sided.p > significance_p_cutoff){
            export_df$sig_synth_direction <- "No"
        }
        export_df
    })
    syn_pval_df <- as.data.frame(do.call(rbind, syn_pval_list))                                  
                                         
    ##degradation                                     
    deg_pval_list <- lapply(genes_to_export, function(x){
        bg_deg_mean <- mean(output_list$resampled_bg_deg_mat[x,])
        gene_x_deg <- output_list$KD_deg_vector[x]
        if(gene_x_deg <= bg_deg_mean){
            two_sided_p <- (sum(output_list$resampled_bg_deg_mat[x,] <= gene_x_deg)+sum(output_list$resampled_bg_deg_mat[x,] >= 2*bg_deg_mean-gene_x_deg))/resampling_time
            export_df <- data.frame(deg.two.sided.p = two_sided_p, sig_deg_direction="Down")
        }else{
            two_sided_p <- (sum(output_list$resampled_bg_deg_mat[x,] >= gene_x_deg)+sum(output_list$resampled_bg_deg_mat[x,] <= 2*bg_deg_mean-gene_x_deg))/resampling_time
            export_df <- data.frame(deg.two.sided.p = two_sided_p, sig_deg_direction="Up")
        }
        
        ##modify the direction if the pval is not significant
        if(export_df$deg.two.sided.p > significance_p_cutoff){
            export_df$sig_deg_direction <- "No"
        }
        export_df
    })                                                     
    deg_pval_df <- as.data.frame(do.call(rbind, deg_pval_list))
                                         
    ###integrate data into a final dataframe and finally export
    info_df <- data.frame(synth_rate = output_list$KD_synth_vector, deg_rate = output_list$KD_deg_vector, 
                          synth_pval = syn_pval_df$synth.two.sided.p, synth_direction = syn_pval_df$sig_synth_direction, 
                          deg_pval = deg_pval_df$deg.two.sided.p, deg_direction = deg_pval_df$sig_deg_direction)                                     
    output_list[["info_df"]] <- info_df
    output_list                                     
}
```
