## Supplementary file4 for "*PerturbSci-Kinetics*: Dissecting key regulators of transcriptome kinetics through scalable single-cell RNA profiling of pooled CRISPR screens": README.md

# PerturbSci_Kinetics
Reads processing scripts for PerturbSci-Kinetics.
The bioRxiv preprint: https://doi.org/10.1101/2023.01.29.526143
___
### **Background SNP calling (/bg_SNP_calling/SNP_calling_processing.sh)**
#### Key parameters
* Fastq input: Paired-end full-coverage bulk RNA-seq.
* Sample ID: a text file containing the prefix of each sample on each line. R1 and R2 files from the sample should share the same prefix.
* Reference fasta: the fasta file of the reference genome. It is used during reads pileup.
* Index: the STAR index of the reference genome.
* Output folder: the directory for all output files.
* Script folder: the folder for all sub scripts.
* Other parameters include core number, the directories of packages.

#### Steps
1. Trim adapter sequences by automatic detection.
2. STAR alignment.
3. Filter aligned reads.
4. Merge bams and sort the merged bam.
5. Summarize the base identities of reads mapped to each genomic location. 
6. Inherent SNP calling.

#### Key output
* A vcf file containing background mutations in RNA.
___
### **Single cell whole/nascent transcriptomes reprocessing steps (/whole_tx_processing/Main_processing.sh)**
#### Key parameters
* Fastq input: Paired-end PerturbSci-Kinetics demultiplexed whole transcriptome fastq files. 
* Sample ID: a text file containing the prefix of each sample on each line. R1 and R2 files from the sample should share the same prefix.
* Reference fasta: the fasta file of the reference genome. 
* Index: the STAR index of the reference genome.
* Gtf file: the annotation file for the matched reference genome. It is used in feature counting.
* Reference SNP file: the SNP vcf file generated from the script above. It is used to filter out inherent mutations in the RNA.
* Output folder: the directory for all output files.
* Script folder: the folder for all sub scripts.
* Cutoff: only cell barcodes with reads number > this cutoff will be considered for further processed.
* Custom barcode folder: the folder for all barcodes. 
* RT barcode, ligation barcode: pickle files containing all valid barcode sequences with at most 1 mismatch.
* Barcodes: the text file containing all RT+ligation barcode combinations.
* Other parameters include core number, the directories of packages.

#### Steps
1. Change file names of fastq to make them callable in the following steps.
2. Attach UMI sequences on R1 to headers of R2.
3. Trim potential polyA sequences from the 3'end of R2.
4. STAR alignment.
5. Filter aligned reads.
6. PCR duplicates removal based on both mapped genomic coordinates and UMI.
7. Single-cell sam files generation.
8. Transform the alignment information in single-cell sams to tables at the single-base level.
9. Identify T>C mutations on each single read and extract read names of nascent reads.
10. Extract nascent reads from single-cell sams.
11. Gene-level feature counting on both single-cell whole/nascent sams and re-format the single-cell gene expression matrix.

#### Key output
* Two R object files containing single-cell whole/nascent tx expression count matrix respectively.
___
### **Single cell sgRNA reprocessing steps (/sgRNA_reads_processing/sgRNA_processing.sh)**
#### Key parameters
* Fastq input: Paired-end PerturbSci-Kinetics demultiplexed sgRNA fastq files. 
* Cutoff: only cell barcodes with the number of sgRNA UMI > cutoff will be considered
* SgRNA correction file: pickle files containing all valid sgRNA sequences with at most 1 mismatch.
* SgRNA annotation df: A txt file including the sgRNA names, and corresponding gene symbols. It is used during the expression matrix construction.
* Other parameters are roughly the same as above.

#### Steps
1. Change file names of fastq to make them callable in the following steps.
2. One-step sgRNA identification, de-duplication, and counting.
3. Re-format the single-cell sgRNA expression matrix.

#### Key output
* An R object file containing an single-cell sgRNA expression matrix.
___
### **Paired-end bulk SLAM-seq reads reprocessing steps (/SLAMseq_processing/SLAM_seq_main_processing.sh)**
#### Key parameters
* Parameters are roughly the same as those in single-cell processing scripts.

#### Steps
1. Change file names of fastq to make them callable in the following steps.
2. Attach UMI sequences on R1 to headers of R2.
3. Trim potential adapter sequences from the 3'end of R1 and R2.
4. STAR alignment.
5. Filter aligned reads.
6. PCR duplicates removal by picard.
7. Transform the alignment information in sams to tables at the single-base level.
8. Split the alignment info table into small sub tables.
9. Identify T>C mutations on each read pair.
10. Merge mutation info identified from all sub tables under one sample, and extract names of nascent reads.
11. Extract nascent reads from sams.
12. Gene-level feature counting on both whole/nascent bams and re-format the gene expression matrix.

#### Key output
* Two R object files containing gene x sample whole/nascent tx expression count matrix respectively.
___
### **Key R functions in downstream analysis (/downstream_functions/key_functions.R)**
1. filter_dT_cells(): Get single-cell whole tx expression matrix from the output R.object of the preprocessing script.
2. gene_id2gene_names(): Convert gene ids to gene symbols using the matched gtf file
3. gRNA_cell_reformatting(): Read and reformat the sgRNA single-cell expression matrix to make it compatible with the integradation with whole tx info.
4. match_whole_nascent_txme_with_gRNA(): Integrate whole tx data with sgRNA info, identify sgRNA-based singlets, and return a integrated obj.
5. synth_deg_bootstrapping_NTC_vs_KD(): Calculate synthesis and degradation rates on cell populations. Also perform permutation tests between perturbations and NTC to examine the statistical significance.
