## Supplementary file4 for "*PerturbSci-Kinetics*: Dissecting key regulators of transcriptome kinetics through scalable single-cell RNA profiling of pooled CRISPR screens": gRNA_MM_formatting.R

```
args = commandArgs(trailingOnly=TRUE)

# define the report folder from sci-RNA-seq pipeline
report_folder = args[1]

# define the output folder for output the df_cell, df_gene and gene_count matrix
output_folder = args[1]

suppressMessages(library(Matrix))
suppressMessages(library(tidyverse))
suppressMessages(library(data.table))


sciRNAseq_gene_count_summary <- function (gene_count_folder) {
  gene_matrix_dir = paste(gene_count_folder, "/gRNA_mat.count", sep = "")
  df_gene_dir = paste(gene_count_folder, "/gRNA_annotat.report", sep = "")
  df_cell_dir = paste(gene_count_folder, "/cell_gRNA_annotat.report", sep = "")

  df_gene = read.csv(df_gene_dir, header = F)
  df_cell = read.csv(df_cell_dir, header = F)
  gene_matrix = read.csv(gene_matrix_dir, header = F)
  colnames(df_gene) = c("gRNA_index", "gRNA_name")
  colnames(df_cell) = c("cell_index", "cell_name")
  colnames(gene_matrix) = c("gRNA_index", "cell_index", "count")
  rownames(df_gene) = df_gene$gene_id
  rownames(df_cell) = df_cell$cell_name
  gene_count = sparseMatrix(i = gene_matrix$gRNA_index, j = gene_matrix$cell_index, x = gene_matrix$count)
  df_gene = df_gene[1:nrow(gene_count), ]
  rownames(gene_count) = df_gene$gRNA_name
  colnames(gene_count) = df_cell$cell_name

  return(list(df_cell, df_gene, gene_count))
}
result = sciRNAseq_gene_count_summary(report_folder)
gRNA_df_cell = result[[1]]
gRNA_df_gene = result[[2]]
gRNA_count = result[[3]]
save(gRNA_df_cell, gRNA_df_gene, gRNA_count, file = paste0(output_folder, "/Sci3_gRNA_Summary.RData"))
```
