## Supplementary file4 for "*PerturbSci-Kinetics*: Dissecting key regulators of transcriptome kinetics through scalable single-cell RNA profiling of pooled CRISPR screens": gRNA_sc_counting.py

```
import subprocess
import sys
import os
import gzip
from multiprocessing import Pool
from functools import partial
import pickle
import pandas as pd
import numpy as np

###This script is used to retrieve single cell identities of each gRNA reads and get UMI number of each gRNA tx


def hamming_dist(seq1, seq2):
    return np.count_nonzero(np.array(list(seq1)) != np.array(list(seq2)))


def gRNA_cell_UMI_ident(sample, input_folder, RT_barcode_list, inner_i7_bc_list,\
                        ligation_barcode_list, gRNA_correction_list, mismatch_rate = 1):

    # sample = 'sciATAC3_EXP10_01'
    Read1 = input_folder + "/" + sample + ".R1.fastq.gz"
    Read2 = input_folder + "/" + sample + ".R3.fastq.gz"
    Read3 = input_folder + "/" + sample + ".R2.fastq.gz"

    f1 = gzip.open(Read1)
    f2 = gzip.open(Read2)
    I5_reads = gzip.open(Read3)

    line1 = f1.readline()
    line2 = f2.readline()
    line3 = I5_reads.readline()
    total_line = 0
    filtered_line = 0
    
    UMI_per_cell_dict = dict()
    
    #generate all potential constant region1 and 2 sequences having distance <= 1 with the original sequence.
    #compared with distance calculation, direct sequence matching could be much faster
    ori_constant1 = "CAAGTTGATA"
    constant_mismatch_list1 = set()
    for each_pos in range(len(ori_constant1)):
        ori_list = list(ori_constant1)
        for each_letter in ["A", "G", "C", "T", "N"]:
            ori_list[each_pos] = each_letter
            constant_mismatch_list1.add("".join(ori_list))
    constant_mismatch_list1 = list(constant_mismatch_list1)

    ori_constant2 = "ATCTTGTGGA"
    constant_mismatch_list2 = set()
    for each_pos in range(len(ori_constant2)):
        ori_list = list(ori_constant2)
        for each_letter in ["A", "G", "C", "T", "N"]:
            ori_list[each_pos] = each_letter
            constant_mismatch_list2.add("".join(ori_list))
    constant_mismatch_list2 = list(constant_mismatch_list2)
    
    while (line1):
        total_line += 1
        
        if total_line % 1000000 == 0:
            print("%s Read pairs have been processed!" % total_line)
        
        line1_header = line1
        line2_header = line2
        line1 = f1.readline()
        line2 = f2.readline()
        line3 = I5_reads.readline()
            #print("read1: ", line1)
            # first check if the ligation barcode match with the barcode
        tmp_lig = line3[0:10].decode()
        
        constant_R1 = line1[18:28].decode()
        constant_R2 = line2[10:20].decode()
        
        #use constant regions to filter pattern-matched reads and match i5 ligation barcodes
        if tmp_lig in ligation_barcode_list and constant_R1 in constant_mismatch_list1 \
        and constant_R2 in constant_mismatch_list2:
            #print("constant + lig passed")
            ligation_bc_match = ligation_barcode_list[tmp_lig]
            inner_i7 = line2[0:10].decode()
            #match inner i7 barcodes
            if inner_i7 in inner_i7_bc_list:
                #print("inner i7 passed")
                inner_i7_match = inner_i7_bc_list[inner_i7]
                RT_bc = line1[8:18].decode()
                gRNA_region = line2[35:55].decode()
                #match RT barcode and gRNA sequences
                if RT_bc in RT_barcode_list and gRNA_region in gRNA_correction_list:
                    #print("CB and gRNA passed")
                    RT_bc_match = RT_barcode_list[RT_bc]
                    gRNA_match = gRNA_correction_list[gRNA_region]
                    filtered_line += 1
                    cell_id = sample + inner_i7_match + "." + ligation_bc_match + RT_bc_match
                    UMI = line1[0:8].decode()
                    
                    #start UMI identification
                    if cell_id not in UMI_per_cell_dict:
                        UMI_per_cell_dict[cell_id] = dict()
                        UMI_per_cell_dict[cell_id][gRNA_match] = set([UMI])
                    else:
                        if gRNA_match not in UMI_per_cell_dict[cell_id]:
                            UMI_per_cell_dict[cell_id][gRNA_match] = set([UMI])
                        else:
                            UMI_per_cell_dict[cell_id][gRNA_match].add(UMI)
                    
                    line1 = f1.readline()
                    line1 = f1.readline()
                    line1 = f1.readline()
                    line2 = f2.readline()
                    line2 = f2.readline()
                    line2 = f2.readline()
                else:
                    line1 = f1.readline()
                    line1 = f1.readline()
                    line1 = f1.readline()
                    line2 = f2.readline()
                    line2 = f2.readline()
                    line2 = f2.readline()
                    
            else:
                line1 = f1.readline()
                line1 = f1.readline()
                line1 = f1.readline()
                line2 = f2.readline()
                line2 = f2.readline()
                line2 = f2.readline()
        else:      
            line1 = f1.readline()
            line1 = f1.readline()
            line1 = f1.readline()
            line2 = f2.readline()
            line2 = f2.readline()
            line2 = f2.readline()
        line3 = I5_reads.readline() 
        line3 = I5_reads.readline()
        line3 = I5_reads.readline()
    f1.close()
    f2.close()
    I5_reads.close()
    
    print("sample name: %s, total line: %f, filtered line: %f, filter rate: %f" \
            %(sample, total_line, filtered_line, float(filtered_line) / float(total_line)))
    #print("Start counting UMI on sample: %s" % (sample))
    
    return UMI_per_cell_dict
    

def sc_gRNA_counting(gRNA_annotation_df, all_sample_UMI_per_cell_dict, output_folder, min_UMI_threshold):  
    
    if not os.path.exists(output_folder):
        os.makedirs(output_folder)
    
    gRNA_annotat = open(output_folder + "/gRNA_annotat.report", "w")
    cell_annotat = open(output_folder + "/cell_gRNA_annotat.report", "w")
    output_sparse_mat = open(output_folder + "/gRNA_mat.count", "w")
    
    gRNA_gene_df = pd.read_csv(gRNA_annotation_df, header = 0, sep = "\t")
    n_gRNA = gRNA_gene_df.shape[0]
    gRNA_index_dict = dict(zip(list(gRNA_gene_df["gRNA_seq"]), range(1, n_gRNA+1)))
    
    for each_gRNA in range(n_gRNA):
        gRNA_annotat.write(",".join([str(each_gRNA+1), gRNA_gene_df["names"][each_gRNA]]) + "\n")
    gRNA_annotat.close()
    
    #initiate a cell index
    n_cell = 1
    for cells, gRNA_dict in all_sample_UMI_per_cell_dict.items():
        total_UMI_per_cell = 0
        for UMI in gRNA_dict.values():
            total_UMI_per_cell += len(list(UMI))
        if total_UMI_per_cell >= min_UMI_threshold:
            #only cells with gRNA UMI > min thresholds will be kept for annotation
            cell_annotat.write(",".join([str(n_cell), cells]) + "\n")
            
            for gRNA, UMI in gRNA_dict.items():
                gRNA_index = gRNA_index_dict[gRNA]
                nUMI = len(list(UMI))
                #print([cells, gRNA_index, nUMI])
                output_sparse_mat.write(",".join([str(gRNA_index), str(n_cell), str(nUMI)]) + "\n")
            
            n_cell += 1
            
    gRNA_annotat.close()
    cell_annotat.close()
    output_sparse_mat.close()
            

def gRNA_UMI_ident_parallel(input_folder, sampleID, output_folder, RT_barcode_file, \
                            inner_i7_bc_file, ligation_barcode_file, gRNA_correction_file, \
                            gRNA_annotation_df, min_UMI_threshold, core):
    
    init_message = '''
    --------------------------start attaching UMI-----------------------------
    input folder: %s
    sample ID: %s
    output_folder: %s
    RT barcode file: %s
    ligation barcode file: %s
    gRNA barcode correction file: %s
    inner i7 index barcode file: %s
    ___________________________________________________________________________
    ''' %(input_folder, sampleID, output_folder, RT_barcode_file, ligation_barcode_file,\
          gRNA_correction_file, inner_i7_bc_file)
    
    print(init_message)
    
    
    print("Loading barcode dictionaries...")
    
    # generate the N5 RT barcode list:
    # barcodes_N5 = open(RT_barcode_file_N5, "rb")
    # with barcodes_N5 as f:
    #     RT_barcode_list_N5 = f.read().splitlines()
    # barcodes_N5.close()

    # # generate the N7 RT barcode list:
    # barcodes_N7 = open(RT_barcode_file_N7, "rb")
    # with barcodes_N7 as f:
    #     RT_barcode_list_N7 = f.read().splitlines()
    # barcodes_N7.close()

    barcodes = open(RT_barcode_file, "rb")
    RT_barcode_list = pickle.load(barcodes)
    barcodes.close()

    barcodes = open(ligation_barcode_file, "rb")
    ligation_barcode_list = pickle.load(barcodes)
    barcodes.close()
    
    barcodes = open(gRNA_correction_file, "rb")
    gRNA_correction_list = pickle.load(barcodes)
    barcodes.close()
    
    barcodes = open(inner_i7_bc_file, "rb")
    inner_i7_bc_list = pickle.load(barcodes)
    barcodes.close()
    
    print("Start loading sample names...")

    #for each sample in the sample list, use the read1 file, read2 file, output file
    # and barcode_list to run UMI_attach_read2_barcode_list
    sample_file = open(sampleID)
    sample_list = []
    for line in sample_file:
        sample = line.strip()
        sample_list.append(sample)
    sample_file.close()
    
    print("Start searching for cell identities and UMI ...")
    # parallele for the functions
    p = Pool(processes = int(core))
    #print("Processing core number: ", core_number)
    #gRNA_cell_UMI_ident(sample, input_folder, output_folder, RT_barcode_list, inner_i7_bc_list, ligation_barcode_list, gRNA_correction_list, mismatch_rate = 1)
    func = partial(gRNA_cell_UMI_ident, input_folder = input_folder, RT_barcode_list=RT_barcode_list, 
                   inner_i7_bc_list=inner_i7_bc_list, ligation_barcode_list=ligation_barcode_list, 
                   gRNA_correction_list=gRNA_correction_list, mismatch_rate = 1)
    #sciRNAseq_count(sample, input_folder, exons, genes, gene_end)
    result = p.map(func, sample_list)
    p.close()
    p.join()
    
    #combine all dictionaries together
    combined_dict = {}
    if len(result) == 1:
        combined_dict = result[0]
    elif len(result) == 2:
        combined_dict = {**result[0], **result[1]}
    else:
        combined_dict = {**result[0], **result[1]}
        for each_dict in range(2, len(result)):
            combined_dict = {**combined_dict, **result[each_dict]}
    
    print("Start counting UMI ...")
    #UMI counting and output
    mat_output_folder=output_folder + "/gRNA_report"
    sc_gRNA_counting(gRNA_annotation_df=gRNA_annotation_df, all_sample_UMI_per_cell_dict=combined_dict, 
                     output_folder=mat_output_folder, min_UMI_threshold=min_UMI_threshold)
    
    
    #print the completion message
    com_message = '''All done!'''
    print(com_message)


if __name__ == "__main__":
    input_folder = sys.argv[1]
    sampleID = sys.argv[2]
    output_folder = sys.argv[3]
    RT_barcode_file = sys.argv[4]
    inner_i7_bc_file = sys.argv[5]
    ligation_barcode_file = sys.argv[6]
    gRNA_correction_file = sys.argv[7]
    gRNA_annotation_df = sys.argv[8]
    min_UMI_threshold = int(sys.argv[9])
    core = int(sys.argv[10])
    gRNA_UMI_ident_parallel(input_folder, sampleID, output_folder, RT_barcode_file, 
                            inner_i7_bc_file, ligation_barcode_file, gRNA_correction_file, 
                            gRNA_annotation_df, min_UMI_threshold, core)
```
