## Supplementary file4 for "*PerturbSci-Kinetics*: Dissecting key regulators of transcriptome kinetics through scalable single-cell RNA profiling of pooled CRISPR screens": sgRNA_processing.sh

```
#! /bin/bash

########################### set up the directories ############################
#####set up directories and parameters
# define the fastq folder including all fastq files
fastq_folder=""

# define the PCR group sample id for each fastq file
sample_ID="/example_files/gRNA_sampleID.txt"

#define the scripts for single cell splitting and UMI recording
main_script="/main_scripts/gRNA_sc_counting.py"
reformatting_script="/main_scripts/gRNA_MM_formatting.R"

# define the output folder
all_output_folder=""

# define the core number for parallele processing
core=4

# define the number of gRNA UMI cutoff for keeping cells
cutoff=10

#define the folder of RT barcode dictionary
RT_barcode_file="/barcode_files/RT_bc_210923.pickle2"

#define the folder of ligation barcode dictionary
ligation_barcode_file="/barcode_files/ligation_bc_210923.pickle2"

#define the folder of inner i7 barcode dictionary
inner_i7_bc_file=/barcode_files/simp_inner_i7_220517.pickle2"

#define the folder of gRNA barcode dictionary
gRNA_correction_file="/barcode_files/simp_gRNAseq_screen_220824.pickle2"

#define the folder containing the gRNA annotation file
gRNA_annotation_df="/barcode_files/screen_gRNA_info_table.txt"

################################################################################

#####change the environment for sgRNA counting
bashrc_location=/zxu/.bashrc
source $bashrc_location
conda activate singlecell

#####change the file names of raw fastq.gz
echo "Changing the name of the fastq files..."
for sample in $(cat $sample_ID); do echo changing name $sample; mv $fastq_folder/*$sample*R1_001.fastq.gz $fastq_folder/$sample.R1.fastq.gz; mv $fastq_folder/*$sample*R2_001.fastq.gz $fastq_folder/$sample.R2.fastq.gz; mv $fastq_folder/*$sample*R3_001.fastq.gz $fastq_folder/$sample.R3.fastq.gz; done

#####run the main script
echo "Start identifying single cells, gRNA sequence and UMI..."
python3 $main_script $fastq_folder $sample_ID $all_output_folder $RT_barcode_file $inner_i7_bc_file $ligation_barcode_file $gRNA_correction_file $gRNA_annotation_df $cutoff $core 

#####format the output into a sparse matrix ready for R analysis
report_output_folder=$all_output_folder/gRNA_report
echo "Start exporting the final gRNA expression matrix..."
Rscript $reformatting_script $report_output_folder

echo "All done!"
```
