## Supplementary file4 for "*PerturbSci-Kinetics*: Dissecting key regulators of transcriptome kinetics through scalable single-cell RNA profiling of pooled CRISPR screens": PE_bulkRNAseq_rmdup.sh

```
input_folder=$1
sample=$2
output_folder=$3

picard MarkDuplicates INPUT=$input_folder/${sample}.bam OUTPUT=$output_folder/${sample}.bam REMOVE_DUPLICATES=true \
ASSUME_SORTED=True METRICS_FILE=$output_folder/${sample}.picard_metrics_file.txt VALIDATION_STRINGENCY=LENIENT
echo remove duplicates on sample done: $sample
```
