## Supplementary file4 for "*PerturbSci-Kinetics*: Dissecting key regulators of transcriptome kinetics through scalable single-cell RNA profiling of pooled CRISPR screens": PE_extract_new_reads.sh

```
input_folder=$1 # input folder should include both sam_splitted folder and new_synthesised_reads folder
sample=$2
output_folder=$3
nascent_reads_txt_folder=$4

input_sam=$input_folder/$sample.sam
input_read=$nascent_reads_txt_folder/$sample.txt
picard_zx="/zxu/tools/picard/picard.jar"
java_zx="/zxu/tools/openjdk_v11/jdk-11.0.2/bin/java"
$java_zx -jar $picard_zx FilterSamReads I=$input_sam O=$output_folder/$sample.sam READ_LIST_FILE=$input_read FILTER=includeReadList
echo analyis done: $sample
```
