## Supplementary file4 for "*PerturbSci-Kinetics*: Dissecting key regulators of transcriptome kinetics through scalable single-cell RNA profiling of pooled CRISPR screens": PE_nascent_read_sample_level_integration.R

```
require("tidyverse")
require("BiocParallel")

args = commandArgs(trailingOnly=TRUE)

input_folder=args[1]
sample_ID=args[2]
splitted_id_with_new=args[3]
output_folder=args[4]
core = as.numeric(args[5])

sample_names = (read_csv(sample_ID, col_names = F))$X1
splitted_id = (read_csv(splitted_id_with_new, col_names = F))$X1

mut_ratio_cutoff = 0.3

function_sample <- function(target_id, splitted_id, mut_ratio_cutoff, input_folder, output_folder) {
    cat("Process sample: ", target_id)
    cat("\n")
    
    ###get all file prefix
    all_sub_files_prefix <- splitted_id[grep(pattern=target_id, x=splitted_id)]
    
    all_reads <- lapply(all_sub_files_prefix, function(x){
        read_csv(paste0(input_folder, "/", x, ".csv"), col_names = F)
    })
    
    ###get an integrated table
    all_reads_df <- do.call(rbind, all_reads)
    colnames(all_reads_df) <- c("READ_NAME", "mut_num", "target_mut_num")
    
    ###sum up mut number and collapse the same reads from different sub files
    integrated_reads_table <- all_reads_df %>% group_by(READ_NAME) %>% summarise(mut_num=sum(mut_num), target_mut_num=sum(target_mut_num))
    integrated_reads_table$target_mut_ratio <- integrated_reads_table$target_mut_num/integrated_reads_table$mut_num
    
    ###get passed read names
    passed_new_reads <- integrated_reads_table %>% filter(target_mut_ratio > mut_ratio_cutoff)
    
    ###write the read names
    write.table(passed_new_reads$READ_NAME, file = file.path(output_folder, paste0(target_id, ".txt")), sep = "\t",
              col.names = F, quote = F, row.names = F)
}

bplapply(sample_names, function(target_id) {
    function_sample(target_id=target_id, splitted_id=splitted_id, mut_ratio_cutoff = mut_ratio_cutoff, input_folder = input_folder, output_folder = output_folder)
}, BPPARAM = MulticoreParam(workers = core))
```
