## Supplementary file4 for "*PerturbSci-Kinetics*: Dissecting key regulators of transcriptome kinetics through scalable single-cell RNA profiling of pooled CRISPR screens": PE_select_nascent_reads.R

```
require("tidyverse")
require("BiocParallel")
require("Biostrings")

args = commandArgs(trailingOnly=TRUE)

input_folder=args[1]
sample_ID=args[2]
output_folder=args[3]
core = as.numeric(args[4])
SNP_VCF = args[5]

quality_filter = 45
target_mut_filter_rate = 0.3

if(!file.exists(output_folder)) {
    dir.create(output_folder)
}

sample_names = (read_csv(sample_ID, col_names = F))$X1

#sample_names = sample_names[1:5]

SNP = read.table(SNP_VCF, sep =  "\t", header = T)
SNP$chr_pos = str_c(SNP$Chrom, SNP$Position, SNP$Ref, SNP$Var, sep = "-")

function_sample <- function(target_id, input_folder, output_folder, core, SNP, quality_filter = 45, target_mut_filter_rate = 0.3) {
    cat("Process sample: ", target_id)
    cat("\n")
    align_file = file.path(input_folder, paste0(target_id, ".align"))
    
    ###comsidering not all splitted align files have headers, rebuild headers here
    test_input = read_tsv(align_file, col_types = cols(.default = "c"), col_names=F)
    if(test_input[1,1] == "#Read-Name"){
        test_input <- test_input[2:nrow(test_input),]
    }
        
    colnames(test_input)[1] = "READ_NAME"
    colnames(test_input)[2] = "FLAG"
    colnames(test_input)[3] = "MAPQ"
    colnames(test_input)[4] = "CHROM"
    colnames(test_input)[5] = "READ_POS"
    colnames(test_input)[6] = "BASE"
    colnames(test_input)[7] = "QUAL"
    colnames(test_input)[8] = "REF_POS"
    colnames(test_input)[9] = "REF"
    colnames(test_input)[10] = "CIGAR-OP"
    
    ori_test = test_input
    ori_test$READ_POS = as.numeric(as.character(ori_test$READ_POS))
    ori_test$REF_POS = as.numeric(as.character(ori_test$REF_POS))
    ori_test$FLAG = as.numeric(as.character(ori_test$FLAG))

    test_input = ori_test
    test_input = test_input %>% filter(!is.na(CHROM))
    test_input = test_input %>% filter(BASE != ".", REF != ".")
    test_input$REF = str_to_upper(test_input$REF)
    test_input$BASE = str_to_upper(test_input$BASE)
    ####only choose PE reads that are properly aligned and only select bases have CIGAR "M" and have mismatches
    test_input = test_input %>% filter(((BASE) != (REF)) & (FLAG == 83|FLAG == 163|FLAG == 99|FLAG == 147) & (`CIGAR-OP` == "M"))
    
    if(nrow(test_input) == 0) {
        return(-1)
    }
    
    # Since jvarkit use plus strand sequence as concensus readout 
    # Bases on reads mapped to the minus strand will be transformed to the plus strand
    # And Varscan uses plus strand sequences as output also, so directly removing SNP can be done here
    # Remove SNPs
    test_input$chr_pos = str_c(test_input$CHROM, test_input$REF_POS, test_input$REF, test_input$BASE, sep = "-")
    test_input = test_input %>% filter(!(chr_pos %in% SNP$chr_pos))
    
    if(nrow(test_input) == 0) {
        return(-1)
    }
    test_input = test_input[(sapply(test_input$QUAL, utf8ToInt)) > quality_filter, ]
    if(nrow(test_input) == 0) {
        return(-1)
    }

    # Count the number of total mutations in each read
    tmp_mut_num = test_input %>% group_by(READ_NAME) %>% summarise(mut_num = n())
    # Count the T>C number on reads mapped to the plus strand and A>G number on reads mapped to the minus strand
    #99/163 means reads are aligned to the plus strand, 83/147 means reads are aligned to the minus strand
    tmp_target_mut_num = (test_input %>% filter(((FLAG == 99|FLAG == 163) & REF == "T" & BASE == "C")| ((FLAG == 83|FLAG == 147) & REF == "A" & BASE == "G")) %>% group_by(READ_NAME) %>% summarise(target_mut_num = n()))
    tmp_mut_num = left_join(tmp_mut_num, tmp_target_mut_num)
    tmp_mut_num$target_mut_num = ifelse(is.na(tmp_mut_num$target_mut_num), 0, tmp_mut_num$target_mut_num)
    
    ### now the dataframe has 3 columns: READ_NAME, mut_num, target_mut_num
    ### export this table and merge later after mutation counting
    write.table(tmp_mut_num, file = file.path(output_folder, paste0(target_id, ".csv")), sep = ",",
              col.names = F, quote = F, row.names = F)
    
}

bplapply(sample_names, function(target_id) {
    function_sample(target_id, input_folder = input_folder, output_folder = output_folder, core = core, SNP = SNP, quality_filter = quality_filter, target_mut_filter_rate = target_mut_filter_rate)
}, BPPARAM = MulticoreParam(workers = core))
```
