## Supplementary file4 for "*PerturbSci-Kinetics*: Dissecting key regulators of transcriptome kinetics through scalable single-cell RNA profiling of pooled CRISPR screens": SLAM_seq_main_processing.sh

```
#! /bin/bash

################################Change parameters and directories########################################

bashrc_location=/asziraki/original_pipeline/sci-RNA-seq3_pipeline/.bashrc
# 'original_pipeline' conda environment is necessary to run the pipeline
# the scRNA-seq pipeline accept a input folder, and then use the default parameter for sequencing processing and generating the gene count matrix for downstream analysis

# The script is for processing sci-RNA-seq3 sequencing reads in UW genomic science cluster. For running in other environment, some modules, R and python packages are needed to be installed.

# define the fastq folder including all fastq files
fastq_folder="/zxu/230316_SLAMseq/raw_data"

#define the reference genome file used for mutation calling
ref_genome_fa="/zxu/gencode_ref_anno/GRCh38.primary_assembly.genome.fa"

#define the common variants file called from the reference cell line for scifate SNP filtering
ref_SNP_var_file="/zxu/230316_SLAMseq/ref_HEK_vcf/SNP/external_HEK293_ref_SNP.vcf"

# define the PCR group sample id for each fastq file
sample_ID="/zxu/230316_SLAMseq/intermediate_data/sampleID.txt"

# define the output folder
all_output_folder="/zxu/230316_SLAMseq/intermediate_data"

# define the core number for parallele processing
core=16 
samtools_core=4 

# define the location of index files for reads alignment with STAR
index="/zxu/gencode_ref_anno/hg38_STAR_index"

# define the gtf file for gene counting
gtf_file="/zxu/gencode_ref_anno/gencode.v38.primary_assembly.annotation.gtf.gz"


# Define the location of the sub script folder
script_folder="/zxu/scripts/bulk_PERNAseq"

#define the bin of python (python V2.7)
python_path="/asziraki/anaconda3/envs/original_pipeline/bin"

#define paths of specific softwares
java_zx="/zxu/tools/openjdk_v11/jdk-11.0.2/bin/java"
sam2tsv_zx="/zxu/tools/jvarkit/dist/sam2tsv.jar"

# define the location of the R script for multi-core processing
R_script=$script_folder/sci3_bash_input_ID_output_core.R
script_path=$script_folder

#########################################################################################################

now=$(date)
echo "Current time : $now"

############ change the names of the files

###set the env
conda deactivate
source $bashrc_location
conda activate original_pipeline

input_folder=$fastq_folder
echo "Changing the name of the fastq files..."
for sample in $(cat $sample_ID); do echo changing name $sample; mv $input_folder/*$sample*R1_001.fastq.gz $input_folder/$sample.R1.fastq.gz; mv $input_folder/*$sample*R3_001.fastq.gz $input_folder/$sample.R2.fastq.gz; done

################# Trimming read1 and read2
echo
echo "Start trimming reads..."
echo $(date)

raw_fastq=$fastq_folder
trimmed_fastq=$all_output_folder/trimmed_fastq
bash_script=$script_path/PE_bulkRNAseq_trim.sh

mkdir -p $trimmed_fastq

Rscript $R_script $bash_script $raw_fastq $sample_ID $trimmed_fastq $core

############align the reads with STAR, filter the reads based on q > 30, and remove duplicates by picard

#define the output folder for mapping
input_folder=$trimmed_fastq
STAR_output_folder=$all_output_folder/STAR_alignment
filtered_sam_folder=$all_output_folder/filtered_sam
rmdup_sam_folder=$all_output_folder/rmdup_sam

#align read2 to the index file using STAR
echo "Start alignment using STAR..."
echo input folder: $input_folder
echo sample ID file: $sample_ID
echo index file: $index
echo output_folder: $STAR_output_folder

#make the output folder
mkdir -p $STAR_output_folder

###set the env
source /zxu/.bashrc
conda activate singlecell

#remove the index from the memory
STAR --genomeDir $index --genomeLoad Remove
#start the alignment
for sample in $(cat $sample_ID); do echo Aligning $sample;STAR --runThreadN $core --outSAMstrandField intronMotif --genomeDir $index --readFilesCommand zcat --readFilesIn $input_folder/$sample*R1.fastq.gz $input_folder/$sample*R2.fastq.gz --outFileNamePrefix $STAR_output_folder/$sample --genomeLoad LoadAndKeep --outFilterMismatchNoverLmax 0.3 --outFilterMatchNminOverLread 0.2 --outFilterScoreMinOverLread 0.2 --outFilterMultimapNmax 1; done
#remove the index from the memory
STAR --genomeDir $index --genomeLoad Remove
echo "All alignment done."

#Filter and sort the sam file
echo
echo "Start filter and sort the sam files..."
echo input folder: $STAR_output_folder
echo output folder: $filtered_sam_folder
mkdir -p $filtered_sam_folder
    
for sample in $(cat $sample_ID); do
samtools view -bh -q 30 -@ $core $STAR_output_folder/$sample*.sam|samtools sort -@ $core -|samtools view -bh -@ $core ->$filtered_sam_folder/$sample.bam
samtools index -@ $core $filtered_sam_folder/$sample.bam
done

echo
echo "Start removing duplicates..."
echo input folder: $filtered_sam_folder
echo output folder: $rmdup_sam_folder
mkdir -p $rmdup_sam_folder

bash_script=$script_folder/PE_bulkRNAseq_rmdup.sh
Rscript $R_script $bash_script $filtered_sam_folder $sample_ID $rmdup_sam_folder $core

###convert to sam for alignment file generation
for sample in $(cat $sample_ID)
do samtools view -h -@ $core $rmdup_sam_folder/${sample}.bam > $rmdup_sam_folder/${sample}.sam
done

echo "Deduplication is done."

################### generate base-resolution alignment files 
echo "Start generating reads alignment files."

input_folder=$rmdup_sam_folder
alignment_folder=$all_output_folder/alignment_for_mut_call
mkdir -p $alignment_folder

bash_script=$script_folder/PE_seq_align.sh

Rscript $R_script $bash_script $input_folder $sample_ID $alignment_folder $core $ref_genome_fa

echo "Alignment files have been generated."

################### split the big alignment file for more efficient processing
input_folder=$all_output_folder/alignment_for_mut_call
output_folder=$all_output_folder/splitted_alignment_for_mut_call
mkdir -p $output_folder

bash_script=$script_folder/PE_bulkRNAseq_split_alignment_file.sh

Rscript $R_script $bash_script $input_folder $sample_ID $output_folder $core

echo "Alignment splitting is done."

##generate a sample id list for input
ls $output_folder > $all_output_folder/splitted_alignment_id.txt
sed -i "s/.align//g" $all_output_folder/splitted_alignment_id.txt

################### filter mutations on reads and get nascent reads names
echo "Start searching for nascent reads from single cell sams."

input_folder=$all_output_folder/splitted_alignment_for_mut_call
output_folder=$all_output_folder/new_reads_txt_Monly
splitted_id=$all_output_folder/splitted_alignment_id.txt
mkdir -p $output_folder

process_R_script=$script_folder/PE_select_nascent_reads.R

Rscript $process_R_script $input_folder $splitted_id $output_folder $core $ref_SNP_var_file

ls $output_folder > $all_output_folder/splitted_id_with_new.txt
sed -i "s/.csv//g" $all_output_folder/splitted_id_with_new.txt

echo "Nascent reads have been retrieved."

#################### Integrate the mutations identified at the sample level
input_folder=$all_output_folder/new_reads_txt_Monly
output_folder=$all_output_folder/sampled_level_new_reads_txt_Monly
splitted_id_with_new=$all_output_folder/splitted_id_with_new.txt
mkdir -p $output_folder

integration_R_script=$script_folder/PE_nascent_read_sample_level_integration.R

Rscript $integration_R_script $input_folder $sample_ID $splitted_id_with_new $output_folder $core

echo "new reads sams have been integrated."

###considering the storage issue, delete the alignment file after this step
rm -rf $all_output_folder/alignment_for_mut_call
rm -rf $all_output_folder/splitted_alignment_for_mut_call

#################### extract these reads from rmdup sam files
echo "Start extracting nascent reads from each file."

input_folder=$rmdup_sam_folder
new_reads_txt_folder=$all_output_folder/sampled_level_new_reads_txt_Monly
output_folder=$all_output_folder/new_reads_sam_Monly
mkdir -p $output_folder

bash_script=$script_folder/PE_extract_new_reads.sh

Rscript $R_script $bash_script $input_folder $sample_ID $output_folder $core $new_reads_txt_folder

echo "new reads sams have been generated."

####also transform to bam, sort and index
for sample in $(cat $sample_ID)
do samtools view -bh -@ $core $output_folder/${sample}.sam | samtools sort -@ $core - > $output_folder/${sample}.bam
samtools index -@ $core $output_folder/${sample}.bam
done

################### calculate the reads number
fastq_folder=$fastq_folder
trimmed_folder=$trimmed_fastq
alignment=$STAR_output_folder
filtered_sam=$filtered_sam_folder
rm_dup_sam=$rmdup_sam_folder
new_reads_sam=$all_output_folder/new_reads_sam_Monly
report_folder=$all_output_folder/report/read_num

echo
echo "Start calculating the reads number..."

#make the report folder
mkdir -p $report_folder

#calculate the read number and output the read number into the report folder
echo sample,total reads,after after trimming,uniquely aligned reads,After remove duplicates,Nascent reads>$report_folder/read_number.csv
for sample in $(cat $sample_ID); do echo calculating $sample; echo $sample,$(expr $(zcat $fastq_folder/$sample*R1*.gz|wc -l) / 2),$(expr $(zcat $trimmed_folder/$sample*R1*.gz|wc -l) / 2),$(samtools view $filtered_sam/$sample.bam|wc -l),$(samtools view $rm_dup_sam/$sample.bam|wc -l),$(samtools view $new_reads_sam/$sample.bam|wc -l)>>$report_folder/read_number.csv; done
echo "Read number calculation is done."

#################### gene level feature counts
total_bam_folder=$all_output_folder/rmdup_sam
new_bam_folder=$all_output_folder/new_reads_sam_Monly
report_folder=$all_output_folder/report/read_counts
mkdir -p $report_folder

for sample in $(cat $sample_ID)
do featureCounts -a $gtf_file -o $report_folder/${sample}.total.count -T $core -t gene -g gene_name -p -s 2 $total_bam_folder/${sample}.bam
featureCounts -a $gtf_file -o $report_folder/${sample}.new.count -T $core -t gene -g gene_name -p -s 2 $new_bam_folder/${sample}.bam
done

conda deactivate
```
