## Supplementary file4 for "*PerturbSci-Kinetics*: Dissecting key regulators of transcriptome kinetics through scalable single-cell RNA profiling of pooled CRISPR screens": Main_processing.sh

```
#! /bin/bash

###################define the directories########################

## define the directory of raw fastq files
fastq_folder=""

## define the identifier for sample identification
sample_ID="/example_files/sampleID.txt"

## define the output folder
all_output_folder=""

## define the core number for parallel processing
core=16
samtools_core=4

## define the number of UMI cutoff for splitting single cell; cells with UMIs less than this number will be discarded
cutoff=200

## define the location of index files for reads alignment with STAR and the annotation
index=""
gtf_file=""

## define the reference genome used for mutation calling
ref_genome_fa=""
ref_SNP_var_file="/example_files/mutation.vcf"

## define the script folder
script_folder="/main_scripts"

## define the bin of python
python_path="/asziraki/anaconda3/envs/original_pipeline/bin"

# define the location of the ligation barcodes (they are in the script folder)
custom_barcode_folder="/barcode_files"
ligation_barcode=$custom_barcode_folder/ligation_bc_210923.pickle2

# define the location of the RT barcodes
RT_barcode=$custom_barcode_folder/RT_bc_210923.pickle2

# define the location of the combined RT and ligation barcodes
barcodes=$custom_barcode_folder/combined_ligation_RT_210923.txt

# define the location of the R script for multi-core processing
R_script=$script_folder/sci3_bash_input_ID_output_core.R
script_path=$script_folder

#################################################################

# Activate the main env for easysci processing
bashrc_location=/asziraki/projects/AS_20200820_single_cell_pipeline/Scripts/original_pipeline/sci-RNA-seq3_pipeline/.bashrc
source $bashrc_location
conda activate original_pipeline

now=$(date)
echo "Current time : $now"

############ UMI attach
# the script take an input folder, a sample ID list, an output folder, the RT barcode list, the ligation barcode list and core number. Then it extract the RT barcode from read1, the ligation barocde from read2, correct them to the nearest RT and ligation barcode (with edit distance <= 1), and attach the RT and ligation barcode and UMI sequence to the read name of read3. Reads with unmatched RT or ligation barcodes are discarded. Reads with matched sgRNA capture primer sequence are also discarded.

input_folder=$fastq_folder
output_folder=$all_output_folder/UMI_attach
script=$script_folder/UMI_barcode_attach.py
echo "Changing the name of the fastq files..."
for sample in $(cat $sample_ID); do echo changing name $sample; mv $input_folder/*$sample*R1_001.fastq.gz $input_folder/$sample.R1.fastq.gz; mv $input_folder/*$sample*R2_001.fastq.gz $input_folder/$sample.R2.fastq.gz; mv $input_folder/*$sample*R3_001.fastq.gz $input_folder/$sample.R3.fastq.gz; done

echo "Attaching barcode and UMI...."
mkdir -p $output_folder
$python_path/python $script $input_folder $sample_ID $output_folder $ligation_barcode $RT_barcode $core
echo "Barcode transformed and UMI attached."

############# Trimming the read2
echo
echo "Start trimming the read2 file..."
echo $(date)

trimmed_fastq=$all_output_folder/trimmed_fastq
UMI_attached_R2=$all_output_folder/UMI_attach
bash_script=$script_path/sci3_trim.sh

Rscript $R_script $bash_script $UMI_attached_R2 $sample_ID $trimmed_fastq $core

############# align the reads with STAR, filter the reads based on q > 30, and remove duplicates based on UMI sequence and tagmentation site

# define the output folder for mapping
input_folder=$trimmed_fastq
STAR_output_folder=$all_output_folder/STAR_alignment
filtered_sam_folder=$all_output_folder/filtered_sam
rmdup_sam_folder=$all_output_folder/rmdup_sam

# align read2 to the index file using STAR

echo "Start alignment using STAR..."
echo input folder: $input_folder
echo sample ID file: $sample_ID
echo index file: $index
echo output_folder: $STAR_output_folder

# make the output folder
mkdir -p $STAR_output_folder

# change to the environment for alignment

conda deactivate
source /home/zxu/.bashrc
conda activate singlecell

# remove the index from the memory

STAR --genomeDir $index --genomeLoad Remove

# start the alignment
for sample in $(cat $sample_ID); do echo Aligning $sample;STAR --runThreadN $core --outSAMstrandField intronMotif --genomeDir $index --readFilesCommand zcat --readFilesIn $input_folder/$sample*gz --outFileNamePrefix $STAR_output_folder/$sample --genomeLoad LoadAndKeep; done

#remove the index from the memory
STAR --genomeDir $index --genomeLoad Remove
echo "All alignment done."

# change to the main pipeline environment

conda deactivate
source $bashrc_location
conda activate original_pipeline

############# Filter and sort the sam file

echo
echo "Start filter and sort the sam files..."
echo input folder: $STAR_output_folder
echo output folder: $filtered_sam_folder
bash_script=$script_path/sci3_filter.sh
Rscript $R_script $bash_script $STAR_output_folder $sample_ID $filtered_sam_folder $samtools_core

# make a folder for rmdup_sam_folder, 
# Then for each filtered sam file, remove the duplicates based on UMI and barcode, chromatin number and position
# Remove duplicates based on UMI sequence (exact match) and tagmentation site

echo
echo "Start removing duplicates..."
echo input folder: $filtered_sam_folder
echo output folder: $rmdup_sam_folder
mkdir -p $rmdup_sam_folder
bash_script=$script_path/sci3_rmdup_nomismatch.sh
Rscript $R_script $bash_script $filtered_sam_folder $sample_ID $rmdup_sam_folder $core

################# split the sam file based on the barcode, and mv the result to the report folder

sam_folder=$all_output_folder/rmdup_sam
output_folder=$all_output_folder/sam_splitted

echo
echo "Start splitting the sam file..."
echo samfile folder: $sam_folder
echo sample list: $sample_ID
echo ouput folder: $output_folder
echo barcode file: $barcodes
echo cutoff value: $cutoff

bash_script=$script_path/sci3_split.sh
Rscript $R_script $bash_script $sam_folder $sample_ID $output_folder $core $barcodes $cutoff

cat $output_folder/*sample_list.txt > $output_folder/All_samples.txt
cp $output_folder/All_samples.txt $output_folder/../barcode_samples.txt

# output the report the report/barcode_read_distribution folder
mkdir -p $output_folder/../report/barcode_read_distribution
mv $output_folder/*.txt $output_folder/../report/barcode_read_distribution/
mv $output_folder/*.png $output_folder/../report/barcode_read_distribution/
echo
echo "All sam file splitted."

################### calculate the reads number

fastq_folder=$fastq_folder
trimmed_folder=$trimmed_fastq
UMI_attach=$UMI_attached_R2
alignment=$STAR_output_folder
filtered_sam=$filtered_sam_folder
rm_dup_sam=$rmdup_sam_folder
report_folder=$all_output_folder/report/read_num

echo
echo "Start calculating the reads number..."

# make the report folder
mkdir -p $report_folder

# calculate the read number and output the read number into the report folder
echo sample,total reads,after filtering barcode,after trimming,uniquely aligned reads,After remove duplicates>$report_folder/read_number.csv
for sample in $(cat $sample_ID); do echo calculating $sample; echo $sample,$(expr $(zcat $fastq_folder/$sample*R2*.gz|wc -l) / 4),$(expr $(zcat $UMI_attach/$sample*R2*.gz|wc -l) / 4),$(expr $(zcat $trimmed_folder/$sample*R2*.gz|wc -l) / 4),$(samtools view $filtered_sam/$sample.sam|wc -l),$(samtools view $rm_dup_sam/$sample.sam|wc -l)>>$report_folder/read_number.csv; done
echo "Read number calculation is done."


################### generate single cell alignment files 

echo "Start generating single cell reads alignment files."

input_folder=$all_output_folder/sam_splitted
sc_alignment_folder=$all_output_folder/sc_alignment
mkdir -p $sc_alignment_folder
sc_barcode_list=$all_output_folder/barcode_samples.txt
bash_script=$script_path/seq_align.sh

#change to the env for nascent reads processing
conda deactivate
source /home/zxu/.bashrc
conda activate singlecell

Rscript $R_script $bash_script $input_folder $sc_barcode_list $sc_alignment_folder $core $ref_genome_fa

echo "Single cell alignment files have been generated."


################### filter mutations on reads and get nascent reads names

echo "Start searching for nascent reads from single cell sams."

input_folder=$all_output_folder/sc_alignment
output_folder=$all_output_folder/new_reads_txt_Monly
mkdir -p $output_folder

process_R_script=$script_path/select_nascent_reads.R

# filter the newly synthesised reads for each single cell
Rscript $process_R_script $input_folder $sc_barcode_list $output_folder $core $ref_SNP_var_file

echo "Single cell nascent reads have been retrieved."


################### extract these reads from splitted sam files
echo "Start extracting nascent reads from single cell sams."

input_folder=$all_output_folder/sam_splitted
new_reads_txt_folder=$all_output_folder/new_reads_txt_Monly
output_folder=$all_output_folder/new_reads_sam_splitted_Monly
mkdir -p $output_folder

bash_script=$scifate_script_folder/extract_new_reads.sh

Rscript $R_script $bash_script $input_folder $sc_barcode_list $output_folder $core $new_reads_txt_folder

echo "new reads single cell sams have been generated."

# generate a nascent only cell name file
ls $output_folder > $all_output_folder/nascent_barcode_samples_Monly.txt
sed -i "s/.sam//g" $all_output_folder/nascent_barcode_samples_Monly.txt

################### gene count on nascent txome
echo "Start gene counting on nascent txome."

# change to the main env for easysci processing
conda deactivate
source $bashrc_location
conda activate original_pipeline

output_folder=$all_output_folder/nascent_txme_report_Monly/human_mouse_gene_count
input_folder=$all_output_folder/new_reads_sam_splitted_Monly
script=$script_path/sciRNAseq_count.py
sample_ID=$all_output_folder/nascent_barcode_samples_Monly.txt

$python_path/python $script $gtf_file $input_folder $sample_ID $core

echo "Make the output folder and transfer the nascent reads counting files..."
mkdir -p $output_folder

find $input_folder/ -name "*.count" -print0 | sort -z | xargs -r0 cat > $output_folder/count.MM
find $input_folder/ -name "*.count" -delete
find $input_folder/ -name "*.report" -print0 | sort -z | xargs -r0 cat > $output_folder/report.MM
find $input_folder/ -name "*.report" -delete
mv $input_folder/*_annotate.txt $output_folder/

echo "All output files are transferred."

################# gene count on whole txome
echo "Start gene counting on whole txome."

# count reads mapping to genes
output_folder=$all_output_folder/whole_txme_report/human_mouse_gene_count
input_folder=$all_output_folder/sam_splitted
script=$script_path/sciRNAseq_count.py
sample_ID=$all_output_folder/barcode_samples.txt

$python_path/python $script $gtf_file $input_folder $sample_ID $core

echo "Make the output folder and transfer the files..."
mkdir -p $output_folder

find $input_folder/ -name "*.count" -print0 | sort -z | xargs -r0 cat > $output_folder/count.MM
find $input_folder/ -name "*.count" -delete
find $input_folder/ -name "*.report" -print0 | sort -z | xargs -r0 cat > $output_folder/report.MM
find $input_folder/ -name "*.report" -delete
mv $input_folder/*_annotate.txt $output_folder/

echo "All output files are transferred."

################# transform output files to an easy to read in R file

# change to the env for count matrix reformatting
conda deactivate
conda activate original_pipeline_final_step

R_script=$script_path/gene_count_processing_sciRNAseq_exon_intron.R
Rscript $R_script $all_output_folder/whole_txme_report

R_script=$script_path/gene_count_processing_sciRNAseq_exon_intron.R
Rscript $R_script $all_output_folder/nascent_txme_report_Monly
conda deactivate

now=$(date)
echo "Current time : $now"
```
