## Supplementary file4 for "*PerturbSci-Kinetics*: Dissecting key regulators of transcriptome kinetics through scalable single-cell RNA profiling of pooled CRISPR screens": rm_dup_barcode_UMI_no_mismatch.py

```
'''
This script accept a input sorted sam file, a output sam file, then it will remove
duplicates based on the barcode + UMI (edit distance <= 1), and chromatin and start site, at the same
time, it will output the duplication number for each read, and generate the histogram plot for the read
per duplication number
'''
from Levenshtein import distance
import sys
import matplotlib as mpl
import numpy as np
mpl.use('Agg')
import matplotlib.pyplot as plt
import pandas as pd
sys.setrecursionlimit(1000000)
def rm_dup_samfile(samfile, output_file):
    f1 = open(samfile)
    f2 = open(output_file, 'w')
    f3 = open(output_file+'.csv', 'w')
    pre_barcode = []
    pre_line = []
    unique_id = []
    pre_chrom = 0
    pre_site = 0
    dup = False
    
    pre_dup_num = 0
    cur_dup_num = 0
    
    for line in f1:
        
        if (line[0] == '@'):
            f2.write(line)
        else:
            name = (((line.split('\t'))[0]).split(','))
            barcode_UMI = name[0] + name[1]
            chrom_num = (line.split('\t'))[2]
            start_site = (line.split('\t'))[3]
            
            if ((start_site == pre_site) and (chrom_num == pre_chrom)):    
                dup = False
                for each_barcode in pre_barcode:
                    if each_barcode == barcode_UMI:
                        dup = True
                        break
                if dup == False:
                    pre_dup_num = cur_dup_num
                    cur_dup_num = 1
                    f2.write(line)
                    pre_barcode.add(barcode_UMI)
                    f3.write('%d' %(pre_dup_num))
                    f3.write('\n')
                else:
                    cur_dup_num += 1               
            else:
                pre_dup_num = cur_dup_num
                cur_dup_num = 1
                f2.write(line)
                pre_chrom = chrom_num
                pre_site = start_site
                pre_barcode = set()
                pre_barcode.add(barcode_UMI)
                if (pre_dup_num != 0):
                    f3.write("%d" % (pre_dup_num))
                    f3.write('\n')
    
    f1.close()
    f2.close()
    f3.close()
    
    '''
    #plot the histogram for the read duplication number
    dups = (pd.read_csv(output_file+'.csv', header=None))[0]
    fig = plt.figure()
    plt.hist(dups, bins=100)
    plt.xlabel("Duplication number")
    plt.ylabel("Read number")
    fig.savefig(output_file + '.png')
    '''

if __name__ == "__main__":
    samfile = sys.argv[1]
    output_file = sys.argv[2]
    rm_dup_samfile(samfile, output_file)
```
