## Supplementary file4 for "*PerturbSci-Kinetics*: Dissecting key regulators of transcriptome kinetics through scalable single-cell RNA profiling of pooled CRISPR screens": sci3_rmdup_nomismatch.sh

```
input_folder=$1
sample=$2
output_folder=$3
mismatch=$4

python="/rugpfs/fs0/cao_lab/scratch/asziraki/anaconda3/envs/original_pipeline/bin/python2.7"
python_script="/main_scripts/rm_dup_barcode_UMI_no_mismatch.py"

echo Filtering sample: $sample

$python $python_script $input_folder/$sample.sam $output_folder/$sample.sam $mismatch

echo Filtering $sample done.
```
