## Supplementary file4 for "*PerturbSci-Kinetics*: Dissecting key regulators of transcriptome kinetics through scalable single-cell RNA profiling of pooled CRISPR screens": sci3_split.sh

```
input_folder=$1
sample=$2
output_folder=$3
barcode_file=$4
cutoff=$5

python="/rugpfs/fs0/cao_lab/scratch/asziraki/anaconda3/envs/original_pipeline/bin/python2.7"
python_script="/main_scripts/sam_split.py"

$python $python_script $input_folder/$sample.sam $barcode_file $output_folder $cutoff
echo splitting sample done: $sample
```
