## Supplementary file4 for "*PerturbSci-Kinetics*: Dissecting key regulators of transcriptome kinetics through scalable single-cell RNA profiling of pooled CRISPR screens": sciRNAseq_count.py

```
# This script take as input a gtf file, a sam file input folder, a sample ID file, and the number of cores, then run
# the count exons and genes in parallel, and then output the report file and output file into the output file

import itertools
import collections
import numpy as np
import pandas as pd
from multiprocessing import Pool
from multiprocessing import *
import HTSeq
import sys
from functools import partial
import logging

# read in the sam file and then count genes
# First, for each reads, for each gapped reads, find the contained exons, and select in the intersected exons of all unions;
# If not find in intersected exons, for the contained exons, select in the combined union of all exons,
# If not find in any exons, then intersect in genes, and find the contained exons, and select in the intersected exons of all unions;
# If not find in intersected genes, for the contained exons, select in the combined union of all genes,

# From the above, for all selected genes, if the length of selected gene is only 1, then the gene is the target gene. 
# If the gene list has more than 1 gene, then  substract the end point of the genes and compare the distance to the end point of 
# reads, and compare the end point, and then filter the genes with the shortest length of gene (If the there are more than
# one gene to the 100 + end_distance, then report "_ambiguous")

def sciRNAseq_count(sample, input_folder, exons, genes, gene_end, gene_annotat, sample_ID):
    input_sam = input_folder + "/" + sample + ".sam"
    report = input_folder + "/" + sample + ".report"
    count_output = input_folder + "/" + sample + ".count"
    
    counts = collections.Counter()
    sam_file = input_sam
    almnt_file = HTSeq.SAM_Reader(sam_file)
    sam_name = sample
    cell_ID = sample_ID.index(sample) + 1

    perfect_inter_exon = 0
    nearest_inter_exon = 0
    perfect_combine_exon = 0
    nearest_combine_exon = 0
    perfect_inter_gene = 0
    nearest_inter_gene = 0
    perfect_combine_gene = 0
    nearest_combine_gene = 0
    
    print("Start read the input file: " + sam_file + "....")

    for alnmt in almnt_file:
        #print alnmt
        if not alnmt.aligned:
            counts["_unmapped"] += 1
            continue
        
        if alnmt.iv.chrom not in genes.chrom_vectors:
            counts["_unmapped"] += 1
            continue

        # First check the intersectin with exons
        gene_id_intersect = set()
        gene_id_combine = set()
        inter_count = 0
        for cigop in alnmt.cigar:
            if cigop.type != "M":
                continue

            for iv,val in exons[cigop.ref_iv].steps():
                #print iv, val
                gene_id_combine |= val
                if inter_count == 0:
                    gene_id_intersect |= val
                    inter_count += 1
                else:
                    gene_id_intersect &= val
                #print "intersect set:", gene_id_intersect
                #print "combine set:", gene_id_combine
        # first check the intersection set
        if len(gene_id_intersect) == 1:
            gene_id = list(gene_id_intersect)[0]
            counts[gene_id] += 1
            perfect_inter_exon += 1
        elif len(gene_id_intersect) > 1:
            gene_id = find_nearest_gene(alnmt.iv.end_d, gene_id_intersect, gene_end)
            counts[gene_id] += 1
            nearest_inter_exon += 1
        else:
            # if there no intersection match, then find the union sets
            if len(gene_id_combine) == 1:
                gene_id = list(gene_id_combine)[0]
                counts[gene_id] += 1
                perfect_combine_exon += 1
            elif len(gene_id_combine) > 1:
                gene_id = find_nearest_gene(alnmt.iv.end_d, gene_id_combine, gene_end)
                counts[gene_id] += 1
                nearest_combine_exon += 1
            else:
                # if there is no intersection match or union match, then search for genes to find the intronic match
                gene_id_intersect = set()
                gene_id_combine = set()
                inter_count = 0
                for cigop in alnmt.cigar:
                    if cigop.type != "M":
                        continue
                    for iv,val in genes[cigop.ref_iv].steps():
                        gene_id_combine |= val
                        if inter_count == 0:
                            gene_id_intersect |= val
                            inter_count += 1
                        else:
                            gene_id_intersect &= val

                if len(gene_id_intersect) == 1:
                    gene_id = list(gene_id_intersect)[0] + "_intron"
                    counts[gene_id] += 1
                    perfect_inter_gene += 1

                elif len(gene_id_intersect) > 1:
                    gene_id = find_nearest_gene(alnmt.iv.end_d, gene_id_intersect, gene_end) + "_intron"
                    counts[gene_id] += 1
                    nearest_inter_gene += 1

                else:
                    # if there no intersection match, then find the union sets
                    if len(gene_id_combine) == 1:
                        gene_id = list(gene_id_combine)[0] + "_intron"
                        counts[gene_id] += 1
                        perfect_combine_gene += 1

                    elif len(gene_id_combine) > 1:
                        gene_id = find_nearest_gene(alnmt.iv.end_d, gene_id_combine, gene_end) + "_intron"
                        counts[gene_id] += 1
                        nearest_combine_gene += 1

                    else:
                        counts["_no_feature"] += 1
                        
    print("File name: ", sam_file)
    print("1: Perfect intersect exon match: ", perfect_inter_exon)
    print("2: Nearest intersect exon match: ", nearest_inter_exon)
    print("3: Perfect combine exon match: ", perfect_combine_exon)
    print("4: Nearest combine exon match: ", nearest_combine_exon)
    print("5: Perfect intersect gene match: ", perfect_inter_gene)
    print("6: Nearest intersect gene match: ", nearest_inter_gene)
    print("7: Perfect combine gene match: ", perfect_combine_gene)
    print("8: Nearest combine gene match: ", nearest_combine_gene)
    print("9: ambiguous match for exons: ", counts["_ambiguous"])
    print("10: ambiguous match for genes: ", counts["_ambiguous_intron"])
    print("11: No match: ",  counts["_no_feature"])
    print("Sam file analysis finished~")
    
    with open(report, 'w') as report:
        report.write("1" + "," + str(cell_ID) + "," + str(perfect_inter_exon) + "\n")
        report.write("2" + "," + str(cell_ID) + "," + str(nearest_inter_exon) + "\n")
        report.write("3" + "," + str(cell_ID) + "," + str(perfect_combine_exon) + "\n")
        report.write("4" + "," + str(cell_ID) + "," + str(nearest_combine_exon) + "\n")
        report.write("5" + "," + str(cell_ID) + "," + str(perfect_inter_gene) + "\n")
        report.write("6" + "," + str(cell_ID) + "," + str(nearest_inter_gene) + "\n")
        report.write("7" + "," + str(cell_ID) + "," + str(perfect_combine_gene) + "\n")
        report.write("8" + "," + str(cell_ID) + "," + str(nearest_combine_gene) + "\n")
        report.write("9" + "," + str(cell_ID) + "," + str(counts["_ambiguous"]) + "\n")
        report.write("10" + "," + str(cell_ID) + "," + str(counts["_ambiguous_intron"]) + "\n")
        report.write("11" + "," + str(cell_ID) + "," + str(counts["_no_feature"]) + "\n")
        
    with open(count_output, 'w') as count_output:
        for gene in counts:
            if (gene in ["_unmapped", "_ambiguous", "_ambiguous_intron", "_no_feature"]):
                continue
            else:
                line = str(gene_annotat.loc[gene,4]) + "," + str(cell_ID) + "," + str(counts[gene]) + "\n"
                count_output.write(line)
    return 0

# Define a function that accept the end position of the alignments, and a set of genes, and the dictionary of the
# gene end, and then return the gene with the nearest end to the alignment

# for each gene element in the gene_id_intersect, I will find the absolute distance between the alignment end and
# the transcript end, and select the minimum distance for each gene in the list, and then compare the genes, and use
# the nearest gene as the gene id for output

def find_nearest_gene(al_end, gene_id_intersect, gene_end):
    gene_id_end = {}
    for gene in gene_id_intersect:
        if gene in gene_end:
            gene_id_end[gene] = (abs(np.array(list(gene_end[gene])) - al_end)).min()
        else:
            print "****************Found one gene without transcript annotation*****************", "Gene name: ", gene
    # filter the gene with the least distance. If there are two genes with the least distance, then  "_ambiguous" 
    # would be returned
    gene_end_min = np.min(gene_id_end.values())
    count = 0
    #print ("gene end distance: ", gene_id_end.values())
    for gene in gene_id_end:
        if (gene_id_end[gene] < gene_end_min + 100):
            count += 1
            gene_id = gene
    if count > 1:
        gene_id = "_ambiguous"
    
    return(gene_id)
'''
def sciRNAseq_count_mf(sample, input_folder, exons, genes, gene_end):
    try:
        return sciRNAseq_count(sample, input_folder, exons, genes, gene_end)
    except Exception:
        #logging.exception("f(%r) failed" % (sample, input_folder, exons, genes, gene_end,))
        print Exception.args()
'''

def sciRNA_count_parallel(gtf_file, input_folder, sample_ID, core_number):
    # read in the gtf file, and then construct the genome interval for exons, genes, and gene end dictionary
    gtf_file = HTSeq.GFF_Reader(gtf_file, end_included=True)
    gene_annotat_file = input_folder + "/gene_name_annotate.txt"
    cell_annotat_file = input_folder + "/cell_annotate.txt"
    report_annotate_file = input_folder + "/report_annotate.txt"
    
    gene_annotat = open(gene_annotat_file, "w")
    cell_annotat = open(cell_annotat_file, "w")
    report_annotate = open(report_annotate_file, "w")
    
    exons = HTSeq.GenomicArrayOfSets( "auto", stranded=True )
    genes = HTSeq.GenomicArrayOfSets( "auto", stranded=True )
    gene_end = {}
    exon_n = 0
    gene_n = 0
    transcript_n = 0
    gene_count = 0
    print("Start generating exon genomic arrays....")
    print("Start generating gene genomic arrays....")
    print("Start calculating transcript end of genes....")

    for feature in gtf_file:
        if feature.type == "exon":
            exon_n += 1
            exons[ feature.iv ] += feature.attr["gene_id"]
        elif feature.type == "gene":
            gene_n +=1
            genes[ feature.iv ] += feature.attr["gene_id"]
            gene_count += 1
            
            # for human and mouse gtf file
            message = (feature.attr["gene_id"] + "," + feature.attr["gene_type"] + "," 
                       + "exon" + "," + feature.attr["gene_name"] + "," + str(gene_count) + "\n")
            
            gene_annotat.write(message)
            
            gene_count += 1
            
            # for human and mouse gtf file
            message = (feature.attr["gene_id"] + "_intron" + "," + feature.attr["gene_type"] + "," 
                       + "intron" + "," + feature.attr["gene_name"] + "_intron" + "," + str(gene_count) + "\n")
            gene_annotat.write(message)
            
        elif feature.type == "transcript":
            transcript_n += 1
            #print "feature gene name: ", feature.attr["gene_id"]
            if feature.attr["gene_id"] in gene_end.keys():
                gene_end[ feature.attr["gene_id"] ].add(feature.iv.end_d)
            else:
                gene_end[ feature.attr["gene_id"] ] = set()
                gene_end[ feature.attr["gene_id"] ].add(feature.iv.end_d)

    print("Detected gene number: ", gene_n)
    print("Detected transcript number: ", transcript_n)
    print("Detected exon number: ", exon_n)
    
    gene_annotat.close()
    
    gene_annotat = pd.read_csv(gene_annotat_file, header=None)
    gene_annotat.index =  gene_annotat[0]
    
    sample_ID = list(pd.read_csv(sample_ID, header=None)[0])
    
    # generate the cell ID annotate file
    cell_count = 0
    for i in sample_ID:
        cell_count += 1
        message = i + "," + str(cell_count) + "\n"
        cell_annotat.write(message)
    cell_annotat.close()
    
    # Generate the report annotate file
    report_annotate.write("1, Perfect intersect exon match\n")
    report_annotate.write("2, Nearest intersect exon match\n")
    report_annotate.write("3, Perfect combine exon match\n")
    report_annotate.write("4, Nearest combine exon match\n")
    report_annotate.write("5, Perfect intersect gene match\n")
    report_annotate.write("6, Nearest intersect gene match\n")
    report_annotate.write("7, Perfect combine gene match\n")
    report_annotate.write("8, Nearest combine gene match\n")
    report_annotate.write("9, ambiguous match for exons\n")
    report_annotate.write("10, ambiguous match for genes\n")
    report_annotate.write("11, No match\n")
    report_annotate.close()


    # parallele for the functions
    p = Pool(processes = int(core_number))
    #print("Processing core number: ", core_number)
    func = partial(sciRNAseq_count, input_folder = input_folder, exons=exons, genes=genes, gene_end = gene_end, gene_annotat = gene_annotat, sample_ID = sample_ID)
    #sciRNAseq_count(sample, input_folder, exons, genes, gene_end)
    result = p.map(func, sample_ID)
    p.close()
    p.join()

    print("All analysis done~")
'''
def p_sciRNAseq_count(sample):
    print("get sample: ", sample)
    input_sam = input_folder + "/" + sample + ".sam"
    report = input_folder + "/" + sample + ".report"
    count_output = input_folder + "/" + sample + ".count"
    sciRNAseq_count(sample, input_sam, report, count_output, exons, genes, gene_end)

'''
    
if __name__ == "__main__":
    gtf_file = sys.argv[1]
    input_folder = sys.argv[2]
    sample_ID = sys.argv[3]
    core_number = sys.argv[4]
    sciRNA_count_parallel(gtf_file, input_folder, sample_ID, core_number)
```
