## Supplementary file4 for "*PerturbSci-Kinetics*: Dissecting key regulators of transcriptome kinetics through scalable single-cell RNA profiling of pooled CRISPR screens": select_nascent_reads.R

```
require("tidyverse")
require("BiocParallel")
require("Biostrings")

args = commandArgs(trailingOnly=TRUE)

input_folder=args[1]
sample_ID=args[2]
output_folder=args[3]
core = as.numeric(args[4])
SNP_VCF = args[5]

quality_filter = 45
target_mut_filter_rate = 0.3

if(!file.exists(output_folder)) {
    dir.create(output_folder)
}

sample_names = (read_csv(sample_ID, col_names = F))$X1

SNP = read.table(SNP_VCF, sep =  "\t", header = T)
SNP$chr_pos = str_c(SNP$Chrom, SNP$Position, SNP$Ref, SNP$Var, sep = "-")

function_sample <- function(target_id, input_folder, output_folder, core, SNP, quality_filter = 45, target_mut_filter_rate = 0.3) {
    cat("Process sample: ", target_id)
    cat("\n")
    align_file = file.path(input_folder, paste0(target_id, ".align"))
    test_input = read_tsv(align_file, col_types = cols(.default = "c"))
    colnames(test_input)[1] = "READ_NAME"
    colnames(test_input)[2] = "FLAG"
    colnames(test_input)[5] = "READ_POS"
    colnames(test_input)[6] = "BASE"
    colnames(test_input)[7] = "QUAL"
    colnames(test_input)[8] = "REF_POS"
    colnames(test_input)[9] = "REF"
    
    ori_test = test_input
    ori_test$READ_POS = as.numeric(as.character(ori_test$READ_POS))
    ori_test$REF_POS = as.numeric(as.character(ori_test$REF_POS))
    ori_test$FLAG = as.numeric(as.character(ori_test$FLAG))

    test_input = ori_test
    test_input = test_input %>% filter(!is.na(CHROM))
    test_input = test_input %>% filter(BASE != ".", REF != ".")
    test_input$REF = str_to_upper(test_input$REF)
    test_input$BASE = str_to_upper(test_input$BASE)
    ####only choose reads that are properly aligned and only select bases have CIGAR "M"
    test_input = test_input %>% filter(((BASE) != (REF)) & (FLAG == 0|FLAG == 16) & (`CIGAR-OP` == "M"))
    
    if(nrow(test_input) == 0) {
        return(-1)
    }
    
    # Since jvarkit use plus strand sequence as concensus readout 
    # Bases on reads mapped to the minus strand will be transformed to the plus strand
    # And Varscan uses plus strand sequences as output also, so directly removing SNP can be done here
    # Remove SNPs
    test_input$chr_pos = str_c(test_input$CHROM, test_input$REF_POS, test_input$REF, test_input$BASE, sep = "-")
    test_input = test_input %>% filter(!(chr_pos %in% SNP$chr_pos))
    
    if(nrow(test_input) == 0) {
        return(-1)
    }
    test_input = test_input[(sapply(test_input$QUAL, utf8ToInt)) > quality_filter, ]
    if(nrow(test_input) == 0) {
        return(-1)
    }

    # Count the number of total mutations in each read
    tmp_mut_num = test_input %>% group_by(READ_NAME) %>% summarise(mut_num = n())
    # Count the T>C number on reads mapped to the plus strand and A>G number on reads mapped to the minus strand
    tmp_target_mut_num = (test_input %>% filter((FLAG == 0 & REF == "T" & BASE == "C") | (FLAG == 16 & REF == "A" & BASE == "G"))
                          %>% group_by(READ_NAME) %>% summarise(target_mut_num = n()))
    tmp_mut_num = left_join(tmp_mut_num, tmp_target_mut_num)
    tmp_mut_num$target_mut_num = ifelse(is.na(tmp_mut_num$target_mut_num), 0, tmp_mut_num$target_mut_num)
    tmp_mut_num$target_mut_ratio = tmp_mut_num$target_mut_num / tmp_mut_num$mut_num
    filtered_reads = tmp_mut_num %>% filter(target_mut_ratio < target_mut_filter_rate)
    tmp_filter_mut_num = test_input %>% filter(!(READ_NAME %in% filtered_reads$READ_NAME))
    
    if(nrow(tmp_filter_mut_num) == 0) {
        return(-1)
    }    
    
    #since all alignments have been converted to plus strand, don't need to keep FLAG=16&A>G any more.
    output_read_name = unique((tmp_filter_mut_num %>% filter((FLAG == 0 & REF == "T" & BASE == "C")|(FLAG == 16 & REF == "A" & BASE == "G")))$READ_NAME)
    mut_rate = length(output_read_name) / length(unique(ori_test$READ_NAME))     
    output_read_name = data.frame("new_read" = output_read_name)

    write.table(output_read_name, file = file.path(output_folder, paste0(target_id, ".txt")), 
              col.names = F, quote = F, row.names = F)
    
    return(mut_rate)
}

newly_synthesised_read <- bplapply(sample_names, function(target_id) {
    function_sample(target_id, input_folder = input_folder, output_folder = output_folder, core = core, SNP = SNP, quality_filter = quality_filter, target_mut_filter_rate = target_mut_filter_rate)
}, BPPARAM = MulticoreParam(workers = core))

newly_synthesised_read = unlist(newly_synthesised_read)
summary_result = data.frame(sample_name = sample_names, mutation_reads_rate = newly_synthesised_read)
write_csv(summary_result, path = file.path(output_folder, "summary.csv"))
summary_result = summary_result %>% filter(mutation_reads_rate >= 0) %>% select(sample_name)
write.table(summary_result, file = file.path(output_folder, "sample_id.txt"), 
              col.names = F, quote = F, row.names = F)
```
