## Supplementary file4 for "*PerturbSci-Kinetics*: Dissecting key regulators of transcriptome kinetics through scalable single-cell RNA profiling of pooled CRISPR screens": seq_align.sh

```
input_folder=$1
sample=$2
output_folder=$3
fa_file=$4

echo generate alignment file: $sample
java_zx="/tools/openjdk_v11/jdk-11.0.2/bin/java"
sam2tsv_zx="/tools/jvarkit/dist/sam2tsv.jar"
$java_zx -jar $sam2tsv_zx -R $fa_file $input_folder/$sample.sam \
> $output_folder/$sample.align
```
