## Supplementary file4 for "*PerturbSci-Kinetics*: Dissecting key regulators of transcriptome kinetics through scalable single-cell RNA profiling of pooled CRISPR screens": UMI_barcode_attach.py

```
"""
Created on Tue Apr  5 22:16:54 2016

@author: Junyue

Modified 03/30/2021 by Andras Sziraki to handle the extraction of barcode from the new sciNEXT protocol.
This script corrects RT and ligation barcodes 1 edit distance away.
"""

import subprocess
import sys
from Levenshtein import distance
import gzip
from multiprocessing import Pool
from functools import partial
import pickle

'''
    this script accept a read1 file, a read2 file, a read3 file, a output_file, a ligation barcode list,
    an RT barcode list,
    and mismatch rate, then it open the read1, read2 and read3, output file,
    then extract the RT barcode and UMI sequence in the read 1 file and the ligation barcode from the read 2 file, and convert the
    barcode to the real barcode in the barcode list based on the mismatch rate (this step does not happen),
    then it attach the barcodes and UMI sequence to the read name of the read3 file
'''    
    
def UMI_attach_read2_barcode_list(sample, input_folder, output_folder, ligation_barcode_list, RT_barcode_list, mismatch_rate = 1):
    #open the read1, read2, and output file
    Read1 = input_folder + "/" + sample + ".R1.fastq.gz"
    Read2 = input_folder + "/" + sample + ".R3.fastq.gz"
    Read3 = input_folder + "/" + sample + ".R2.fastq.gz"
    output_file = output_folder + "/" + sample + ".R2.fastq.gz"
    mismatch_rate = int(mismatch_rate)
    f1 = gzip.open(Read1)
    f2 = gzip.open(Read2)
    f3 = gzip.open(output_file, 'wb')
    f4 = gzip.open(Read3)
    
    line1 = f1.readline()
    line2 = f2.readline()
    line3 = f4.readline()
    total_line = 0
    reads_from_unspecific_targeted_primer = 0
    filtered_line = 0
    
    #generate all potential constant region1 sequences having distance <= 1 with the original sequence.
    #compared with distance calculation, direct sequence matching could be much faster
    ori_constant = "CAAGTTGATA"
    constant_mismatch_list = set()
    for each_pos in range(len(ori_constant)):
        ori_list = list(ori_constant)
        for each_letter in ["A", "G", "C", "T", "N"]:
            ori_list[each_pos] = each_letter
            constant_mismatch_list.add("".join(ori_list))
    constant_mismatch_list = list(constant_mismatch_list)
    
    while (line1):
        total_line += 1
        line1 = f1.readline()
        line3 = f4.readline()
        #print("read1: ", line1)
        # first check if the ligation barcode match with the barcode
        tmp_lig = line3[0:10]
        
        #check if this read is from unspecifc binding of gRNA target primer
        potential_constant_R1 = line1[18:28]
        
        if potential_constant_R1 not in constant_mismatch_list:
        
            if tmp_lig in ligation_barcode_list:

                ligation_bc_match = ligation_barcode_list[tmp_lig]
                # check RT barcode
                target_RT = line1[8:18]
                #print("target_RT: ", target_RT)

                if target_RT in RT_barcode_list:
                    barcode = RT_barcode_list[target_RT]
                    filtered_line += 1
                    UMI = line1[:8]
                    first_line = '@' + ligation_bc_match + barcode + ',' + UMI + ',' + line2[1:]
                    #print("read2: ", first_line)
                    f3.write(first_line)

                    second_line = f2.readline()
                    f3.write(second_line)

                    third_line = f2.readline()
                    f3.write(third_line)

                    four_line = f2.readline()
                    f3.write(four_line)

                    line2 = f2.readline()

                else:
                    line2 = f2.readline()
                    line2 = f2.readline()
                    line2 = f2.readline()
                    line2 = f2.readline()

            else:
                line2 = f2.readline()
                line2 = f2.readline()
                line2 = f2.readline()
                line2 = f2.readline()
        else:
            line2 = f2.readline()
            line2 = f2.readline()
            line2 = f2.readline()
            line2 = f2.readline()
            
            reads_from_unspecific_targeted_primer += 1    

        line1 = f1.readline()
        line1 = f1.readline()
        line1 = f1.readline()

        line3 = f4.readline() 
        line3 = f4.readline()
        line3 = f4.readline()

    f1.close()
    f2.close()
    f3.close()
    f4.close()
    print("sample name: %s, total line: %f, targeted primer unspecific binding line: %f, filtered line: %f, filter rate: %f" 
          %(sample, total_line, reads_from_unspecific_targeted_primer, filtered_line, \
            float(filtered_line) / float(total_line)))

# this function accept an input folder and a output folder and then generate the output file with the index
def attach_UMI_files(input_folder, sampleID, output_folder, ligation_barcode_file, RT_barcode_file, core):
    
    init_message = '''
    --------------------------start attaching UMI-----------------------------
    input folder: %s
    sample ID: %s
    output_folder: %s
    ligation barcode file: %s
    RT barcode file: %s
    ___________________________________________________________________________
    ''' %(input_folder, sampleID, output_folder, ligation_barcode_file, RT_barcode_file)
    
    print(init_message)
    
    print("Load ligation barcode dictionary...")
    
    # generate the ligation barcode list
    barcodes = open(ligation_barcode_file, "rb")
    ligation_barcode_list = pickle.load(barcodes)
    barcodes.close()
    
    print("Load RT barcode dictionary...")
    
    # generate the RT barcode list:
    barcodes = open(RT_barcode_file, "rb")
    RT_barcode_list = pickle.load(barcodes)
    barcodes.close()
    
    #for each sample in the sample list, use the read1 file, read2 file, output file
    # and barcode_list to run UMI_attach_read2_barcode_list
    sample_file = open(sampleID)
    sample_list = []
    for line in sample_file:
        sample = line.strip()
        sample_list.append(sample)
    sample_file.close()
    
    # parallele for the functions
    p = Pool(processes = int(core))
    #print("Processing core number: ", core_number)
    func = partial(UMI_attach_read2_barcode_list, input_folder = input_folder, output_folder=output_folder, ligation_barcode_list = ligation_barcode_list, RT_barcode_list=RT_barcode_list, mismatch_rate = 1)
    #sciRNAseq_count(sample, input_folder, exons, genes, gene_end)
    result = p.map(func, sample_list)
    p.close()
    p.join()
    
    #print the completion message
    com_message = '''~~~~~~~~~~~~~~~UMI attachment done~~~~~~~~~~~~~~~~~~'''
    print(com_message)
    
if __name__ == "__main__":
    input_folder = sys.argv[1]
    sampleID = sys.argv[2]
    output_folder = sys.argv[3]
    ligation_barcode_file = sys.argv[4]
    RT_barcode_file = sys.argv[5]
    core=sys.argv[6]
    attach_UMI_files(input_folder, sampleID, output_folder, ligation_barcode_file, RT_barcode_file, core)
```
